# Molecular phylogenetics of the superfamily Stromboidea (Caenogastropoda): new insights from increased taxon sampling

**DOI:** 10.1101/2024.05.23.595472

**Authors:** Alison R. Irwin, Philippe Bouchet, J. Alistair Crame, Elizabeth M. Harper, Gijs C. Kronenberg, Ellen E. Strong, Suzanne T. Williams

## Abstract

The superfamily Stromboidea is a clade of morphologically distinctive gastropods which include the iconic Strombidae, or ‘true conchs’. In this study, we present the most taxonomically extensive phylogeny of the superfamily to date, using fossil calibrations to produce a chronogram and extant geographical distributions to reconstruct ancestral ranges. From these results, we confirm the monophyly of all stromboidean families; however, six genera are not monophyletic using current generic assignments (Strombidae: *Lentigo, Canarium*, *Dolomena*, *Doxander*; Xenophoridae: *Onustus, Xenophora*). Within Strombidae, analyses resolve an Indo-West Pacific (IWP) clade sister to an East Pacific/Atlantic clade, together sister to a second, larger IWP clade. Our results also indicate two pulses of strombid diversification within the Miocene, and a Tethyan/IWP origin for Strombidae – both supported by the fossil record. However, conflicts between divergence time estimates and the fossil record warrant further exploration. Species delimitation analyses using the COI barcoding gene support several taxonomic changes. We synonymise *Euprotomus aurora* with *Euprotomus bulla*, *Strombus alatus* with *Strombus pugilis*, *Dolomena abbotti* with *Dolomena labiosa*, and *Dolomena operosa* with *Dolomena vittata*. We identified cryptic species complexes within *Terebellum terebellum*, *Lambis lambis*, *“Canarium” wilsonorum, Dolomena turturella* and *Maculastrombus mutabilis*. We reinstate *Rimellopsis laurenti* as a species (previously synonymised with *R. laurenti*) and recognise *Harpago chiragra rugosus* and *Lambis truncata sowerbyi* valid at the rank of species. Finally, we establish several new combinations, rendering *Lentigo*, *Dolomena*, and *Canarium* monophyletic: *Lentigo thersites*, *Dolomena robusta*, *Dolomena epidromis*, *Dolomena turturella*, *Dolomena taeniata, Dolomena vanikorensis*, *D. vittata*, *“Canarium” wilsonorum*, *Hawaiistrombus scalariformis*, *Maculastrombus mutabilis*, *Maculastrombus microurceus*.

## 1 Introduction

Members of the gastropod superfamily Stromboidea Rafinesque, 1815 inhabit predominantly shallow tropical and subtropical marine habitats worldwide, with some ranging into deeper and/or temperate waters. Species are characterised by varied and often elaborate shell morphologies (Savazzi, 1991), and some are well known for their large, colourful eyes (Fig. 1). Owing to these distinctive features, the Stromboidea, and especially the Strombidae Rafinesque, 1815, have been the subject of numerous comparative morphological studies, with discussion on generic relationships and biogeographic patterns (e.g. Simone, 2005; Latiolais et al., 2006; Bandel, 2007).

**Fig. 1.**
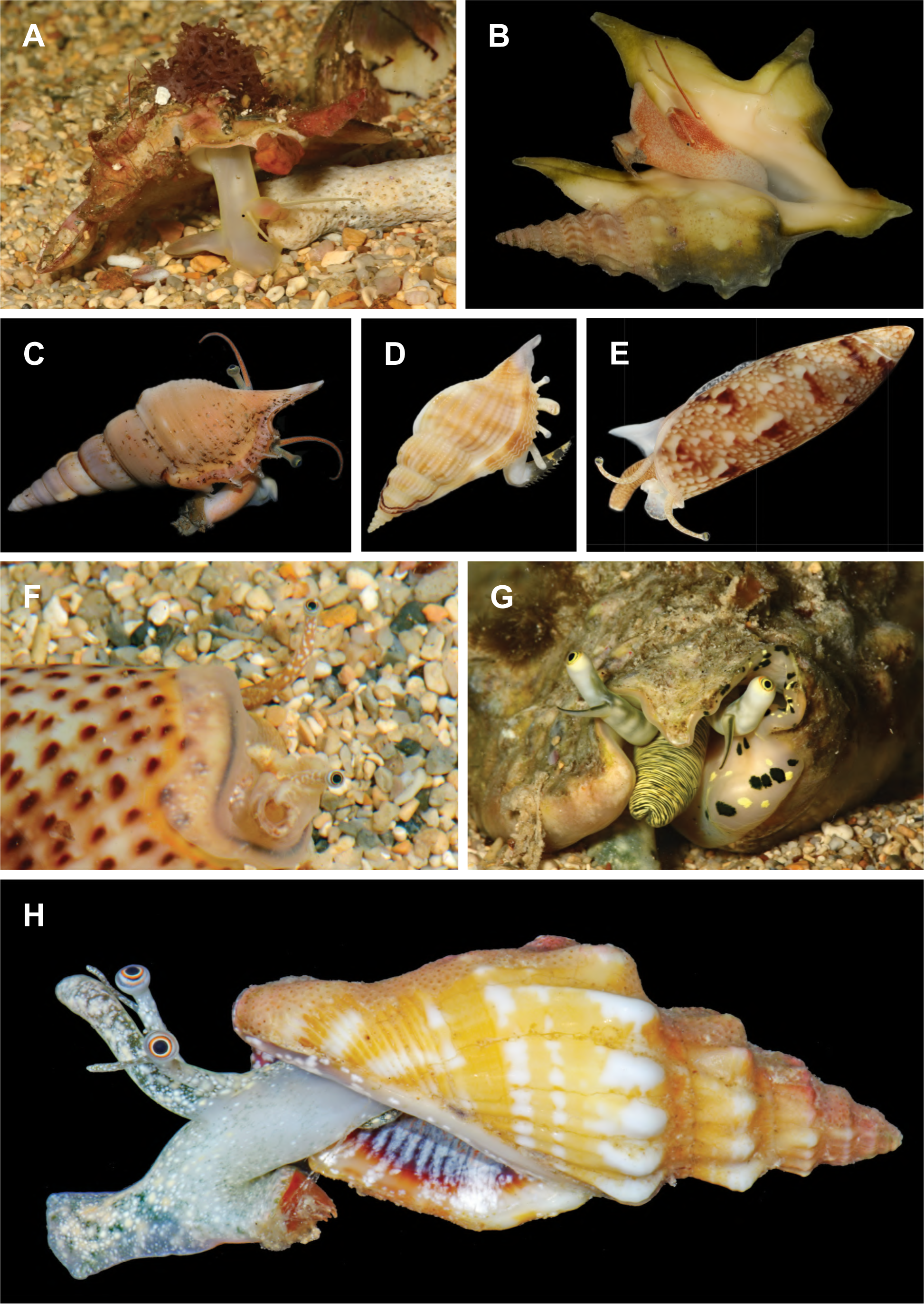
A-H. Live stromboid specimens: (A) xenophorid *Xenophora solarioides*, Île Sainte-Marie, New Caledonia; (B) aporrhaid *Aporrhais pespelecani*, MNHN-IM-2019-17225, Corsica; (C) rostellariid *Rimellopsis powisii,* MNHN-IM-2007-34492, Philippines; (D) rostellariid *Varicospira cancellata,* MNHN-IM-2013-47468, Papua New Guinea; (E) seraphsid *Terebellum terebellum* D, MNHN-IM-2013-11701, Papua New Guinea; (F) seraphsid *T. terebellum*, Île Néaé, New Caledonia; (G) strombid *Lentigo lentiginosus*, Îlot Tenia, New Caledonia; (H) strombid *Canarium elegans,* MNHN-IM-2013-81518, New Caledonia. Images A, F and G taken by David Massemin; remaining images taken by MNHN. Specimens C-E were sequenced in this study (Suppl. Mat. 2); see MNHN database for full collection data for specimens B and H.

Despite this substantial interest in the group, a robust, taxonomically complete phylogeny for Stromboidea is lacking. Recent phylogenetic studies focus primarily on Strombidae, whereas taxon sampling of the other five stromboid families is limited (Latiolais et al., 2006; Irwin et al., 2021; Machkour-M’Rabet et al., 2021; Li et al., 2022a). This has prevented analysis of generic relationships, or confirmation of reciprocal monophyly among families (except for Strombidae and Xenophoridae Troschel, 1852; Irwin et al., 2021). Analyses within the most recent and taxon rich phylogenetic study to date did not support the monophyly of several strombid genera (*Canarium*, *Dolomena*, and *Lentigo*), revealing the need for a systematic revision of group (Irwin et al., in review).

Studies with more complete taxonomic representation are also useful tools in investigating patterns of biogeography and diversification. When calibrated with the fossil record, phylogenetic analyses allow us to estimate timings of key radiation and divergence events and generate hypotheses on the ecological and geographical factors underpinning these events. Tectonic events and concomitant changes to habitats, coastlines, sea level, temperature, and ocean circulation (Wilson & Rosen, 1998; Williams & Duda, 2008) have had well-documented impacts on several other gastropod groups (e.g., Kohn, 1990; Vermeij, 1996; Meyer, 2003; Williams, 2007; Reid et al., 2010). The rich stromboidean fossil record makes this group an excellent subject for time-calibrated phylogenetic studies, with earliest records dating from the Early Jurassic (Aporrhaidae Gray, 1850; Gründel et al., 2009, but see Tracey et al., 1993).

In this study, we reconstruct the most taxon rich phylogeny of the Stromboidea to date, with a focus on Strombidae. We include sequence data from 53% of the currently recognised stromboidean species diversity (59% of strombid species), doubling the taxon sampling in Irwin et al. (in review). This dataset comprises at least one species from all currently accepted strombid genera (except *Mirabilistrombus* and *Amabiliplicatus*); from these, type species are only missing for *Canarium, Hawaiistrombus, Laevistrombus, Titanostrombus* and *Neodilatilabrum* (Suppl. Mat. 1). We use this framework to inform the systematics of the group and use species delimitation methods to infer primary species hypotheses. We also identify stromboid fossils suitable for calibrating a phylogeny, to estimate the timing of major diversification events and geographic divergences.

## 2 Methods

### 2.1 Sample choice

Sampling included at least one representative from 87% of currently recognised stromboidean genera (MolluscaBase, 2023) (Suppl. Mat. 1), and 93 recognised species, representing 381 newly sequenced specimens; see Suppl. Mat. 2 for specimen data. Other sequences were sourced from GenBank; those lacking voucher data (Suppl. Mat. 2) were included if one or more other specimen of that species (according to initial delimitation analyses) with voucher data was available. Exceptions were made for MN703099 *Stellaria solaris*, MK327366 *Onustus exutus* and DQ525214 *Canarium wilsonorum* due to interesting initial results and their importance to systematic discussions. Identifications of genetically unique specimens lacking voucher data are tentative but were consistent with phylogenetic placement in trees in preliminary analyses. Outgroups were two littorinid species chosen based on a previous study (Irwin et al., 2021), with sequences from multiple specimens (Suppl. Mat. 2).

### 2.2 DNA extractions, polymerase chain reactions and Sanger sequencing

DNA was extracted from ethanol-preserved tissue following Irwin et al. (2021). Fragments of nuclear 28S rRNA gene (28S) and three mitochondrial genes (cytochrome oxidase subunit I, COI; 16S rRNA, 16S; and 12S rRNA, 12S), were amplified and sequenced following Williams and Ozawa (2006), except most 28S sequences were obtained with a forward primer designed for Stromboidea (Irwin et al., 2021). Sequencing was undertaken at NHMUK or MNHN.

### 2.3 Sequence analysis

Sequences were assembled and edited using Sequencher v. 5.4.6 (GeneCodes, Ann Arbor, MI). Several datasets were produced: (1) datasets for single gene phylogenetic analyses comprised 199 12S sequences, 208 16S sequences, and 157 28S sequences (including outgroups; Suppl. Mat. 2); (2) datasets for species delimitation analyses comprised 452 and 454 COI sequences (excluding and including the outgroups, respectively); (3) an ingroup-only dataset for the *BEAST analysis comprised all sequence data (472 specimens); (4) an ingroup-only dataset for the fossil-calibrated BEAST analysis comprised 100 terminals (one representative for each putative species delimited in this study). The BEAST dataset included a chimeric sequence for *Varicospira crispata* comprising sequences from different specimens, and three species lacking COI data: *Dolomena japonica*; *Dolomena campbellii*; *Dolomena wienekei* (Suppl. Mat. 2).

Ribosomal RNA genes were aligned in an iterative process via PASTA (Mirarab et al., 2014), using MAFFT L-INS-I (Katoh et al., 2009) to align, OPAL (Wheeler & Kececioglu, 2007) to merge adjacent subset alignments pairs, FASTTREE (Price et al., 2009) to estimate a maximum likelihood tree, GTR+CAT as the nucleotide substitution model, 50% subproblem and centroid decomposition with 5 iterations and the best alignment determined by likelihood value, with minor adjustments to alignments made by eye. Ambiguously aligned regions were excluded via Gblocks 0.91b (Castresana, 2000), with smaller final blocks, gap positions within final blocks and less strict flanking positions allowed; alignment of COI was unambiguous. Putative editing errors were trimmed from GenBank sequence ends (e.g., COI frameshifting indels). The best nucleotide substitution models were chosen via ModelTest-NG (Darriba et al., 2020) using the Akaike Information Criterion: COI, 28S, GTR+I+G; 16S, HKY+I+G; 12S, TrN93+I+G. Variable and phylogenetically informative sites were identified via IQ-TREE (Minh et al., 2013).

### 2.4 Species delimitation

The results of different species delimitation methods for identifying evolutionarily significant units (ESUs) using COI were compared: (1) single threshold general mixed Yule-coalescent model (GMYC; Fujisawa and Barraclough, 2013); (2) single-rate (bPTP; Zhang et al., 2013) and (3) multi-rate Bayesian Poisson Tree Processes (mPTP; Kapli et al., 2017); (4); Automatic Barcode Gap Discovery (ABGD; Puillandre et al., 2012); (5) Assemble Species by Automatic Partitioning (ASAP; Puillandre et al., 2021). These use different criteria to delineate species boundaries (phenetic criteria, ASAP and ABGD; phylogenetic criteria, GMYC, bPTP and mPTP), and vary in performance in other groups, including gastropods (e.g., Aksenova et al., 2018; Strong & Whelan, 2019; Goulding et al., 2023). ESUs were considered putative species (i.e., primary species hypotheses; Puillandre et al., 2012) in *BEAST and BEAST analyses if: (1) ESUs formed a well-supported clade in both BEAST and IQ-TREE COI analyses; (2) ESUs were delimited by at least one method of both phenetic (ABGD and ASAP) and phylogenetic (GMYC, bPTP and mPTP) criteria each. Average interspecific and intraspecific genetic distances for COI were calculated via the Kimura two-parameter model for species with changes proposed to circumscription or rank (K2P; Kimura, 1980) via MEGA v.10.2.6 (Stecher et al., 2020).

ASAP and ABGD analyses were performed with three substitution models (JC, K2P and simple p-distances) via the online servers (ASAP, https://bioinfo.mnhn.fr/abi/public/asap; ABGD, https://bioinfo.mnhn.fr/abi/public/abgd) with default parameters. Only initial partitions were considered in the ABGD analysis (Puillandre et al., 2012). An ultrametric COI tree for the GMYC analysis was produced using Bayesian Inference (BI; Huelsenbeck et al., 2001) via BEAST v.1.10.4 (Drummond & Rambaut, 2007) without fossil calibrations and with an uncorrelated relaxed lognormal clock, a constant population size coalescent prior (more conservative than a Yule prior; Monaghan et al., 2009), and a UPGMA starting tree. The analysis ran for 150,000,000 generations (sampling frequency, 5,000). Convergence was checked via Tracer v.1.7.1 (Rambaut et al., 2018); ESS values >200 indicated adequate sampling. The final tree was produced in TreeAnnotator based on 27,000 trees with maximum clade credibility (MCC) and median node heights, and posterior probabilities (PP) calculated by BEAST. We used SPLITS (Ezard et al., 2009) in R v.4.1.3 (R Core Team, 2022) to identify species boundaries. Maximum likelihood (ML; Felsenstein, 1981) trees were produced in IQ-TREE for PTP analyses using the ultrafast bootstrap (UFBoot) feature (Hoang et al., 2018). The mPTP and bPTP models were run via mPTP v.0.2.4 (https://github.com/Pas-Kapli/mptp). Confidence of delimitations was assessed via ten MCMC chains (generations, 100,000,000; sampling frequency, 5,000) with 10% burnin; convergence of independent runs was verified with Average Standard Deviation of Delimitation Support Values (ASDDSV; Kapli et al., 2017).

### 2.5 Phylogenetic reconstructions

Single-gene BI analyses for 12S, 16S and 28S were produced via MrBayes (v.3.2.2; Huelsenbeck and Ronquist, 2001). Four MCMC chains were run for 100,000,000 generations (sample frequency, 1,000). We examined .p files for stationarity and convergence in Tracer; consensus trees were obtained after a 10% burnin. A starting tree for the concatenated BEAST analysis was produced via *BEAST, v.2.6.6 (Bouckaert et al., 2019); the phylogenetic inference of this method is suggested to be superior (Heled and Drummond, 2010). The *BEAST analysis ran for 900,000,000 generations (sample frequency, 5,000) without fossil calibrations (due to computational limitations) and with a Yule tree prior for species-level analyses, a constant coalescent model for population-level analyses, a lognormal clock, and a dataset partitioned by gene fragment. The final species tree was an MCC tree with median node heights and 10% burnin. The time-calibrated phylogeny was produced in BEAST with a dataset partitioned by gene fragment, each gene allowed to evolve at a different rate, a Yule speciation prior, the *BEAST tree as the starting topology, and an uncorrelated relaxed, lognormal clock with four fossil calibrations (section 2.6). We ran the analysis for 180,000,000 generations (sampling frequency, 3,000), and examined log files for convergence. The final species tree was a MCC tree with median node heights, 10% burnin, and PP node support.

### 2.6 Fossil calibrations

Based on the earliest undisputed fossil representatives from each family (including extinct subfamilies), preliminary BEAST and *BEAST analyses, and previous mitogenome work (Irwin et al., 2021), four fossil calibrations were used in the BEAST analysis (Table 1; for traits used to identify fossils to family level, see Suppl. Mat. 3). Initial *BEAST analyses did not recover a monophyletic Rostellariidae Gabb, 1868; therefore, a rostellariid fossil was used to calibrate the crown age of Rostellariidae + Seraphsidae Gray, 1853 (Table 1). Initial BEAST analyses with separate calibrations for Aporrhaidae and Struthiolariidae Gabb, 1868 had highly unrealistic node ages; therefore, an aporrhaid fossil was used to calibrate the crown age of Aporrhaidae + Stuthiolariidae (Table 1). A prior study by the authors used the same calibrations without justification (Irwin et al., in review); a justification of these choices is given below.

**Table 1.** Fossil calibrations used in BEAST analyses, with age of earliest fossil representative and corresponding geologic age from Gradstein et al. (2020) and calibration parameters. For complete details and justification of fossil representatives used, see sections 2.6.2–5.

| Name | Age of earliest representative | Geologic age | Mean in real space | Log stdev | Offset | 95% interval |
| --- | --- | --- | --- | --- | --- | --- |
| **Xenophoridae** | 100.5 – 93.9 | Cenomanian (Upper Cretaceous) | 9 | 8 | 93.9 | 95.2 – 117.5 |
| **Aporrhaidae + Struthiolariidae** | 199.3 – 190.8 | Sinemurian (Lower Jurassic) | 12 | 10 | 190.8 | 193.6 – 221.2 |
| **Rostellariidae + Seraphsidae** | 83.6 – 66.0 | Campanian/Maastrichtian (Upper Cretaceous) | 9 | 7 | 66.0 | 67.6 – 91.0 |
| **Strombidae** | 41.2 – 47.8 | Lutetian (Middle Eocene) | 5 | 5 | 41.2 | 42.1 – 59.3 |

#### 2.6.2 Aporrhaidae + Struthiolariidae

A specimen of *Toarctocera subpunctata* (Münster, 1844) from the Late Toarcian (Baden Württemberg) is the oldest complete aporrhaid specimen recorded in the literature (Gründel et al., 2009, fig. 2). However, numerous older, incomplete specimens (e.g., Terquem, 1855, pl. 17, fig. 5; Moore, 1867, pl. 14, figs 23–25; Piette, 1876, pl. 1, figs 2, 3; Haas, 1953, pl. 16, figs 51, 55; Schubert et al. 2008, fig. 5c) suggest an earlier origin for Aporrhaidae. A specimen of *Alaria hudlestoni* E. Wilson, 1887 from the lower Sinemurian (Lower Lias, Gloucestershire, Wilson, 1887, pl. 5, fig. 13, dated by Gründel et al., 2009) is a damaged shell, but has more intact diagnostic characters than earlier representatives (Tracey et al., 1993), and was used to date the crown age of Aporrhaidae + Struthiolariidae (Table 1).

**Fig. 2.**
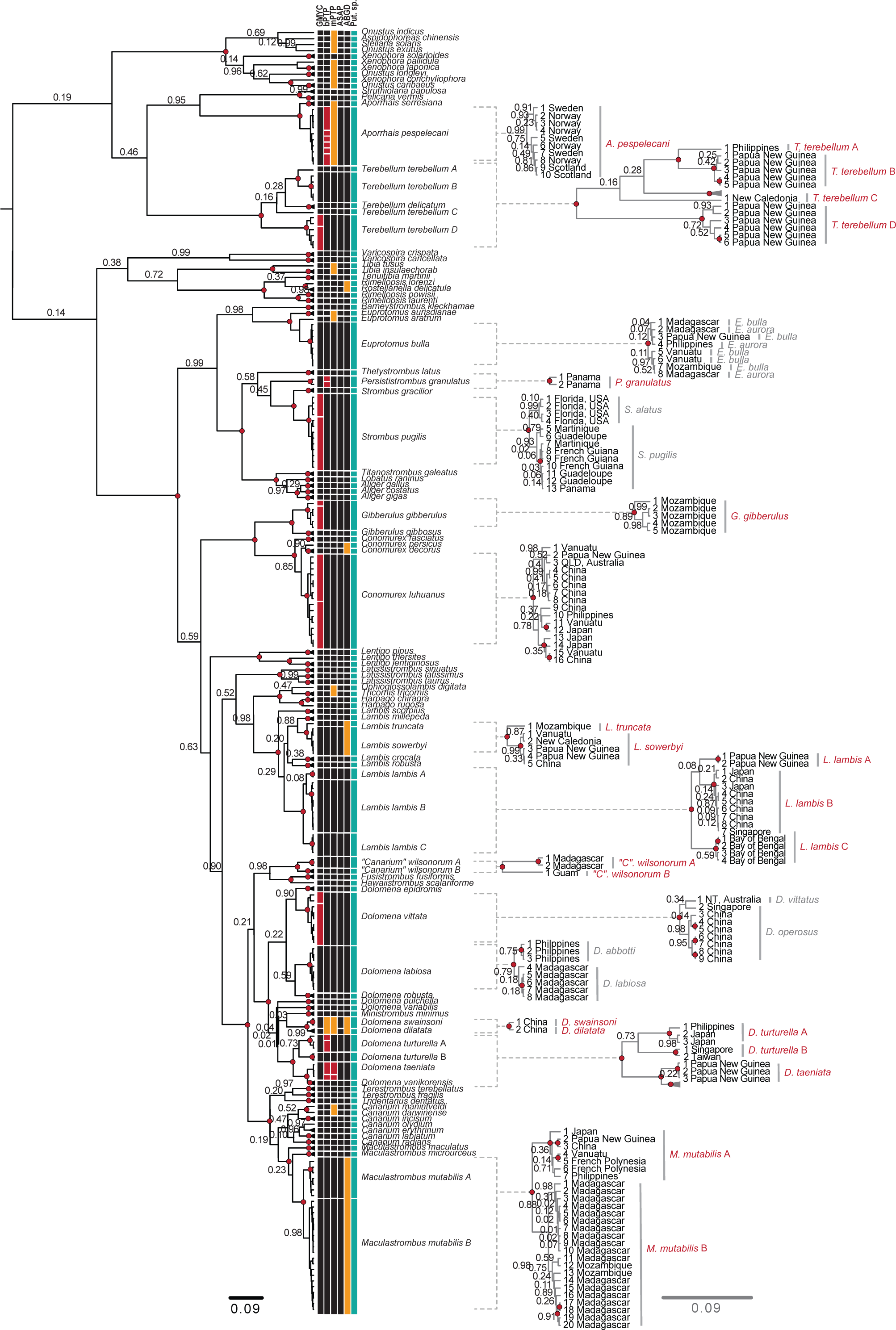
Bayesian analysis of COI sequences via BEAST. ESUs delimited by five methods (section 3.2) are depicted via blocks (black, accepted ESUs; red, unaccepted splits; orange, unaccepted lumps), with putative species (green) on the right with names listed (black font). Unless any delimitation method proposed new splits or accepted lumps, species with multiple specimens were collapsed at tips (indicated by triangles; see Suppl. Mat. 5 for complete tree with all tips annotated), and relevant topologies were enlarged to the right (grey branches), with specimen number and country of collection given at tips, and all support values on branches (Suppl. Mat. 2). If primary species hypotheses lumped taxa, nominal species are listed in grey to the right; other species names are in red. Support values are posterior probabilities (PP); intraspecific PP were removed for legibility in the full tree (see Suppl. Mat. 5 for all support values).

#### 2.6.3 Xenophoridae

The earliest Xenophoridae is generally dated to the Late Cretaceous (Ponder, 1983). Early specimens are often internal casts, with a trochiform shell shape and scars, or impressions, from the agglutination of foreign objects as the only useful characters for identification to family level (Wade, 1926; Stephenson, 1952 (Fig. 1A; Suppl. Mat. 3). These scars may be confused with taphonomic artefacts. The earliest xenophorid specimen (Ponder, 1983; Tracey et al., 1993; Kiel and Perrilliat, 2001) is a poorly preserved cast of *Xenophora*? sp. with impressions possibly from agglutinated shell fragments (Stephenson, 1952 (USNM PAL 105677, not figured), from the Cenomanian (Woodbine Formation, Texas). A cast and an incomplete shell of *Xenophora leprosa* (Morton, 1834) from the Campanian/Maastrichtian have more convincing scars (Ripley Formation, Tennessee: Wade, 1926, pl. 56, figs 7–8; Corsicana Marl, Texas: Stephenson, 1941, figs 17–19). *Acanthoxenophora* Perilliat & Vega 2001 is recorded from the Campanian and possibly the Santonian (Krüger, 2006, fig. 11; Pietzonka et al., 2023, fig. 5). However, preliminary analyses with these calibrations led to highly unrealistic node ages; therefore, we use *Xenophora*? sp. (Cenomanian) to date the crown age (Table 1).

#### 2.6.4 Rostellariidae + Seraphsidae

Hypotheses on the earliest rostellariid range from the Cenomanian-Turonian to the Maastrichtian (Roy, 1996; Bandel, 2007). The oldest specimen according to Tracey et al. (1993) is ?*Dientomochilus stueri* from the Coniacian (Condat, France: Cossmann, 1904, pl. 9, figs 5–6); however, this is an incomplete internal mould. *Eucalyptraphorus palliatus* (Forbes, 1846) is reported from the Turonian-Coniacian and Maastrichtian-Campanian (Trichinopoly and Ariyalur Groups, India: Stoliczka, 1868, pl. 2, figs 18–20; Acharyya and Lahiri, 1991); the former is a mould, and the latter an incomplete shell. A *Calyptraphorus itamaracensis* specimen (Gramame Formation, Pernambuco, Brazil: Muñiz, 1993, pl. 11, figs 5, 8), reported from the Campanian, was considered the oldest representative by Perilliat and Vega (1997); however, this formation has since been identified as Maastrichtian (e.g. El Gadi and Brookfield, 1999). Therefore, we use *E. palliatus* (Campanian/Maastrichtian) to date the crown age of Rostellariidae + Seraphsidae (Table 1).

#### 2.6.5 Strombidae

The earliest strombid record is *Stromboconus suessi* (Bayan 1870, p. 480) from the Middle Eocene of Roncà, Italy. This specimen is not figured; however, other fossils of *S. suessi* are recorded from the Upper Lutetian of the same locality (Wieneke et al., 2023), which we used to date the crown age of the strombid clade (Table 1).

### 2.7 Diversification rate and ancestral range reconstructions

A relative cladogenesis (RC) test, conducted via R package Geiger v.2.0.7 (Pennell et al., 2014), was used to detect significant increases in diversification rate in the time-calibrated tree. The gamma (γ) statistic (Pybus and Harvey, 2000) was used to detect differences in diversification rate using ‘lineages through time’ (LTT) plots, for (1) stromboidean and (2) strombid lineages. Ancestral ranges were estimated via R package BioGeoBEARS v.1.1.2 (Matzke, 2013), using the time-calibrated tree and a maximum range size of four. Geographic ranges were based on literature records (see Suppl. Mat. 3). For putative cryptic species, only localities for specimens with molecular data available were included. Eight marine biogeographical regions were from Spalding et al. (2007), modified based on stromboidean ranges: 1, Temperate North-East Atlantic; 2, Western Indo-West Pacific (IWP); 3, Central IWP; 4, Eastern IWP; 5, Temperate New Zealand and Australia; 6, Tropical Eastern Pacific; 7, Tropical West Atlantic; 8, Tropical East Atlantic. The best model according to AIC and AICc was BAYAREA-like + j (d = 6.3 x 10^-5^; e = 5.6 x 10^-3^; j = 0.014; LnL = −224.0) (Landis et al., 2013; Matzke, 2014).

## 3 Results

### 3.1 Sequence analysis

COI alignments of 658 bp included 356 variable and 266 phylogenetically informative. For the BEAST dataset, 530 bp of 12S sequence (83% of 640 bp), 496 bp of 16S (90% of 554 bp), and 1,347 bp of 28S (83% of 1,630 bp) remained after removal of ambiguous blocks. Variable and phylogenetically informative sites, respectively, were as follows: 288 and 271 for 12S; 199 and 172 for 16S; 350 and 239 for 28S. For MrBayes and *BEAST datasets, total sequence length and variable and phylogenetically informative sites, respectively, were as follows: 520 bp, 288 and 272 sites for 12S; 496 bp, 212 and 178 sites for 16S; 1,353 bp, 378 and 253 sites for 28S.

### 3.2 Species delimitation analyses

COI species delimitation analyses suggested 97 putative stromboid species (not including the three species lacking COI data; Suppl. Mat. 2, 4). Most of the species that differed from current species concepts have sister taxa that are sympatric or co-occur in at least one of the same ecoregions (as defined in section 2.8) (Table 2). Estimated divergence times between cryptic sister species/clades were ≥ 5 Myr, with varying K2P distances (3.7–12.3%); intraspecific K2P distances ranged from <0.1–1.7% (Table 2). Here, cryptic species are defined as those in conflict with current taxonomy. All putative species were delimited by ASAP across all distance metrics (JC, ASAP-score 3.5, p = 0.018, W = 2.6 x 10^-4^, T_d_ = 0.029; K2P, ASAP-score 4.0, p = 0.018, W = 2.6 x 10^-4^, T_d_ = 0.029; p-distance, ASAP-score 3.5, p = 0.020, W = 1.7 x 10^-4^, T_d_ = 0.028) (Fig. 2; Suppl. Mat. 4, 5). ABGD recovered 92 ESUs using the JC model (p = 0.04); p-values for other models were not significant. Support for the GMYC model was significant (likelihood ratio test (LR): GMYC, 3932; null model (null), 3860; ratio: 145, p <0.001); the threshold time (−0.012) resulted in 79 clusters (95% CI, 78–82) and 102 ESUs (including species represented by a single specimen; 95% CI, 99–105) (Fig. 2; Suppl. Mat. 4, 5). Support for the bPTP model was significant (LR: bPTP, 1905; null, 1495; p <0.001). MCMC chains converged in PTP analyses (both ASDDSV <0.001; mPTP, 86 ESUs; bPTP, 104 ESUs) (Fig. 2; Suppl. Mat. 4, 5).

**Table 2.** Summary of delimitation analyses for taxa with changes to rank or circumscription (Suppl. Mat. 2). Species numbered (n) in order of clades in Fig. 2, with species name, geographical range (based on genetic sequences only), sister clades/species (based on relationships in the BEAST tree; Fig. 3), whether sister clades/species co-occur in at least one of the eight biogeographical regions in Fig. 3 (see section 2.8), divergence time between sister clades (nearest Myr; from Fig. 3), the occurrence of fixed differences between sister clades/species for sequence from 28S (Y, yes; N, no; NA, no 28S data for this species and/or sister clade or species), average K2P distance based on COI sequence data within each ESU, and between the ESU listed and the sister clade/species separated by the smallest genetic distance using COI sequence only (NA, no COI data for this species and/or sister clade or species), reciprocal monophyly (rec. mon.) of clades/species in MrBayes (12S, 16S, 28S; Suppl. Mat. 8) or BEAST (COI; Fig. 2; Suppl. Mat. 5) gene trees (Y, yes; N, no; NA, no sequence data).

| 1. Cryptic species complexes |  |  |  |  |  |  |  |  |  |  |  |  |
| --- | --- | --- | --- | --- | --- | --- | --- | --- | --- | --- | --- | --- |
| N. | Species | Geographical range | Sister species/clade | Co-occurrence in ecoregion | Divergence time (Myr) | Fixed diffs 28S | % K2P inter COI | % K2P intra COI | COI rec. mon. | 12S rec. mon. | 16S rec. mon. | 28S rec. mon. |
| **15** | *Terebellum terebellum* A | Philippines | *T. terebellum* B | Y | 27 | NA | 7.8 | NA | NA | NA | NA | NA |
| **16** | *Terebellum terebellum* B | Papua New Guinea | *T. terebellum* A | Y | 27 | NA | 7.8 | 0.6 | Y | Y | Y | Y |
| **18** | *Terebellum terebellum* C | New Caledonia | *T. delicatum* | Y | 34 | Y | 12.1 | NA | NA | NA | NA | NA |
| **19** | *Terebellum terebellum* D | Papua New Guinea | *T. terebellum* C<br>+ *T. delicatum* | Y | 39 | Y | 12.1 | 2.3 | Y | Y | Y | Y |
| **64** | *Lambis lambis* A | Papua New Guinea | *L. lambis* B + *L. lambis* C | Y | 8 | Y | 5.2 | <0.1 | Y | Y | Y | N |
| **65** | *Lambis lambis* B | Japan, China, Singapore | *L. lambis* C | N | 5 | NA | 4.7 | 0.3 | Y | Y | Y | N |
| **66** | *Lambis lambis* C | Bay of Bengal | *L. lambis* B | N | 5 | NA | 4.7 | 0.3 | Y | NA | NA | NA |
| **67** | *"Canarium" wilsonorum* A | Madagascar | *"C". wilsonorum* B | N | 21 | NA | 9.6 | 0.9 | Y | NA | NA | NA |
| **68** | *"Canarium" wilsonorum* B | Guam | *"C". wilsonorum* A | N | 21 | NA | 9.6 | NA | NA | NA | NA | NA |
| **80** | *Dolomena turturella* A | Philippines, Japan | *Dolomena turturella* B | Y | 19 | N | 8.2 | 0.3 | Y | Y | Y | N |
| **81** | *Dolomena turturella* B | Singapore, Taiwan | *Dolomena turturella* A | Y | 19 | N | 8.2 | 0.6 | Y | Y | Y | N |
| **96** | *Maculastrombus mutabilis* A | Papua New Guinea, Vanuatu, Philippines, Japan, China, French Polynesia | *M. mutabilis* B | N | 7 | N | 4.6 | 1.4 | Y | N | N | N |
| 97 | *Maculastrombus mutabilis* B | Madagascar, Mozambique | *M. mutabilis* A | N | 7 | N | 4.6 | 0.9 | Y | Y | Y | N |
| **2. Newly synonymised species** |  |  |  |  |  |  |  |  |  |  |  |  |
| 32 | *Euprotomus bulla* | Madagascar, Mozambique, Papua New Guinea, Philippines, Vanuatu | *E. aratrum* + *E. aurisdianae* | Y | 22 | Y | 5.7 | 0.4 | Y | Y | Y | N |
| 36 | *Strombus pugilis* | Florida (USA), French Guiana, Guadeloupe, Martinique, Panama | *S. gracilior* | Y | 23 | NA | 12.3 | 1.3 | Y | Y | Y | Y |
| 73 | *Dolomena labiosa* | Madagascar, Philippines | *D. wienekei* | Y | 6 | Y | NA | 0.9 | Y | Y | N | N |
| 72 | *Dolomena vittata* | Singapore, China, Australia | *D. campbelli* | Y | 9 | NA | 8 | 1.7 | Y | Y | Y | NA |
| **3. Previously synonymised species** |  |  |  |  |  |  |  |  |  |  |  |  |
| 27 | *Rimellopsis powisii* | Philippines | *R. laurenti* | Y | 14 | Y | 7.7 | 0.3 | Y | Y | Y | N |
| 28 | *Rimellopsis laurenti* | Vanuatu, New Caledonia | *R. powisii* | Y | 14 | Y | 7.7 | 0.4 | Y | Y | Y | N |
| **4. Changes to taxonomic rank – subspecies raised to species** |  |  |  |  |  |  |  |  |  |  |  |  |
| 56 | *Harpago chiragra* | Papua New Guinea, Japan, China | *H. rugosa* | Y | 17 | N | 9.5 | 0.5 | Y | Y | Y | N |
| 57 | *Harpago rugosa* | New Caledonia | *H. chiragra* | Y | 17 | N | 9.5 | NA | NA | NA | NA | NA |
| 60 | *Lambis truncata* | Mozambique | *L. sowerbyi* | N | 6 | N | 3.7 | NA | NA | NA | NA | NA |
| 61 | *Lambis sowerbyi* | Papua New Guinea, New Caledonia, Vanuatu, China | *L. truncata* | N | 6 | N | 3.7 | 0.3 | Y | N | Y | N |

Delimitation results conflicted with some current species concepts in Strombidae, Seraphsidae and Rostellariidae. For Strombidae, the 69 putative species with multiple COI sequences received high to maximal support in all COI trees (PP = 0.99–100; BS = 94–100) (Fig. 2; Suppl. Mat. 4–6). All methods recognised *Dolomena abbotti* and *D. labiosa* as one species (*D. labiosa*) with maximal support, as well as *Euprotomus aurora* and *E. bulla* (now *E. bulla*). All methods except GMYC recognised *Strombus pugilis* and *S. alatus* as one species (*S. pugilis*) with maximal ASV support for bPTP and mPTP, indicating that data supported the ML solution; low support for GMYC suggested either within-species structure or very close sister species (GMYC = 0.65/0.65) (Fig. 2; Suppl. Mat. 4–6). ASAP, ABGD, mPTP and bPTP recognised *Dolomena vittata* (formerly *Doxander*) and *D. operosus* as one species (*D. vittata*) with maximal ASV support for bPTP and mPTP (Fig. 2; Suppl. Mat. 4–6). By contrast, GMYC delimited two ESUs with maximal support: 1) *D. vittata* (Singapore) + *D. operosus*; 2) *D. vittata* (China) (Fig. 2; Suppl. Mat. 4–6). ABGD, mPTP and bPTP recognised *Dolomena swainsoni* and *D. dilatata* as one species with maximal support. No taxonomic changes are suggested for these species, since there is no voucher for *D. swainsoni* and so we cannot rule out the possibility that the specimen we sequenced as ‘*D. swainsoni’* was misidentified (Fig. 2; Suppl. Mat. 4–6). Several cryptic species complexes were identified: *Lambis lambis* was split into three putative species (A-C), and *Maculastrombus mutabilis, Dolomena turturella* and *“Canarium” wilsonorum* were split into two species each (A, B), all with maximal support except *M. mutabilis* B (GMYC = 0.63) and *D. turturella* A (GMYC = 0.97; bPTP = 0.74, mPTP = 0.9). (Fig. 2; Suppl. Mat. 4–6). All methods delimited the following sister taxa (formerly subspecies) as separate species with maximal support: *Lambis truncata* and *L. sowerbyi*, and *Harpago chiragra* and *H. rugosus* (Fig. 2; Suppl. Mat. 4–6). For Rostellariidae, putative species had maximal support in all COI phylogenetic analyses (except *Rostellariella delicatula*; PP = 0.98, BS = 67) and all delimitation methods (Fig. 2; Suppl. Mat. 4–6). Here, *Rimellopsis laurenti* is separate from *R. powisii* (Fig. 2; Suppl. Mat. 4– 6) with which it is often synonymised (Whitehead, 1996). For Seraphsidae, five putative cryptic species of *Terebellum terebellum* (e.g., Fig. 1E) had maximal support in all COI analyses and all delimitation methods except GMYC (GMYC = 0.81/1) (Fig. 2; Suppl. Mat. 4–6).

### 3.3 Phylogenetic analyses

Strombidae was recovered with high to maximal support in all trees (PP = 0.96–1; BS = 100%), as was Seraphsidae (PP = 0.95–1; BS = 100%), Xenophoridae (PP = 0.97–1; BS = 100%), Aporrhaidae and Struthiolariidae (both PP = 1; BS = 100%) (Figs 2–3; Suppl. Mat. 5–8). The BEAST tree recovered Rostellariidae as monophyletic with low support (PP = 0.65), contrary to the *BEAST tree which recovered *Varicospira* as sister to the remaining Rostellariidae + Seraphsidae, with variable support (PP = 0.55–0.84) (Fig 1; Suppl. Mat. 7). Individual gene trees differed in topology; only 12S recovered Rostellariidae (PP = 0.98) (Fig. 2, Suppl. Mat. 5–8).

**Fig. 3.**
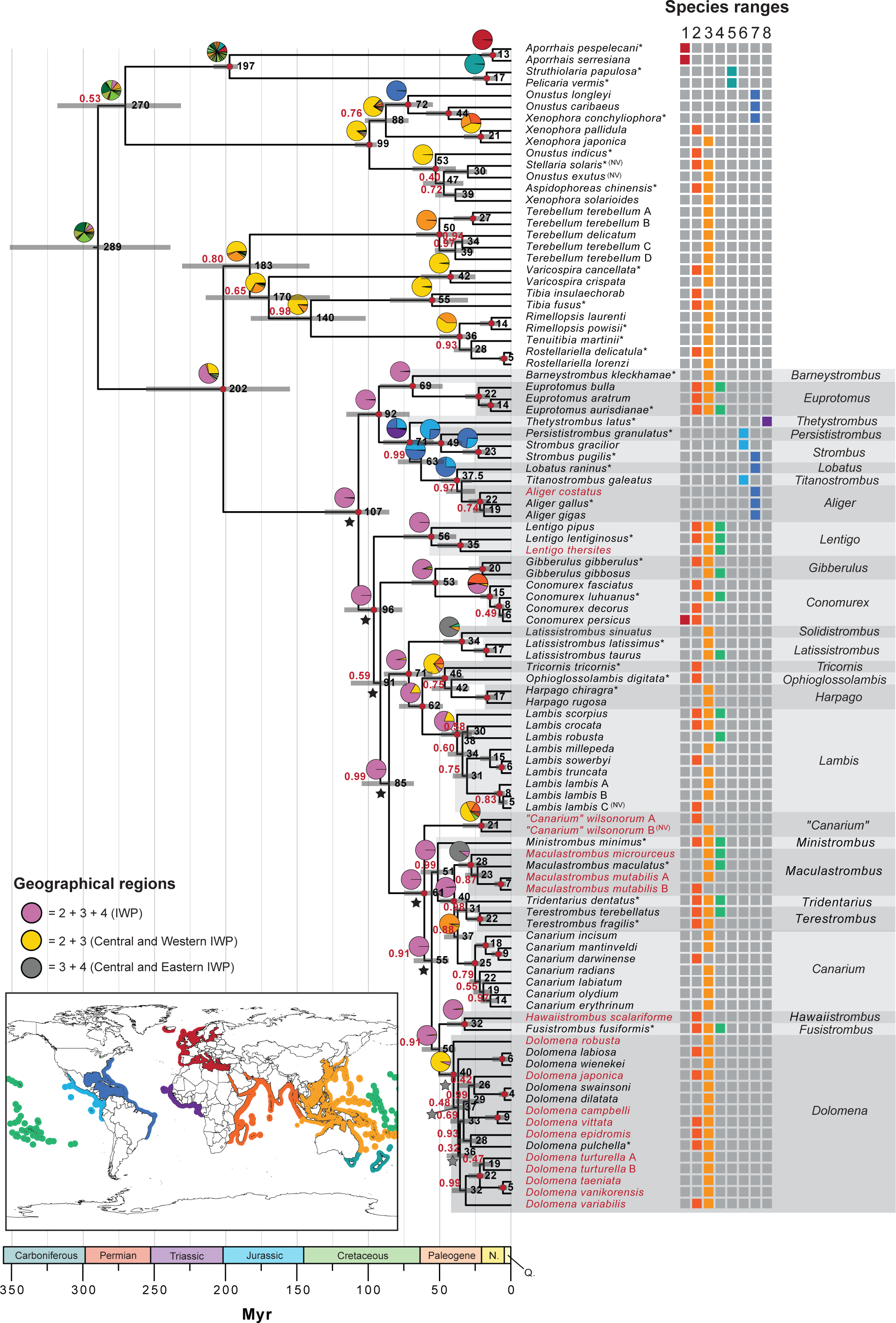
Time-calibrated Bayesian analysis of COI, 12S, 16S, and 28S data as implemented in BEAST. Scale is given below with geologic age. Grey bars are 95% highest posterior density intervals for node ages; mean node ages (black) are rounded to the nearest Myr. Posterior probability (PP) values (red) are on branches (red circles, PP = 1). Pie charts show probabilities of ancestral ranges for major clades, estimated via BioGeoBEARS (regions 1–8; see map and section 2.8), including ranges overlapping regions (1) 2 + 3 + 4, (2) 2 + 3, and (3) 3 + 4 (see key). Stars indicate relative cladogenesis significance levels (black, p <0.05; grey, p <0.01). Asterisks (*) mark putative type taxa (except for *Terebellum terebellum* and *Lambis lambis* where the ‘real’ type is unknown); species with changed generic assignments are in red font. NV, GenBank sequences lacking voucher data. Strombid genera are marked for ease of discussion.

Within Strombidae, several genera were not monophyletic. For *Dolomena,* this was due to the inclusion of *Laevistrombus,* the type species of *Doxander*, and the monospecific *Labiostrombus* with maximum support (Fig. 3). Unlike *Doxander, Laevistrombus* was monophyletic with maximum support in all analyses; however, *Dolomena* has priority, so *Laevistrombus, Labiostrombus* and *Doxander* are no longer considered valid (Fig. 3; Suppl. Mat. 7). Species currently assigned to *Canarium* formed four well supported clades, as follows. (1) *Hawaiistrombus scalariformis* was sister to *Fusistrombus fusiformis*, which together are sister to the clade here assigned to *Dolomena*, as described above, with good support (PP = 0.91–1) (Fig. 3). (2) A clade containing species formerly synonymised with the type species of *Canarium* (*C. urceus,* not included in this study) was recovered as sister to *Tridentarius + Terestrombus* with moderate to maximal support (PP = 0.88–1) (Fig. 3). Together, these were sister to (3) a clade containing the type species of *Maculastrombus,* and several other species currently assigned to *Canarium*, recovered with maximal support (Fig. 3). (4) *“Canarium” wilsonorum* is resolved as sister to all species currently assigned to *Canarium*, plus *Ministrombus, Maculastrombus, Terestrombus, Tridentarius, Dolomena, Labiostrombus, Laevistrombus, Doxander,* and *Fusistrombus* (Fig. 3; Suppl. Mat. 7). All concatenated and COI analyses reject the monophyly of *Lentigo* due to the inclusion of the type species of *Thersistrombus*, resolving *L. pipus + (L. lentiginosus + L. thersites*) with high to maximal support (PP = 0.98–1; BS = 98– 100). As *Lentigo* has priority, *Thersistrombus* is no longer considered valid (Figs 1G, 3; Suppl. Mat. 7. Analyses reject the monophyly of xenophorids *Xenophora* and *Onustus*; however, this is not the focus of this study (Figs 2–3; Suppl. Mat. 5–8).

### 3.4 Taxonomic changes

We hereby recognise the following new combinations (see Suppl. Mat. 2): *Lentigo thersites* (Swainson, 1823) (previously *Thersistrombus*), *Dolomena robusta* (G. B. Sowerby III) (previously *Neodilatilabrum*), *Dolomena epidromis* (Linnaeus, 1758) (previously *Labiostrombus*), *Dolomena vittata* (Linnaeus, 1758) (previously *Doxander* Wenz, 1940), *Dolomena turturella* (Röding, 1798), *Dolomena taeniata* (Quoy & Gaimard, 1834)*, Dolomena vanikorensis* (Quoy & Gaimard, 1834) (all four previously *Laevistrombus*), *“Canarium” wilsonorum* (Abbott, 1967), *Hawaiistrombus scalariformis* (Duclos, 1833), *Maculastrombus mutabilis* (Swainson, 1821), *M. microurceus* Kira, 1959 (all four previously *Canarium*). These changes to generic assignments mean that *Lentigo*, *Dolomena*, and *Canarium*, are now monophyletic. Two taxa hitherto ranked as subspecies are now recognised as distinct species: *Harpago rugosus* (G. B. Sowerby II, 1842) (previously *H. chiragra rugosus*) and *Lambis sowerbyi* (Mörch, 1872) (previously *L. truncata sowerbyi*); also, we reinstate *Rimellopsis laurenti* Duchamps, 1992) as a species (previously synonymised with *Rimellopsis powisii* (Petit de la Saussaye, 1840) (Fig. 1C; Suppl. Mat. 2). We also recognise *Euprotomus aurora* Kronenberg 2002 and *Euprotomus bulla* (Röding, 1798) as one species (*E. bulla*), and similarly, *Strombus pugilis* Linnaeus, 1758 and *Strombus alatus* Gmelin, 1791 (*S. pugilis*), *Dolomena labiosa* (W. Wood, 1828) and *Dolomena abbotti* Dekkers & Liverani, 2011 (*D. labiosa*) and *D. vittata* (Linnaeus, 1758) and *Dolomena operosa* (Röding, 1798) (*D. vittata*) (Suppl. Mat. 2). We synonymise *Eustrombus* Wenz, 1940 with *Aliger* Thiele, 1929, synonymise *Millepes* Mörch, 1852 with *Lambis* Röding, 1798 as is supported by Kronenberg (1993), and confirm the synonymy of *Macrostrombus* Petuch, 1994 with *Aliger sensu* Dekkers (2008). Finally, we confirm the placement of *Ministrombus variabilis* (Swainson, 1820) in *Dolomena* as established by Liverani (2014).

### 3.5 Diversification and dispersal through time

The BEAST chronogram suggests that Struthiolariidae and Aporrhaidae diverged 197 Mya (95% highest probability density interval, HPD: 192–208 Mya), and that Xenophoridae diverged from lineages giving rise to Struthiolariidae and Aporrhaidae 270 Mya (95% HPD: 232–317 Mya), radiating around 99 Mya (95% HPD: 95–109 Mya) (Table 1; Fig. 3). These results suggest Seraphsidae diverged from Rostellariidae 183 Mya (95% HPD: 142–230 Mya) and radiated 50 Mya (95% HPD: 35–66 Mya) (Table 1; Fig. 3). An older radiation was suggested for Strombidae and Rostellariidae than is known from the fossil record: 107 Mya (95% HPD: 86–130 Mya) and 183 Mya (95% HPD: 127–213 Mya), respectively (Table 1; Fig. 3). Significant increases in diversification rate were identified in the strombid lineage (RC, p <0.01–0.05); six occurred at nodes with moderate to maximal support (PP = 0.91–1), and three at younger nodes lacking support (PP <0.50–0.59) (Fig. 3). LTT plots suggested two younger diversification events in Strombidae, at approximately 17–23 Mya and 6–9 Mya (Fig. 4B). Both LTT plots were convex, suggesting decreasing speciation rates through time (Harvey et al., 1994); this was supported for Strombidae by a significant negative γ-value (γ = -3.13, p <0.01), implying an early burst in diversification (Pybus and Harvey, 2000), but not for Stromboidea (γ = -0.06, p = 0.95) (Fig. 4). Ancestral ranges of Stromboidea and the clades containing (1) Xenophoridae + Struthiolariidae + Aporrhaidae and (2) Strombidae + Rostellariidae + Seraphsidae were all unresolved (Fig. 3). The ancestral ranges of Xenophoridae and Strombidae were resolved with moderate and high probability, respectively (Central and Western Pacific, p = 0.87; IWP, p = 0.99) (Fig. 3).

**Fig. 4.**
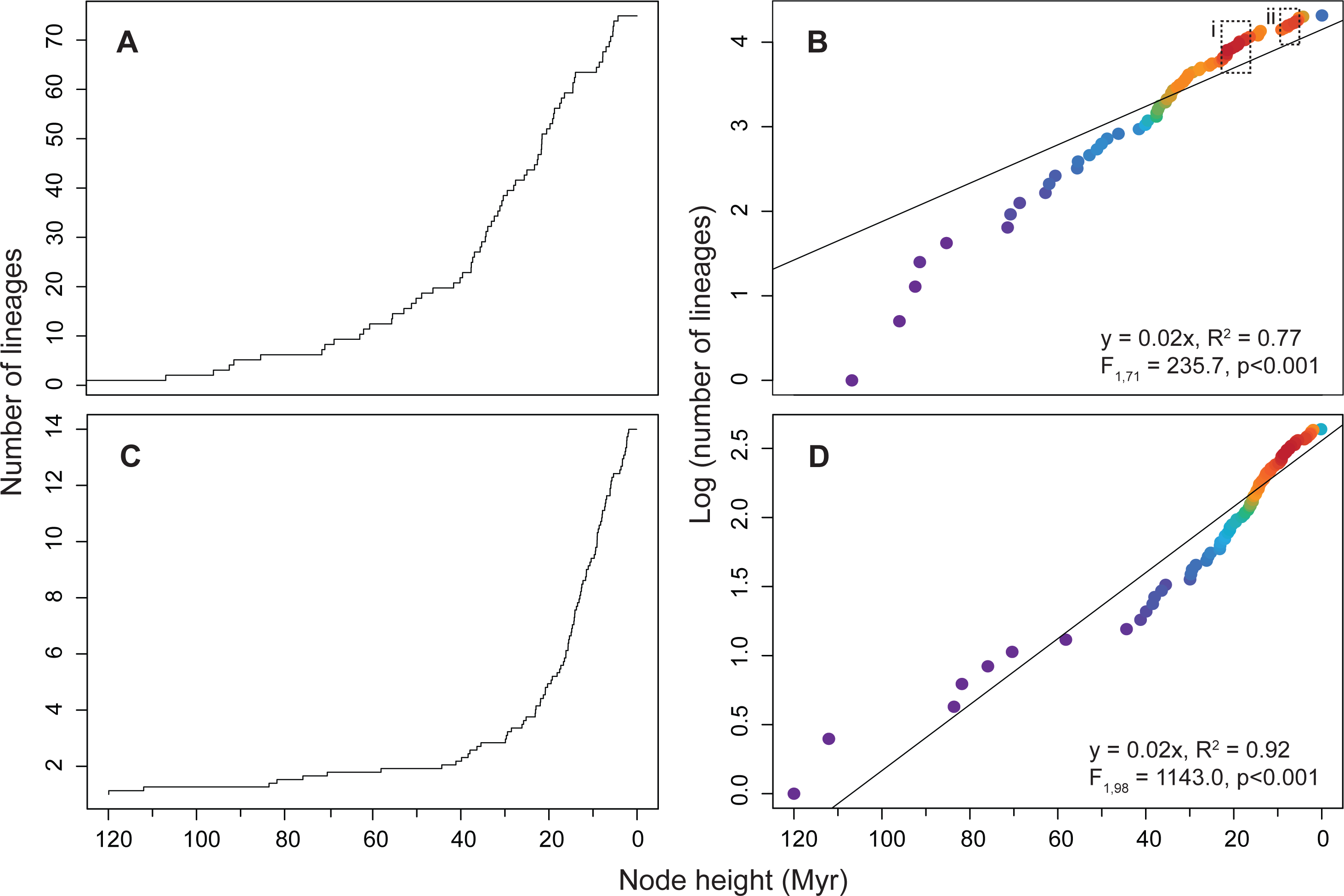
A-D. Lineage through time plots for (A-B) Strombidae only and (C-D) Stromboidea: (A, C), number of observed lineages through time; (B, D), log-transformed observed lineages through time, coloured by density (red = high density, purple = low density). The BEAST tree used was a chronogram; therefore, node heights on the x-axis correspond to time (Myr). A linear model (black line) was fit using R; adjusted R^2^ values, F-statistic, degrees of freedom and p-value of the linear model are reported. Periods of highest increase in cladogenesis marked by boxes (dashed black lines): i, ca. 17–23 Mya (Early Miocene); ii, ca. 6–9 Mya (Late Miocene).

## 4 Discussion

### 4.1 Phylogenetic analysis

The rapidly changing systematics of Stromboidea, and Strombidae in particular, is driven by a substantial scientific interest and a large morphological diversity of shells within the groups (Fig. 1). Since the first molecular phylogeny of Strombidae (Latiolais et al., 2006), the number of strombid genera has dramatically increased (e.g., Bandel, 2007; Dekkers, 2012a; Dekkers and Maxwell, 2020; Liverani et al., 2021); yet sparse molecular sampling has hindered discussions of relationships. This more taxonomically comprehensive phylogenetic study is therefore a much-needed step in shaping our understanding of strombid systematics and highlights the need of both molecular and morphological data to robustly test taxonomic and phylogenetic hypotheses. Using molecular data, this study shows that several stromboidean groups require systematic revision. Here, all relationships among strombid genera as defined in this study were recovered with moderate to maximal support, except for the clade sister to *Lentigo*, which was recovered with low support (PP = 0.59) (Fig. 3). These results differ from the strombid phylogenies containing fewer taxa but produced with mitogenome data (Irwin et al., 2021; Machkour-M’Rabet et al., 2021), which resolved (*Strombus* + *Aliger*) + (*Conomurex* + ((*Harpago* + *Lambis*) + (*Tridentarius* + *Dolomena*))) with varied support. Additional mitogenome sequences in Li et al. (2022a) resulted in the same topology, plus with *Lentigo* as sister to all other strombids, and *Euprotomus* Gill, 1870 as sister to *Strombus* + *Aliger*.

#### 4.1.1 Systematics

Results suggest that six stromboid genera are not monophyletic using current generic assignments, and that several species boundaries differ from current species concepts (Fig. 2; Suppl. Mat. 4, 5). As Strombidae was the focus of the sampling effort, changes to generic boundaries are only proposed for this family for genera not recovered as monophyletic, except for *Aliger*. We also raise *Lambis sowerbyi*, *Harpago rugosus,* and *Rimellopsis laurenti* (formerly *L. truncata sowerbyi, H. chiragra rugosus* and *R. powisii*, respectively), to species level based on species delimitations (Tables 3, 4; Fig. 2; Suppl. Mat. 5, 4). These species, as well as all other newly recognised species in this study, were recovered as monophyletic in at least one of the single gene trees (where data were available) (Table 2).

Here, we redefine *Aliger* to include *Macrostrombus* and *Eustrombus*, based on the short internodal distances among these genera, although they are reciprocally monophyletic (Table 2). The clade including these genera was recovered with maximal support (Fig. 3) and shares general morphological characters with nearly all species from the Tropical Eastern Pacific and Tropical West Atlantic (e.g., expanded outer lip, glazed parietal area). Analyses did not recover the monophyly of *Lentigo* (Figs 2–3; Suppl. Mat. 5–8); thus, we recognise *Thersistrombus* Bandel, 2007 as a synonym of *Lentigo* (Table 2). This is a morphologically diverse clade, and we could find no shared characters between *Thersistrombus* and *Lentigo* except for a columellar callus that is extended on the ventral side of the shell. As such, this grouping is a conservative measure based on the paraphyly of *Lentigo*, and exploration of further characters, including anatomical and shell micro sculpture characters is needed. *Canarium*, *Dolomena* and *Doxander* were polyphyletic in all analyses (Figs 2–3; Suppl. Mat. 5–8). The most parsimonious solution to this, based on the BEAST chromogram, is to recognise the following groups (listed here as A-G) at the genus level; however, we are unable to propose strong characters in support of *Canarium* and *Dolomena* as defined here. (A) As the clade sister to *“C.” wilsonorum* had only moderate support (PP = 0.91) and *“C”. wilsonorum* bears a morphological similarity to the type of *Canarium*, this species may belong to *Canarium*. Further molecular work may help to resolve this; thus, *“C”. wilsonorum* is assigned to *“Canarium”* as a conservative measure instead of introducing a new genus name (Table 2). Note that these results do not support the former assignment of *“C”. wilsonorum* to *Conundrum* (Liverani et al., 2021) (Fig. 3). (B-C) The clade sister to *Fusistrombus fusiformis* + *Hawaiistrombus scalariformis* received only moderate support (PP = 0.91). As these generic names already exist, *Fusistrombus* is kept as valid (with *Neostrombus* as a junior objective synonym) and *Hawaiistrombus* (type: *Strombus hellii*) is accepted, given the morphological disparity between these taxa (Table 2). (D-E) *Ministrombus* (excluding *Dolomena variabilis*) and *Maculastrombus* (including *M. microurceus* and *M. mutabilis*) are retained (Table 2). (F) Thus, *Canarium* is restricted to the remaining currently assigned species. (G) Due to the polyphyly of *Dolomena* and the low resolution of relationships in the clade containing *Neodilatilabrum, Dolomena, Doxander, Labiostrombus* and *Laevistrombus*, this clade is redefined as *Dolomena*, resolved with maximal support (Fig. 3; Suppl. Mat. 2). The relationships of type species *Dolomena pulchella* are unsupported; thus, the true name-bearing clade comprising *Dolomena* is uncertain.

#### 4.1.2 Hybrids in Strombidae

The literature contains several references to strombid specimens possessing intermediate shell morphologies, interpreted as putative intrageneric (e.g., Dekkers, 2012b; Kronenberg, 2013; Liverani & Wieneke, 2016) and intergeneric hybrids, though none were tested with molecular data. While claims of intrageneric hybrids are plausible, intergeneric hybrids (*Latissistrombus latissimus* x *L. lambis*, *Conomurex decorus* x *Gibberulus gibberulus*, *L. latissimus* x *H. chiragra*; Kronenberg, 2008; Dekkers and Maxwell, 2018; Maxwell and Ocean, 2022) should be viewed with suspicion, as these occur between lineages separated for substantial periods, according to the chronogram: *Latissistrombus* x *Lambis* or *Harpago,* 71 Mya (95% HPD: 56–88 Mya); *Conomurex* x *Gibberulus,* 53 Mya (95% HPD: 38–69 Mya) (Fig. 3). As far as we are aware, intergeneric hybrids are unknown in natural molluscan populations, and appear in very few other groups (e.g., Franco-Trecu et al., 2016). Molecular data will be informative for understanding these taxa.

### 4.2 Divergence and dispersal within the evolutionary history of Stromboidea

Divergence time estimation using molecular data can offer a new perspective on the evolutionary history of a group, although conflicts with the fossil record often add uncertainty to results. Here, Strombidae is estimated to have originated during the Cretaceous, which predates the oldest undisputed strombid fossil by at least 59 Myr (Fig. 3; Table 1). This could be attributed to inadequacy of the model (Brown and Smith, 2018) and/or incompleteness of the fossil record; for example, the fossil record in the Indo-West Pacific (IWP) is poorly studied (Harzhauser et al., 2018), despite being the modern global biodiversity hotspot. Preservation is also often poor in these gastropod fossils due to the instability of aragonitic shells (by contrast, calcite preserves well; James et al., 2005; Cherns et al., 2009), which is particularly problematic when the oldest fossils lack distinctive characteristics. Thus, the following discussion on historical biogeography and diversification is tentative and warrants further exploration.

#### 4.2.1 Timing and causes of diversification

Time-calibrated phylogenies facilitate discussion of changes in cladogenesis rate relative to other periods in time. However, note that lineages through time (LTT) plots, while useful, are based on extant species and do not consider 95% highest probability density intervals - this is often problematic for reconstructing evolutionary histories (Figs 3–4) (Louca and Pennell, 2020; Helmstetter et al., 2022). In the fossil record, the number of strombid genera increases during the Miocene (Kronenberg and Harzhauser, 2012); similarly, our results suggest increases in the rate of cladogenesis in the Early and Late Miocene (Fig. 4B). Increased cladogenesis of other shallow-water gastropods in the Miocene (Kohn, 1990; Vermeij, 1996; Meyer, 2003; Williams, 2007; Williams and Duda, 2008; Reid et al., 2010) are often associated with the eastward shift of the global biodiversity hotspot from the Tethys to its current position in the central IWP (Wilson and Rosen 1998; Renema et al., 2008; Leprieur et al., 2016). This shift is attributed to tectonic events, including the Early Miocene formation of the Gomphotherium land bridge, which limited Tethys-IWP faunal exchange (Harzhauser et al., 2007), and the collision of the Australia and New Guinea plate with Pacific arcs and the southeast Asian plate margin ca. 25 Mya, which led to the creation of shallow-water habitats and extended coastlines (Kohn, 1990; Wilson and Rosen 1998; Meyer, 2003; Williams and Duda, 2008). These events facilitated the expansion of seagrass habitats and diversification of zooxanthellate corals ca. 20–25 Mya, increasing habitat complexity and cladogenesis rate of many benthic groups (Brasier, 1975; Reuter et al., 2011). As strombids are predominately associated with seagrass beds and coral rubble (Stoner and Waite, 1991), this event also may have driven cladogenesis within Strombidae, approximately 23 Mya (Fig. 4).

By contrast, relative cladogenesis analyses suggested several earlier increases in diversification rate in the Middle Eocene (p <0.01) (Fig. 3). This supports the progressive increase in taxonomic diversity suggested in the fossil record, from the Early Cenozoic to a Middle Eocene maximum in the western Tethys (Crame and McGowan, 2022). A pulse of diversification is particularly evident within the Anglo-Paris Basin; for example, the wide range of molluscs found within the Calcaire Grossier Formation is similar in biodiversity to the present day IWP hotspot (Cossmann and Pissarro, 1904–1913; Merle, 2008). By contrast, aporrhaid diversity and geographic distribution gradually declined during the Early Cenozoic, after the K/Pg mass extinction (Roy 1994). Roy (1996) suggested that this coincided with the reciprocal rise in strombid and rostellariid genera during the Paleocene and Eocene and may reflect a biotic replacement event and/or different responses to the same environmental change.

#### 4.2.2 Biogeographical reconstruction

Interestingly, time-calibrated analyses within this study suggest similar divergence times between East Pacific/Atlantic (EP/A) and IWP sister groups in the shallow-water Strombidae (92 Mya) and largely offshore Xenophoridae (88 Mya) (Fig. 3). Some biogeographic barriers to dispersal likely had a significant impact on both offshore and shallow-water groups, such as the rise of the Isthmus of Panama (Macpherson et al., 2010). In Strombidae and Xenophoridae, divergence between EP/A and IWP sister groups followed the initiation of the widening of the Atlantic (110–65 Mya), which constituted a significant oceanic barrier to larval dispersal (Hou and Li, 2018). For larvae crossing the modern Atlantic via one of the main surface currents, the journey is estimated to take 40–60 days at 14–90 cm s^-1^ (Scheltema, 1971). The pelagic larval duration of *Aliger gigas* is ca. 60 days (Stoner, 2003), making larval drift more likely before the Atlantic opening was at its widest.

Previous molecular studies supported a monophyletic EP/A strombid clade derived from a paraphyletic IWP clade (Latiolais et al., 2006; Irwin et al., 2021; Machkour-M’Rabet et al., 2021). By contrast, our results for both Strombidae and Xenophoridae resolve an IWP clade sister to an EP/A clade, together sister to a second, more diverse IWP clade (Fig. 3), which supports a previous study with lower taxon sampling (Irwin et al., in review). Ancestral range reconstructions suggest an IWP origin for strombids (Fig. 3), which is not in conflict with the Tethyan origin supported by the oldest definitive strombid fossil (Bayan, 1870; Wieneke et al., 2023; Table 1). In the IWP, the strombid fossil record extends only to the Miocene (e.g. Bose et al., 2021), except for one incomplete specimen assigned to Strombidae from the Middle Eocene of Indonesia (*Jogjacartanus sultani*; Leloux and Wesselingh 2009, p. 657, pl. 235, figs 6, 7). Although this paucity is attributed to the Miocene shift in biodiversity towards the IWP, there is still much about the pre-Miocene IWP that remains unknown.

### 4.3 Future work

It is crucial to recognise that putative species discussed here require further exploration within an integrative taxonomic framework (Puillandre et al., 2012, 2020). Firstly, in all instances where “cryptic lineages” were identified, we were unable to confidently assign any to the typical form. Both *Lambis lambis* and *Maculastrombus mutabilis* have several historical synonyms that might be appropriate for some of the cryptic species revealed by these results. Also, sequence data for two putative cryptic species, *“C”. wilsonorum* B and *L. lambis* C, have no voucher data (Suppl. Mat. 2). We note that the *L. lambis* cryptic species reported by Li et al. (2022b) are here identified as *L. sowerbyi* and *L. lambis* B; nevertheless, our results suggest a species complex within *L. lambis* that requires further attention. We also note that the geographic range of *Dolomena variabilis* (formerly *Ministrombus*) extends to New Caledonia (Fig. 2; Suppl. Mat. 2, 5), which Maxwell (2022) restricts to *M. caledonicus*, indicating the species may have been oversplit taxonomically. We strongly recommend further investigation into the species complex of *T. terebellum*; while four cryptic species were identified (*T. terebellum* A-D), some were only represented by juveniles within this study. This, together with highly variable shell patterns and overlapping geographic regions (Table 2), means that we could not exclude the possibility that one or two of these species could represent *T. hubrechti* Poppe and Tagaro, 2016 or *T. simoni* Dekkers, S. J. Maxwell and Congdon, 2019. Furthermore, the shell colour pattern on the type specimen of *T. terebellum* is faded (Jung and Abbott, 1967, pp. 445–454, pl. 323), which complicates identification of the type species. Finally, two xenophorid genera (*Xenophora* and *Onustus*), and four strombid genera (*Lentigo, Canarium, Dolomena* and *Doxander*) are not monophyletic and these groups require further study with increased taxon sampling to resolve their relationships, particularly those of Xenophoridae; 63% of currently accepted xenophorid species currently lack molecular data, including the new genus *Ponderiana* Nappo, Bini & Santucci, 2022. In particular, *Canarium* and *Dolomena* should be revisited more fully with regards to both morphological and molecular data, given that this study lacks the type species for *Canarium* and that the deeper nodes containing *Dolomena pulchella* (type species of *Dolomena*) are unsupported (Fig. 3).

## 5 Conclusion

This study advances our understanding of stromboidean systematics, as well as the patterns of speciation and biogeography which have shaped the evolution of the superfamily. We provide the largest phylogenetic analysis for Stromboidea so far, focusing on relationships within Strombidae. The time-calibrated analysis supports the hypothesis that changes in habitat resulting from tectonic activity at the Oligocene/Miocene boundary led to a period of rapid cladogenesis for both shallow-water and offshore gastropods in the Central IWP. Interestingly, LTT plots suggest two distinct pulses of strombid diversification within the Miocene. This radiation occurred earlier than is currently known from the fossil record; however, both ancestral range reconstructions and fossils support a Tethyan/IWP origin for Strombidae.

## Acknowledgements

Some material in this study originates from several research expeditions organized for the *Our Planet Reviewed* (MNHN and ProNatura International) and *Tropical Deep-Sea Benthos* (MNHN and the Institut de Recherche pour le Développement) programs: Philippines, PANGLAO 2004, AURORA 2007; Papua New Guinea, BIOPAPUA, MADEEP, KAVIENG 2014, PAPUA NIUGINI; Vanuatu, SANTO 2006; New Caledonia, CONCALIS, EXBODI, KANACONO, KOUMAC; French Polynesia, TUHAA PAE 2013; Madagascar, ATIMO VATAE, MIRIKY; Mozambique, INHACA 2011; French Caribbean, KAURUBENTHOS, KARUBENTHOS 2, MADIBENTHOS; French Guiana, GUYANE 2014, ILES DU SALUT; Senegal, Dakar’09 – see expeditions.mnhn.fr for more information. Expeditions operated in accordance with the Nagoya Protocol for access to genetic resources. For specimen loans, we thank Nicolas Puillandre (MNHN), Andreia Salvador (NHMUK), Yasunori Kano (University of Tokyo), John Slapcinsky (University of Florida), Serge Gofas (University of Málaga), José Templado (Museo Nacional de Ciencias Naturales, MNCN), Tan Koh Siang (National University of Singapore, NUS), Tan Siong Kiat (NUS), Richard Willan (Museum and Art Gallery of the Northern Territory), Bruce Marshall (Museum of New Zealand Te Papa Tongarewa), Kerry Walton (NMNZ), Annie Machordom (MNCN) and Fabio Crocetta (Stazione Zoologica Anton Dohrn, SZN). For curation of vouchers, we thank Virginie Héros, Barbara Buge, Julien Brisset (all MNHN), Andreia Salvador (NHMUK), Serge Gofas, Carlos Jimenez (Enalia Physis Environmental Research Centre), Magdalene Papatheodoulou (Department of Fisheries and Marine Research, Cyprus), and Elisa Cenci (SZN). We are grateful to Claire Griffin (NHMUK) for Sanger sequencing. We thank the USNM for providing images of USNM PAL 105677, and the following people for images in Fig. 1: A, F, G, David Massemin; B, Gilles Devauchelle (MNHN); C, Barbara Buge; D, MNHN Kavieng; E, H, Laurent Charles (MNHN).

