## Supplementary Material 1 for "Molecular phylogenetics of the superfamily Stromboidea (Caenogastropoda): new insights from increased taxon sampling"

**Supplementary Material 1.** Currently recognized stromboidean genera (MolluscaBase, 2023), with their type species, arranged by family (MolluscaBase, 2023). Mode of designation of type species: OD = original designation; M = monotypy; SD = subsequent designation; AT = absolute tautonymy; TRN, typification of replaced name. Asterisks (*) mark taxa that are not represented in the phylogeny.

| Family | Genus | Type species |
|---|---|---|
| **Aporrhaidae Gray, 1850** | *Aporrhais* da Costa, 1778 | *Aporrhais quadrifidus* da Costa, 1778 (M) [= *Aporrhais pespelecani* (Linnaeus, 1758)] |
| | *Arrhoges* Gabb, 1868 * | *Rostellaria occidentalis* H. Beck, 1836 * (M) |
| **Rostellariidae Gabb, 1868** | *Rimellopsis* Lambiotte, 1979 | *Rostellaria powisii* Petit de la Saussaye, 1840 (OD) |
| | *Rostellariella* Thiele, 1929 | *Rostellaria delicatula* G. Nevill, 1881 (OD) |
| | *Tenuitibia* Dekkers, 2020 | *Gladius martinii* Marrat, 1877 (OD) |
| | *Tibia* Röding, 1798 | *Murex fusus* Linnaeus, 1758 (SD) |
| | *Varicospira* Eames, 1952 | *Strombus cancellatus* Lamarck, 1816 (OD) |
| **Seraphsidae Gray, 1853** | *Terebellum* Bruguière, 1798 | *Conus terebellum* Linnaeus, 1758 (subsequent M) |
| **Strombidae Rafinesque, 1815** | *Aliger* Thiele, 1929 | *Strombus gallus* Linnaeus, 1758 (OD) |
| | *Barneystrombus* Blackwood, 2009 | *Strombus kleckhamae* Cernohorsky, 1971 (OD) |
| | *Canarium* Schumacher, 1817 | *Canarium ustulatum* Schumacher, 1817 (M) [= *Canarium urceus* (Linnaeus, 1758) *] |
| | *Conomurex* P. Fischer, 1884 | *Strombus luhuanus* Linnaeus, 1758 (M) |
| | *Dolomena* Wenz, 1940 | *Strombus pulchellus* Reeve, 1851 (OD) |
| | *Doxander* Wenz, 1940 | *Strombus vittatus* Linnaeus, 1758 (OD) |
| | *Euprotomus* Gill, 1870 | *Strombus aurisdianae* Linnaeus, 1758 (M) |
| | *Fusistrombus* Bandel, 2007 | *Strombus fusiformis* G. B. Sowerby II, 1842 (OD) |
| | *Gibberulus* Jousseaume, 1888 | *Strombus gibberulus* Linnaeus, 1758 (M) |

| | | |
|---|---|---|
| | *Harpago* Mörch, 1852 | *Lambis harpago* Röding, 1798 (AT) |
| | *Hawaiistrombus* Bandel, 2007 | *Strombus hellii* Kiener, 1843 (OD) * |
| | *Labiostrombus* Oostingh, 1925 | *Strombus epidromis* Linnaeus, 1758 (TRN) [*Gallinula* Mörch, 1852, non Brisson, 1760] |
| | *Laevistrombus* Abbott, 1960 | *Strombus canarium* Linnaeus, 1758 (OD) * |
| | *Lambis* Röding, 1798 | *Strombus lambis* Linnaeus, 1758 (AT) |
| | *Latissistrombus* Bandel, 2007 | *Strombus taurus* Reeve, 1857 (OD) |
| | *Lentigo* Jousseaume, 1886 | *Strombus lentiginosus* Linnaeus, 1758 (M) |
| | *Lobatus* [Swainson], 1837 | *Strombus bituberculatus* Lamarck, 1822 (M) |
| | *Macrostrombus* Petuch, 1994 | *Strombus costatus* Gmelin, 1791 (OD) |
| | *Maculastrombus* Liverani, Dekkers & S. J. Maxwell, 2021 | *Strombus maculatus* G. B. Sowerby II, 1842 (OD) |
| | *Ministrombus* Bandel, 2007 | *Strombus minimus* Linnaeus, 1771 (OD) |
| | *Mirabilistrombus* Kronenberg, 1998 * | *Strombus listeri* T. Gray, 1852 (OD) * |
| | *Neodilatilabrum* Dekkers, 2008 | *Strombus marginatus* Linnaeus, 1758 (OD) * |
| | *Ophioglossolambis* Dekkers, 2012 | *Strombus digitatus* Perry, 1811 (OD) |
| | *Persististrombus* Kronenberg & H. G. Lee, 2007 | *Strombus granulatus* Swainson, 1822 (OD) |
| | *Strombus* Linnaeus, 1758 | *Strombus pugilis* Linnaeus, 1758 (SD) |
| | *Terestrombus* Kronenberg & Vermeij, 2002 | *Lambis fragilis* Röding, 1798 (OD) |
| | *Thersistrombus* Bandel, 2007 | *Strombus thersites* Swainson, 1823 (OD) |
| | *Thetystrombus* Dekkers, 2008 | *Strombus latus* Gmelin, 1791 (OD) |
| | *Titanostrombus* Petuch, 1994 | *Strombus goliath* Schröter, 1805 (OD) * |
| | *Tricornis* Jousseaume, 1886 | *Strombus tricornis* [Lightfoot], 1786 (M) |
| | *Pelicaria* Gray, 1857 | *Buccinum vermis* Martyn, 1784 (M) |

| | | |
|---|---|---|
| **Struthiolariidae Gabb, 1868** | *Perissodonta* E. von Martens, 1878 * | *Struthiolaria mirabilis* E. A. Smith, 1875 SD) * |
| | *Struthiolaria* Lamarck, 1816 | *Struthiolaria nodulosa* Lamarck, 1816 (M) [=*Struthiolaria papulosa* (Martyn, 1784)] |
| | *Tylospira* G. F. Harris, 1897 * | *Tylospira scutulata* (Gmelin, 1791) (M) * |
| **Xenophoridae Troschel, 1852 (1840)** | *Aspidophoreas* Nappo, Bini & Santucci, 2022 | *Trochus chinensis* R. A. Philippi, 1841 (OD) |
| | *Onustus* Swainson, 1840 | *Trochus indicus* Gmelin, 1791 (OD) |
| | *Ponderiana* Nappo, Bini & Santucci, 2022 * | *Xenophora digitata* E. von Martens, 1878 (OD) * |
| | *Stellaria* Möller, 1832 | *Trochus solaris* Linnaeus, 1764 (M) |
| | *Xenophora* Fischer von Waldheim, 1807 | *Xenophora laevigata* Fischer von Waldheim, 1807 (SD) [=*Xenophora conchyliophora* (Born, 1780)] |
