## Supplementary Material 2 for "Molecular phylogenetics of the superfamily Stromboidea (Caenogastropoda): new insights from increased taxon sampling"

**Supplementary Material 2.** List of specimens used in study with delimited putative species number, family and species name, sampling locality, registration number and GenBank Accession number(s). Species listed in order of clades named in Figure 2, and Suppl. Mat. 4, 5, followed by outgroups used in MrBayes and IQ-TREE analyses (Suppl. Mat. 6, 8). Sequences lacking COI data were not included in delimitation analyses; these were delimited based on single genes trees (Suppl. Mat. 8), and, if no other members of that species were available, were listed as separate putative species. Specimens used in BEAST analyses as representatives of their respective putative species are highlighted in bold, with asterisks (*) marking any chimeras used. All sequences published prior to this study are listed with their respective publication references, with a complete reference list below the table. ***Abbreviations (1: NHMUK material):*** NHMUK – Natural History Museum, London, UK. ***Abbreviations (2: loaned material):*** AORI – Atmosphere and Ocean Research Institute, University of Tokyo, Japan; MNCN – Museo Nacional de Ciencias Naturales; MNHN – Muséum National d'Histoire Naturelle, Paris, France; NMNZ – Museum of New Zealand Te Papa Tongarewa; NTM – Museum and Art Gallery of the Northern Territory, Darwin, Australia; SZN – Stazione Zoologica Anton Dohrn, Italy; UF – Florida Museum of Natural History, University of Florida, USA; ZRC – Zoological Reference Collection, Raffles Museum of Biodiversity Research, National University of Singapore. ***Abbreviations (3: GenBank sequences):*** AM – Australian Museum; ANSP – Academy of Natural Sciences of Drexel University, Philadelphia, USA; BMOO – Moorea Biocode Project; DZMB – German Centre for Marine Biodiversity Research, Senckenberg Institute; ECOCHM – Barcode of Wildlife Project Mexico; FMNH – Field Museum of Natural History, Chicago, USA; GNM – Gothenburg Natural History Museum; HNU – Hainan University, China; KMRS – Kristineberg Marine Research Station, Sweden; NTOU – National Taiwan Ocean University; OUC – Ocean University of China, Qingdao; SAU – Sher-e-Bangla Agricultural University, Dhaka, Bangladesh; TMBC – Tropical Marine Biodiversity Collections of South China Sea; UNAL – Universidad Nacional de Colombia; USNM – Smithsonian National Museum of Natural History, D.C., USA; NV – no voucher; NL – no locality.

| N. | Species | Sampling locality | Latitude | Longitude | Voucher no. | COI | 12S | 16S | 28S | Reference | Comments |
|---|---|---|---|---|---|---|---|---|---|---|---|
| 1 | *Onustus indicus* (Gmelin, 1791) | **1. Nosy Be, Madagascar** | – | – | **UF 424024** | **OR910869** | **OR934161** | **OR934297** | **OR934039** | **This study** | |
| 2 | *Aspidophoreas chinensis* (R. A. Philippi, 1841) | **1. Off Totoro, Nobeoka, Kyushu Is., Japan** | – | – | **AORI YK#4954** | **OR910766** | **OR934176** | **OR934312** | **OR933986** | **This study** | |

| | | | | | | | | | | |
|---|---|---|---|---|---|---|---|---|---|---|
| 3 | *Stellaria solaris* (Linnaeus, 1764) | **1. St. Martin's island, Bangladesh** | **–** | **–** | **SAU M1903SM-03** | **MN703099** | | | | **Habib et al. (2019)** |
| 4 | *Onustus exutus* (Reeve, 1842) | **1. Zhoushan, Zhejiang province, China** | **30.0833** | **122.1** | **NV** | **MK327366** | **MK327366** | **MK327366** | | **Xu et al. (2019)** |
| 5 | *Xenophora solarioides* (Reeve, 1845) | **1. Double Is., New Caledonia** | **-20.4648** | **164.1283** | **MNHN-IM-2013-84859** | **OR911259** | **OR934433** | **OR934484** | **OR934383** | **This study** |
| | | 2. Double Is., New Caledonia | -20.4648 | 164.1283 | MNHN-IM-2013-84860 | OR910997 | OR934198 | OR934336 | OR934067 | This study |
| 6 | *Xenophora pallidula* (Reeve, 1842) | **1. Muttom, Kanyakumari, Tamil Nadu, India** | – | – | **ZRC MOL.9417** | **OR911258** | **OR934432** | **OR934483** | **OR934382** | **This study** |
| 7 | *Xenophora japonica* Kuroda & Habe, 1971 | 1. Miura Peninsula, Honshu Is., Japan | – | – | AORI YK#4953 | OR910996 | OR934197 | OR934335 | OR934066 | This study |
| | | **2. Off Oosezaki, Suruga Bay, Honshu Is., Japan** | **35.05** | **138.7833** | **AORI YK#4037** | **MW244823** | **MW244823** | **MW244823** | **MW244823** | **Irwin et al. (2021)** |
| 8 | *Onustus longleyi* (Bartsch, 1931) | 1. SW Dry Tortugas Keys, Florida, USA | – | – | UF 381274 | OR910870 | OR934162 | OR934298 | OR934040 | This study |
| | | **2. Straits of Florida, USA** | – | – | **UF 323747** | **OR910871** | **OR934163** | **OR934299** | **OR934041** | **This study** |
| 9 | *Xenophora conchyliophora* (Born, 1780) | **1. SW Marquesas Keys, Florida, USA** | **24.5037** | **-82.258** | **UF 26680** | **OR911257** | **OR934431** | **OR934482** | | **This study** |
| 10 | *Onustus caribaeus* (Petit de la Saussaye, 1857) | **1. Off St. Petersburg, Florida, USA** | **27.7388** | **-84.6033** | **UF 351181** | **OR911243** | **OR934417** | **OR934468** | **OR934368** | **This study** |
| | | 2. Off St. Petersburg, Florida, USA | 27 | 84 | UF 351147 | OR910868 | OR934160 | OR934296 | | This study |
| 11 | *Struthiolaria papulosa* (Martyn, 1784) | **1. Orewa Beach, N Auckland, New Zealand** | **-36.5833** | **174.7** | **AORI YK#4036** | **MW244818** | **MW244818** | **MW244818** | **MW244818** | **Irwin et al. (2021)** |

| | | | | | | | | | | |
|---|---|---|---|---|---|---|---|---|---|---|
| | | 2. Waitata Reach, South Is., New Zealand | -41.0022 | 173.9138 | NMNZ M.318561/2 | OR910961 | OR934179 | OR934315 | OR934052 | This study |
| 12 | *Pelicaria vermis* (Martyn, 1784) | **1. Waitata Reach, South Is., New Zealand** | **-41.0255** | **173.9075** | **NMNZ M.318596/1** | **OR911246** | **OR934420** | **OR934471** | **OR934371** | **This study** |
| | | 2. Waitata Reach, South Is., New Zealand | -41.0255 | 173.9075 | NMNZ M.318596/1 | OR910872 | OR934167 | OR934303 | OR934042 | This study |
| 13 | *Aporrhais serresiana* (Michaud, 1828) | 1. Gulf of Naples, Italy | – | – | SZN MOL0047 | OR910711 | OR934075 | OR934212 | OR933972 | This study |
| | | **2. Málaga Bay, Spain** | **36.6135** | **-4.3718** | **MNCN 15.05/95247** | **MW244817** | **MW244817** | **MW244817** | **MW244817** | **Irwin et al. (2021)** |
| | | 3. Málaga Bay, Spain | 36.6135 | -4.3718 | MNCN 15.05/95247 | OR910712 | OR934077 | OR934213 | OR933973 | This study |
| 14 | *Aporrhais pespelecani* (Linnaeus, 1758) | 1. Øresund, Sweden | 55.865 | 12.747 | GNM Gastr 8426V | MG935309 | | | | Lundin (2018) |
| | | 2. Bergen, Norway | 60.3703 | 2.4784 | DZMB MT01020 | KR084699 | | | | Barco et al. (2016) |
| | | 3. Bergen, Norway | 60.3703 | 2.4784 | DZMB MT01018 | KR084761 | | | | Barco et al. (2016) |
| | | 4. Bergen, Norway | 60.3703 | 2.4784 | DZMB MT01019 | KR084497 | | | | Barco et al. (2016) |
| | | 5. Kattegatt, Sweden | 57.055 | 11.717 | GNM Gastr 8420V | MG935118 | | | | Lundin (2018) |
| | | 6. Bergen, Norway | 60.3703 | 2.4784 | DZMB MT01021 | KR084824 | | | | Barco et al. (2016) |
| | | 7. Gullmarsfjord, Sweden | 58.2833 | 11.5167 | KMRS MGE19 | EF528304 | | | | Bourlat et al. (2008) |
| | | 8. Bergen, Norway | 60.3703 | 2.4784 | DZMB MT01017 | KR084735 | | | | Barco et al. (2016) |
| | | **9. E Isle Arran, Scotland** | **55.5849** | **-4.9529** | **NHMUK 20230924** | **OR911210** | **OR934384** | **OR934434** | **PP001795** | **This study** |

| | | | | | | | | | | | |
|---|---|---|---|---|---|---|---|---|---|---|---|
| | | 10. Scotland | – | – | NV | DQ525241 | | | | Latiolais et al. (2016) | |
| | | 11. Málaga Bay, Spain | 36.6135 | -4.3718 | MNCN 15.05/95248 | | OR934074 | | | This study | |
| | | **12. E Dundee, NE Scotland** | 56.4948 | -2.3203 | **NHMUK 20230925** | | **OR934076** | | | **This study** | |
| 15 | *Terebellum terebellum* A | **1. Baclayon, Bohol Is., Philippines** | **9.6233** | **123.9083** | **MNHN-IM-2007-33342** | **OR910971** | **OR934193** | **OR934330** | | **This study** | Four putative cryptic species. True *Terebellum terebellum* Kuroda & Kawamoto, 1956 is type species of *Terebellum.* |
| 16 | *Terebellum terebellum* B | 1. Alexishafen, Papua New Guinea | -5.09 | 145.81 | MNHN-IM-2013-12885 | OR910972 | | | | This study | |
| | | 2. Alexishafen, Papua New Guinea | -5.0883 | 145.8017 | MNHN-IM-2013-13909 | OR910973 | | | | This study | |
| | | **3. Alexishafen, Papua New Guinea** | **-5.09** | **145.81** | **MNHN-IM-2013-12884** | **MW244821** | **MW244821** | **MW244821** | **MW244821** | **Irwin et al. (2021)** | |
| | | 4. N Riwo Is., Papua New Guinea | -5.1417 | 145.8083 | MNHN-IM-2013-17031 | OR910980 | | | | This study | |
| | | 5. N Kabanam Point, Papua New Guinea | -5.1 | 145.8033 | MNHN-IM-2013-13905 | OR910967 | OR934189 | OR934326 | OR934055 | This study | |
| 18 | *Terebellum terebellum* C | **1. Plateau Karembé (Pte Nord), New Caledonia** | **-20.6267** | **164.2883** | **MNHN-IM-2019-1737** | **OR911253** | **OR934427** | **OR934478** | **OR934378** | **This study** | |
| 19 | *Terebellum terebellum* D | 1. Biliau Is., Papua New Guinea | -5.2 | 145.8017 | MNHN-IM-2013-11701 | OR910975 | | | | This study | |
| | | 2. Kavieng Lagoon, Papua New Guinea | -2.625 | 150.775 | MNHN-IM-2013-46856 | OR910977 | | | | This study | |
| | | 3. S Yabob Is., Papua New Guinea | -5.2583 | 145.7883 | MNHN-IM-2013-17611 | OR910981 | | | | This study | |
| | | **4. Madang Harbour, Papua New Guinea** | **-5.2083** | **145.8083** | **MNHN-IM-2013-11022** | **OR910970** | **OR934192** | **OR934329** | **OR934058** | **This study** | |

| | | | | | | | | | | |
|---|---|---|---|---|---|---|---|---|---|---|
| | | 5. N Banap Damon Point, Papua New Guinea | -5.1633 | 145.8067 | MNHN-IM-2013-16805 | OR910974 | | | | This study |
| | | 6. W Kranket Is., Papua New Guinea | -5.1967 | 145.82 | MNHN-IM-2013-16786 | OR910969 | OR934191 | OR934328 | OR934057 | This study |
| 17 | *Terebellum delicatum* Kuroda & Kawamoto, 1956 | **1. Grand Passage, N Province, New Caledonia** | **-19.83** | **163.795** | **MNHN-IM-2007-35460** | **OR911254** | **OR934428** | **OR934479** | **OR934379** | **This study** |
| | | 2. Kavieng Lagoon, Papua New Guinea | -2.625 | 150.775 | MNHN-IM-2013-47193 | OR910968 | OR934190 | OR934327 | OR934056 | This study |
| | | 3. Kavieng Lagoon, Papua New Guinea | -2.625 | 150.775 | MNHN-IM-2013-47194 | OR910976 | | | | This study |
| | | 4. S Panab Is., Papua New Guinea | -5.1717 | 145.8083 | MNHN-IM-2013-16130 | OR910979 | | | | This study |
| | | 5. Madang, Papua New Guinea | -5.2367 | 145.7967 | MNHN-IM-2013-18019 | OR910978 | | | | This study |
| 20 | *Varicospira crispata* (Lamarck, 1842) | **1. E Baler Bay, Philippines** | **15.8507** | **121.8507** | **MNHN-IM-2007-34504*** | **OR910994** | | | | **This study** |
| | | 2. E Baler Bay, Philippines | 15.8507 | 121.8507 | MNHN-IM-2007-34489 | OR910995 | | | | This study |
| | | **3. Cortes, Bohol Is., Philippines** | **9.7033** | **123.8467** | **MNHN-IM-2007-33339*** | | **OR934196** | **OR934334** | | **This study** |
| 21 | *Varicospira cancellata* (Lamarck, 1816) | 1. Kavieng Lagoon, Papua New Guinea | -2.625 | 150.775 | MNHN-IM-2013-47468 | OR910992 | | OR934332 | OR934065 | This study |
| | | **2. Kavieng Lagoon, Papua New Guinea** | **-2.67** | **150.68** | **MNHN-IM-2013-55759** | **MW244822** | **MW244822** | **MW244822** | **MW244822** | **Irwin et al. (2021)** |
| | | 3. Kavieng Lagoon, Papua New Guinea | -2.6467 | 150.7367 | MNHN-IM-2013-55221 | OR910993 | OR934195 | OR934333 | | This study |
| 22 | *Tibia fusus* (Linnaeus, 1758) | **1. E Baler Bay, Philippines** | **15.8868** | **121.8868** | **MNHN-IM-2007-34496** | **OR910984** | | | | **This study** |

| | | | | | | | | | | |
|---|---|---|---|---|---|---|---|---|---|---|
| 23 | *Tibia insulaechorab* Röding, 1798 | **1. Al Salwa, Saudi Arabia** | **16.836** | **42.584** | **UF 521337** | OR911251 | OR934425 | OR934476 | OR934376 | This study |
| | | 2. Al Salwa, Saudi Arabia | 16.836 | 42.584 | UF 521328 | OR910985 | OR934187 | OR934324 | OR934060 | This study |
| 24 | *Tenuitibia martinii* (Marrat, 1877) | 1. Off the Sepik River, Papua New Guinea | -3.9333 | 144.6667 | MNHN-IM-2013-64466 | OR910962 | OR934170 | OR934306 | OR934053 | This study |
| | | 2. Gulf of Huon, S Lae, Papua New Guinea | -6.95 | 147.1333 | MNHN-IM-2013-64502 | OR910964 | | | | This study |
| | | 3. Gulf of Huon, S Lae, Papua New Guinea | -6.95 | 147.1333 | MNHN-IM-2013-64501 | OR910965 | | | | This study |
| | | 4. Gulf of Huon, S Lae, Papua New Guinea | -6.95 | 147.1333 | MNHN-IM-2013-64500 | OR910966 | | | | This study |
| | | **5. SE New Britain, Papua New Guinea** | **-6.1167** | **149.1667** | **MNHN-IM-2013-46144** | **OR910963** | **OR934171** | **OR934307** | **OR934054** | **This study** |
| 25 | *Rostellariella lorenzi* H. Morrison, 2005 | **1. Northern Territory, Timor Sea, Australia** | **-9.8333** | **128** | **NTM P050480** | **OR910952** | **OR934169** | **OR934305** | **OR934048** | **This study** |
| 26 | *Rostellariella delicatula* (G. Nevill, 1881) | 1. Off Mambare Bay, Papua New Guinea | -7.8833 | 148.05 | MNHN-IM-2013-64479 | OR910946 | | | | This study |
| | | 2. Gulf of Huon, S Lae, Papua New Guinea | -6.8667 | 147.0833 | MNHN-IM-2013-64472 | OR910941 | | | | This study |
| | | 3. Astrolabe Bay, Papua New Guinea | -5.3833 | 145.8 | MNHN-IM-2013-64471 | OR910926 | | | | This study |
| | | 4. W Kairiru Is., Papua New Guinea | -3.3167 | 143.45 | MNHN-IM-2013-18837 | OR910934 | | | | This study |
| | | **5. W Kairiru Is., Papua New Guinea** | **-3.3333** | **143.4667** | **MNHN-IM-2013-18767** | **OR911247** | **OR934421** | **OR934472** | **OR934372** | **This study** |
| | | 6. SE New Britain, Papua New Guinea | -6.1333 | 149.1667 | MNHN-IM-2013-46108 | OR910925 | OR934168 | OR934304 | OR934047 | This study |

| | | | | | |
|---|---|---|---|---|---|
| 7. Gulf of Huon, S Lae, Papua New Guinea | -6.8667 | 147.0833 | MNHN-IM-2013-64473 | OR910940 | This study |
| 8. W Kairiru Is., Papua New Guinea | -3.3167 | 143.45 | MNHN-IM-2013-18835 | OR910936 | This study |
| 9. Astrolabe Bay, Papua New Guinea | -5.3667 | 145.8 | MNHN-IM-2013-64461 | OR910931 | This study |
| 10. Off Mambare Bay, Papua New Guinea | -7.8833 | 148.05 | MNHN-IM-2013-64465 | OR910942 | This study |
| 11. Gulf of Huon, S Lae, Papua New Guinea | -6.8667 | 147.0833 | MNHN-IM-2013-64474 | OR910938 | This study |
| 12. Off Mambare Bay, Papua New Guinea | -7.8833 | 148.05 | MNHN-IM-2013-64482 | OR910944 | This study |
| 13. Astrolabe Bay, Papua New Guinea | -5.3833 | 145.8 | MNHN-IM-2013-64469 | OR910928 | This study |
| 14. Off Mambare Bay, Papua New Guinea | -7.8833 | 148.05 | MNHN-IM-2013-64464 | OR910948 | This study |
| 15. Astrolabe Bay, Papua New Guinea | -5.3833 | 145.8 | MNHN-IM-2013-64467 | OR910929 | This study |
| 16. W Kairiru Is., Papua New Guinea | -3.3167 | 143.45 | MNHN-IM-2013-18836 | OR910935 | This study |
| 17. Off Mambare Bay, Papua New Guinea | -7.8833 | 148.05 | MNHN-IM-2013-64478 | OR910947 | This study |
| 18. Astrolabe Bay, Papua New Guinea | -5.3833 | 145.8 | MNHN-IM-2013-64468 | OR910930 | This study |
| 19. Astrolabe Bay, Papua New Guinea | -5.3833 | 145.8 | MNHN-IM-2013-64470 | OR910927 | This study |
| 20. Gulf of Huon, S Lae, Papua New Guinea | -6.8667 | 147.0833 | MNHN-IM-2013-64476 | OR910939 | This study |

| | | | | | | | | | | |
|---|---|---|---|---|---|---|---|---|---|---|
| | | 21. Off Mambare Bay, Papua New Guinea | -7.85 | 148 | MNHN-IM-2013-64503 | OR910943 | | | | This study | |
| | | 22. Off Mambare Bay, Papua New Guinea | -7.8833 | 148.05 | MNHN-IM-2013-64462 | OR910950 | | | | This study | |
| | | 23. Astrolabe Bay, Papua New Guinea | -5.35 | 145.8 | MNHN-IM-2013-9817 | OR910951 | | | | This study | |
| | | 24. Off Mambare Bay, Papua New Guinea | -7.8833 | 148.05 | MNHN-IM-2013-64481 | OR910945 | | | | This study | |
| | | 25. N Aitape, Papua New Guinea | -3.05 | 142.3333 | MNHN-IM-2013-18963 | OR910933 | | | | This study | |
| | | 26. Off Mambare Bay, Papua New Guinea | -7.8833 | 148.05 | MNHN-IM-2013-64463 | OR910949 | | | | This study | |
| | | 27. N Aitape, Papua New Guinea | -3.05 | 142.3 | MNHN-IM-2013-18966 | OR910932 | | | | This study | |
| | | 28. Gulf of Huon, S Lae, Papua New Guinea | -6.8667 | 147.0833 | MNHN-IM-2013-64475 | OR910937 | | | | This study | |
| 27 | *Rimellopsis powisii* (Petit de la Saussaye, 1840) | **1. Lamon Bay, Philippines** | **14.4412** | **121.806** | **MNHN-IM-2007-34490** | **OR910875** | **OR934172** | **OR934308** | **OR934044** | **This study** | |
| | | 2. Lamon Bay, Philippines | 14.5515 | 121.7008 | MNHN-IM-2007-34492 | OR910877 | OR934173 | OR934309 | OR934046 | This study | |
| 28 | *Rimellopsis laurenti* Duchamps, 1992 | 1. S Urélapa Is., W Malo Is., Vanuatu | -15.6583 | 167.025 | MNHN-IM-2007-33363 | OR910920 | | | | This study | Here removed from synonymy of *Rimellopsis powisii* |
| | | 2. Passe du Solitaire, S Province, New Caledonia | -21.7667 | 166.6333 | MNHN-IM-2013-64451 | OR910894 | | | | This study | |
| | | **3. Off Thio, S Province, New Caledonia** | **-21.5333** | **166.35** | **MNHN-IM-2013-64455** | **OR911248** | **OR934422** | **OR934473** | **OR934373** | **This study** | |

| | | | | | | | | |
|---|---|---|---|---|---|---|---|---|
| 4. Scorff Passage, NW Tutuba Is., Vanuatu | -15.5433 | 167.2583 | MNHN-IM-2007-33374 | OR910921 | | | | This study |
| 5. NE Isle Nuu, New Caledonia | -22.2167 | 167.1 | MNHN-IM-2013-64445 | OR910878 | | | | This study |
| 6. Off Thio, S Province, New Caledonia | -21.5333 | 166.3667 | MNHN-IM-2013-64422 | OR910911 | | | | This study |
| 7. Off Yaté, S Province, New Caledonia | -22.1 | 167.0667 | MNHN-IM-2013-64440 | OR910924 | | | | This study |
| 8. Off Thio, S Province, New Caledonia | -21.5333 | 166.35 | MNHN-IM-2013-64429 | OR910917 | | | | This study |
| 9. Off Thio, S Province, New Caledonia | -21.5333 | 166.35 | MNHN-IM-2013-64430 | OR910896 | | | | This study |
| 10. Passe d'Ounia, S Province, New Caledonia | -21.95 | 166.9667 | MNHN-IM-2013-64433 | OR910887 | | | | This study |
| 11. Tutuba Is., Vanuatu | -15.55 | 167.2667 | MNHN-IM-2007-33340 | OR910918 | | | | This study |
| 12. Segond Channel, N Aoré Is., Vanuatu | -15.5267 | 167.2067 | MNHN-IM-2007-33360 | OR910919 | | | | This study |
| 13. Off Thio, S Province, New Caledonia | -21.5333 | 166.3667 | MNHN-IM-2013-64454 | OR910902 | | | | This study |
| 14. Off Thio, S Province, New Caledonia | -21.5333 | 166.3667 | MNHN-IM-2013-64425 | OR910914 | | | | This study |
| 15. Off Yaté, S Province, New Caledonia | -22.1 | 167.0667 | MNHN-IM-2013-64439 | OR910876 | OR934174 | OR934310 | OR934045 | This study |

| | | | | | |
|---|---|---|---|---|---|
| 16. Passe d'Ounia, S Province, New Caledonia | -21.9167 | 166.9167 | MNHN-IM-2013-64508 | OR910891 | This study |
| 17. Off Thio, S Province, New Caledonia | -21.5333 | 166.3667 | MNHN-IM-2013-64427 | OR910916 | This study |
| 18. Passe d'Ounia, S Province, New Caledonia | -21.9167 | 166.9167 | MNHN-IM-2013-64432 | OR910886 | This study |
| 19. Off Thio, S Province, New Caledonia | -21.5333 | 166.3667 | MNHN-IM-2013-64449 | OR910906 | This study |
| 20. Off Thio, S Province, New Caledonia | -21.5333 | 166.3667 | MNHN-IM-2013-64452 | OR910904 | This study |
| 21. Off Thio, S Province, New Caledonia | -21.5333 | 166.3667 | MNHN-IM-2013-64453 | OR910903 | This study |
| 22. Off Thio, S Province, New Caledonia | -21.5333 | 166.35 | MNHN-IM-2013-64448 | OR910910 | This study |
| 23. Off Yaté, S Province, New Caledonia | -22.1 | 167.0667 | MNHN-IM-2013-64457 | OR910922 | This study |
| 24. Passe d'Ounia, S Province, New Caledonia | -21.9167 | 166.9167 | MNHN-IM-2013-64456 | OR910883 | This study |
| 25. Passe d'Ounia, S Province, New Caledonia | -21.9167 | 166.9167 | MNHN-IM-2013-64506 | OR910880 | This study |

| | | | | | | |
|---|---|---|---|---|---|---|
| 26. Off Thio, S Province, New Caledonia | -21.5333 | 166.3667 | MNHN-IM-2013-64421 | OR910905 | | This study |
| 27. Off Thio, S Province, New Caledonia | -21.5333 | 166.35 | MNHN-IM-2013-64443 | OR910909 | | This study |
| 28. Off Thio, S Province, New Caledonia | -21.5333 | 166.35 | MNHN-IM-2013-64444 | OR910908 | | This study |
| 29. Off Thio, S Province, New Caledonia | -21.5333 | 166.35 | MNHN-IM-2013-64438 | OR910899 | | This study |
| 30. Off Thio, S Province, New Caledonia | -21.5333 | 166.35 | MNHN-IM-2013-64458 | OR910901 | | This study |
| 31. Off Thio, S Province, New Caledonia | -21.5333 | 166.3667 | MNHN-IM-2013-64420 | OR910895 | | This study |
| 32. Off Thio, S Province, New Caledonia | -21.5333 | 166.3667 | MNHN-IM-2013-64424 | OR910913 | | This study |
| 33. Off Thio, S Province, New Caledonia | -21.5333 | 166.3667 | MNHN-IM-2013-64426 | OR910915 | | This study |
| 34. Passe d'Ounia, S Province, New Caledonia | -21.9167 | 166.9167 | MNHN-IM-2013-64507 | OR910892 | | This study |
| 35. Off Yaté, S Province, New Caledonia | -22.1 | 167.0667 | MNHN-IM-2013-64441 | OR910923 | | This study |

| | | | | | | |
|---|---|---|---|---|---|---|
| 36. Passe d'Ounia, S Province, New Caledonia | -21.9167 | 166.9167 | MNHN-IM-2013-64428 | OR910879 | | This study |
| 37. Off Thio, S Province, New Caledonia | -21.5333 | 166.35 | MNHN-IM-2013-64447 | OR910900 | | This study |
| 38. Passe d'Ounia, S Province, New Caledonia | -21.9667 | 166.95 | MNHN-IM-2013-64442 | OR910884 | | This study |
| 39. Off Thio, S Province, New Caledonia | -21.5333 | 166.35 | MNHN-IM-2013-64446 | OR910907 | | This study |
| 40. Off Thio, S Province, New Caledonia | -21.5333 | 166.35 | MNHN-IM-2013-64436 | OR910898 | | This study |
| 41. Passe d'Ounia, S Province, New Caledonia | -21.9167 | 166.9167 | MNHN-IM-2013-64437 | OR910889 | | This study |
| 42. Passe du Solitaire, S Province, New Caledonia | -21.7667 | 166.6333 | MNHN-IM-2013-64450 | OR910893 | | This study |
| 43. Off Thio, S Province, New Caledonia | -21.5333 | 166.35 | MNHN-IM-2013-64434 | OR910897 | | This study |
| 44. Passe d'Ounia, S Province, New Caledonia | -21.9667 | 166.95 | MNHN-IM-2013-64459 | OR910890 | | This study |
| 45. Passe d'Ounia, S Province, New Caledonia | -21.9167 | 166.9167 | MNHN-IM-2013-64431 | OR910885 | | This study |

| | | | | | | | | | | |
|---|---|---|---|---|---|---|---|---|---|---|
| | | 46. Passe d'Ounia, S Province, New Caledonia | -21.9667 | 166.95 | MNHN-IM-2013-64504 | OR910882 | | | | This study |
| | | 47. Passe d'Ounia, S Province, New Caledonia | -21.9667 | 166.95 | MNHN-IM-2013-64435 | OR910888 | | | | This study |
| | | 48. Off Thio, S Province, New Caledonia | -21.5333 | 166.3667 | MNHN-IM-2013-64423 | OR910912 | | | | This study |
| | | 49. Passe d'Ounia, S Province, New Caledonia | -21.9167 | 166.9167 | MNHN-IM-2013-64505 | OR910881 | | | | This study |
| 29 | *Barneystrombus kleckhamae* (Cernohorsky, 1971) | 1. New Ireland, Papua New Guinea | -2.7167 | 150.6 | MNHN-IM-2013-58731 | OR910714 | OR934079 | OR934215 | | This study |
| | | **2. New Ireland, Papua New Guinea** | **-2.7167** | **150.6** | **MNHN-IM-2013-58594** | **OR910713** | **OR934078** | **OR934214** | **OR933974** | **This study** |
| | | 3. New Ireland, Papua New Guinea | -2.58 | 150.715 | MNHN-IM-2013-59098 | OR910715 | | | | This study |
| 30 | *Euprotomus aurisdianae* (Linnaeus, 1758) | **1. Kavieng Lagoon, Papua New Guinea** | **-2.66** | **150.665** | **MNHN-IM-2013-53759** | **OR911227** | **OR934401** | **OR934452** | **OR934352** | **This study** |
| | | 2. Ngeruktabel Is., Palau | – | – | NV | DQ525234 | | | | Latiolais et al. (2006) |
| | | 3. Barag Is., Rempi Area, Papua New Guinea | -5.0183 | 145.7983 | MNHN-IM-2013-13445 | | OR934115 | OR934245 | OR933999 | This study |
| | | 4. Kranket Is., Papua New Guinea | -5.2 | 145.8133 | MNHN-IM-2013-14905 | | OR934116 | OR934246 | OR934000 | This study |
| 31 | *Euprotomus aratrum* (Röding, 1798) | **1. Carpark 7, Changi, Singapore** | – | – | **ZRC MOL.14908** | **OR910792** | **OR934114** | **OR934244** | | **This study** |

| | | | | | | | | | | | |
|---|---|---|---|---|---|---|---|---|---|---|---|
| 32 | *Euprotomus bulla* (Röding, 1798) | 1. W sector Lavanono, S Madagascar | -25.39 | 44.8617 | MNHN-IM-2009-15859 | OR910793 | | | | This study | Formerly *Euprotomus aurora* Kronenberg, 2002, a synonym of *Euprotomus bulla* |
| | | 2. In front of Narendry Bay, Madagascar | -14.5317 | 47.4423 | MNHN-IM-2007-36709 | OR910794 | | | | This study | |
| | | 8. SW Sainte Marie Cape, Madagascar | -25.6533 | 44.885 | MNHN-IM-2009-15858 | OR910795 | | | | This study | |
| | | 3. Tome, New Ireland, Papua New Guinea | -2.7017 | 150.88 | MNHN-IM-2013-47457 | OR910796 | OR934117 | OR934247 | OR934001 | This study | |
| | | 4. Doljo Point, Panglao Is., Philippines | 9.5917 | 123.7217 | MNHN-IM-2007-33336 | OR910797 | | | | This study | |
| | | 5. Vanuatu | – | – | MNHN-IM-2007-33359 | OR910798 | | | | This study | |
| | | 6. Wékésa Is., Bruat Channel, Vanuatu | -15.6133 | 167.1417 | MNHN-IM-2007-33356 | OR910799 | | | | This study | |
| | | **7. Paquitequete, Pemba, Mozambique** | – | – | **NHMUK 20060411** | **OR911228** | **OR934402** | **OR934453** | **OR934353** | **This study** | |
| 33 | *Thetystrombus latus* (Gmelin, 1791) | **1. Bay of Goree, Thiwa, Dakar, Senegal** | **14.6117** | **-17.4483** | **MNHN-IM-2013-50048** | **OR911245** | **OR934419** | **OR934470** | **OR934370** | **This study** | |
| | | 2. Anse Bernard, Dakar, Senegal | 14.6617 | -17.4333 | MNHN-IM-2013-50101 | OR910983 | OR934188 | OR934325 | OR934059 | This study | |
| | | 3. Sao Tiago Is., Cabo Verde | – | – | NV | DQ525224 | | | | Latiolais et al. (2006) | |
| 34 | *Persististrombus granulatus* (Swainson, 1822) | 1. Pacific Panama | – | – | NV | DQ525223 | | | | Latiolais et al. (2006) | |
| | | **2. Gulf of Panama** | – | – | **UF 372340** | **OR910873** | **OR934166** | **OR934302** | **OR934043** | **This study** | |
| 35 | *Strombus gracilior* G. B. Sowerby I, 1825 | **1. Gulf of Panama** | – | – | **UF 372341** | **OR910874** | **OR934165** | **OR934301** | | **This study** | |
| | | 2. Pacific Panama | – | – | NV | DQ525209 | | | | Latiolais et al. (2006) | |

| | | | | | | | | | | | |
|---|---|---|---|---|---|---|---|---|---|---|---|
| 36 | *Strombus pugilis* Linnaeus, 1758 | 1. SW Horseshoe Beach, Florida, USA | 29.331 | -83.381 | UF 512221 | OR911249 | OR934423 | OR934474 | OR934374 | This study | Formerly *Strombus alatus* Gmelin, 1791, a synonym of *Strombus pugilis* |
| | | 2. Long Key, Florida, USA | 24.7463 | -80.8333 | UF 367475 | OR910953 | OR934175 | OR934311 | OR934049 | This study | |
| | | 3. Cedar Key, Florida, USA | – | – | NV | DQ525208 | | | | Latiolais et al. (2006) | |
| | | 4. Gulf Stream off Fort Pierce, Florida, USA | 27.4549 | -79.9532 | USNM 1448196 | MW124466 | | | | Pappalardo et al. (2021) | |
| | | 5. E Madame Is., Robert Bay, Martinique | 14.6717 | -60.88 | MNHN-IM-2013-71192 | OR910960 | | | | This study | |
| | | 6. Gris-gris Point, Port-Louis, Guadeloupe | 16.3928 | -61.5228 | MNHN-IM-2013-19521 | OR910957 | | OR934318 | | This study | |
| | | 7. E Madame Is., Robert Bay, Martinique | 14.6717 | -60.88 | MNHN-IM-2013-71190 | MW244819 | MW244819 | MW244819 | MW244819 | Irwin et al. (2021) | |
| | | 8. Coconut Bay, Royale Is., French Guiana | 5.285 | -52.5867 | MNHN-IM-2013-56947 | OR910955 | OR934181 | OR934319 | OR934051 | This study | |
| | | 9. Coconut Bay, Royale Is., French Guiana | 5.285 | -52.5867 | MNHN-IM-2013-57193 | OR910958 | | | | This study | |
| | | 10. French Guiana | 5.6483 | -52.4983 | MNHN-IM-2013-56593 | OR910959 | | | | This study | |
| | | 11. Gris-gris Point, Port-Louis, Guadeloupe | 16.3928 | -61.5228 | MNHN-IM-2013-19581 | OR910956 | | OR934317 | | This study | |

| | | | | | | | | | | |
|---|---|---|---|---|---|---|---|---|---|---|
| | | **12. Gris-gris Point, Port-Louis, Guadeloupe** | **16.3928** | **-61.5228** | **MNHN-IM-2013-19580** | **OR910954** | **OR934180** | **OR934316** | **OR934050** | **This study** |
| | | 13. Bocas Del Toro, Panama | – | – | NV | DQ525207 | | | | Latiolais et al. (2006) |
| 37 | *Titanostrombus galeatus* (Swainson, 1823) | **1. Isleta de Punta Cocos, Pearl Is., Panama** | **–** | **–** | **UF 359680** | **OR910986** | **OR934142** | **OR934272** | **OR934061** | **This study** |
| | | 2. Pacific Panama | – | – | NV | DQ525220 | | | | Latiolais et al. (2006) |
| 38 | *Lobatus raninus* (Gmelin, 1791) | 1. French Guiana | 5.77 | -52.5367 | MNHN-IM-2013-56590 | OR910844 | | | | This study |
| | | 2. Le Prêcheur, Les Abymes, Martinique | 14.8083 | -61.2267 | MNHN-IM-2013-71817 | OR910845 | | | | This study |
| | | 3. Gulf Stream off Fort Pierce, Florida, USA | 27.4645 | -79.9513 | USNM 1450320 | MW124542 | | | | Pappalardo et al. (2021) |
| | | 4. Gulf Stream off Fort Pierce, Florida, USA | 27.4224 | -79.9359 | USNM 1450665 | MW124560 | | | | Pappalardo et al. (2021) |
| | | **5. Borgnèse Point, Le Marin, Martinique** | **14.45** | **-60.8967** | **MNHN-IM-2013-72315** | **OR911237** | **OR934411** | **OR934462** | **OR934362** | **This study** |
| | | 6. French Guiana | 6.4383 | -52.43 | MNHN-IM-2013-56557 | OR910843 | OR934145 | OR934275 | OR934031 | This study |
| | | 7. Exuma Sound, Bahamas | – | – | NV | DQ525226 | | | | Latiolais et al. (2006) |
| | | 8. Vétiver, Martinique | 14.6333 | -61.13 | MNHN-IM-2013-71659 | OR910846 | | | | This study |
| 39 | *Aliger gallus* (Linnaeus, 1758) | 1. Port-Louis, Guadeloupe | 16.4463 | -61.5402 | MNHN-IM-2013-19582 | OR910697 | | OR934203 | | This study |

| | | | | | | | | | | |
|---|---|---|---|---|---|---|---|---|---|---|
| | | 2. Port-Louis, Guadeloupe | 16.3877 | -61.5298 | MNHN-IM-2013-19554 | OR910698 | | | | This study |
| | | **3. N Grande-Terre, Guadeloupe** | **16.6433** | **-61.5267** | **MNHN-IM-2013-60264** | **OR910695** | **OR934072** | **OR934202** | **OR933968** | **This study** |
| | | 4. N Grande-Terre, Guadeloupe | 16.5033 | -61.5233 | MNHN-IM-2013-60402 | OR910699 | | | | This study |
| | | 5. Bay Fort-de-France, Martinique | 14.585 | -61.0533 | MNHN-IM-2013-73168 | OR910700 | | | | This study |
| | | 6. Bay Fort-de-France, Martinique | 14.5747 | -61.0831 | MNHN-IM-2013-72989 | OR910696 | OR934132 | OR934262 | OR933969 | This study |
| 40 | *Aliger costatus* (Gmelin, 1791) | 1. E Loup Bordelais, Le Robert, Martinique | 14.7167 | -60.8467 | MNHN-IM-2013-72614 | OR910849 | OR934155 | OR934291 | OR934034 | This study |
| | | 2. Exuma Sound, Bahamas | – | – | NV | DQ525225 | | | | Latiolais et al. (2006) |
| | | **3. Trou à l'orage, Guadeloupe** | **16.3813** | **-61.5238** | **MNHN-IM-2013-19512** | **OR910848** | **OR934154** | **OR934285** | **OR934033** | **This study** |
| | | 4. Babin Cove (Morne-à-l'eau), Guadeloupe | 16.3408 | -61.5258 | MNHN-IM-2013-19552 | OR910851 | | OR934286 | | This study |
| | | 5. Babin Cove (Morne-à-l'eau), Guadeloupe | 16.3408 | -61.5258 | MNHN-IM-2013-19545 | OR910852 | | OR934287 | | This study |
| | | 6. Babin Cove (Morne-à-l'eau), Guadeloupe | 16.3408 | -61.5258 | MNHN-IM-2013-19544 | OR910853 | | OR934288 | | This study |
| | | 7. Babin Cove (Morne-à-l'eau), Guadeloupe | 16.3408 | -61.5258 | MNHN-IM-2013-19539 | OR910850 | | OR934289 | | This study |
| | | 8. Babin Cove (Morne-à-l'eau), Guadeloupe | 16.3408 | -61.5258 | MNHN-IM-2013-19515 | OR910854 | | OR934290 | | This study |
| | | 9. E Madame Is., Robert Bay, Martinique | 14.6717 | -60.88 | MNHN-IM-2013-71228 | OR910855 | | | | This study |

| | | | | | | | | | | |
|---|---|---|---|---|---|---|---|---|---|---|
| | | 10. Passe SE Fond Blanc, Le François, Martinique | 14.6333 | -60.8533 | MNHN-IM-2013-71016 | OR910856 | | | | This study |
| | | 11. Passe SE Fond Blanc, Le François, Martinique | 14.6333 | -60.8533 | MNHN-IM-2013-71015 | OR910847 | OR934127 | OR934257 | OR934032 | This study |
| 41 | *Aliger gigas* (Linnaeus, 1758) | 1. Trou à l'orage, Guadeloupe | 16.3813 | -61.5238 | MNHN-IM-2013-19564 | OR910705 | | OR934205 | | This study |
| | | 2. Gris-gris Point, Port-Louis, Guadeloupe | 16.3928 | -61.5228 | MNHN-IM-2013-19563 | OR910708 | | OR934210 | | This study |
| | | 3. Lagoon Petite Terre, Guadeloupe | 16.1758 | -61.1112 | MNHN-IM-2013-8735 | OR910707 | | OR934207 | | This study |
| | | 4. Trou à l'orage, Guadeloupe | 16.3813 | -61.5238 | MNHN-IM-2013-19514 | OR910704 | | OR934209 | | This study |
| | | 5. SE Pimantee Point, Trois Rivières, Martinique | 14.4633 | -60.965 | MNHN-IM-2013-71960 | OR910709 | | | | This study |
| | | 6. Berry Is., Bahamas | – | – | NV | DQ525222 | | | | Latiolais et al. (2006) |
| | | 7. Lagoon Petite Terre, Guadeloupe | 16.1758 | -61.1112 | MNHN-IM-2013-8734 | OR910706 | | OR934206 | | This study |
| | | 8. Port-Louis, Guadeloupe | 16.4167 | -61.55 | MNHN-IM-2013-20183 | OR910701 | OR934073 | OR934204 | OR933970 | This study |
| | | 9. San Andrés Archipelago, Colombia | – | – | Sg300-UNAL-SAA | NC_024932 | NC_024932 | NC_024932 | | Márquez et al. (2014) |
| | | 10. Mexico | – | – | ECOCHM 1382 | KU317712 | | | | Martínez-Arce et al. (2015) |

| | | | | | | | | | | |
|---|---|---|---|---|---|---|---|---|---|---|
| | | 11. Mexico | – | – | ECOCHM 1378 | KU317714 | | | | Martínez-Arce et al. (2015) |
| | | 12. Mexico | – | – | ECOCHM 1380 | KU317713 | | | | Martínez-Arce et al. (2015) |
| | | 13. Mexico | – | – | ECOCHM 1377 | KU317715 | | | | Martínez-Arce et al. (2015) |
| | | 14. Cozumel Is. Protected Area, Mexico | – | – | NV | MZ157283 | MZ157283 | MZ157283 | | Machkour-M'Rabet et al. (2021) |
| | | 15. Gulf Stream off Fort Pierce, Florida, USA | 27.4549 | -79.9532 | USNM 1448199 | MW124469 | | | | Pappalardo et al. (2021) |
| | | 16. Diamond Rock, Martinique | 14.445 | -61.0367 | MNHN-IM-2013-71682 | OR910710 | | | | This study |
| | | 17. N Fajou Island, Guadeloupe | 16.3542 | -61.5847 | MNHN-IM-2013-19536 | OR910703 | | OR934208 | | This study |
| | | **18. N Fajou Island, Guadeloupe** | **16.3542** | **-61.5847** | **MNHN-IM-2013-19530** | **OR910702** | **OR934133** | **OR934263** | **OR933971** | **This study** |
| | | 19. S Port Louis, Guadeloupe | 16.3814 | -61.5239 | MNHN-IM-2013-19509 | | | OR934211 | | This study |
| 42 | *Gibberulus gibberulus* (Linnaeus, 1758) | 1. Inhaca Is., Maputo Bay, Mozambique | -26.0733 | 32.9517 | MNHN-IM-2013-64490 | OR910803 | | | | This study |
| | | **2. Inhaca Is., Maputo Bay, Mozambique** | **-26.0733** | **32.9517** | **MNHN-IM-2013-64491** | **OR911229** | **OR934403** | **OR934454** | **OR934354** | **This study** |
| | | 3. Inhaca Is., Maputo Bay, Mozambique | -26.0733 | 32.9517 | MNHN-IM-2013-64488 | OR910801 | | | | This study |

| | | | | | | | | | | |
|---|---|---|---|---|---|---|---|---|---|---|
| | | 4. Inhaca Is., Maputo Bay, Mozambique | -26.0733 | 32.9517 | MNHN-IM-2013-64485 | OR910800 | OR934118 | OR934248 | OR934002 | This study |
| | | 5. Inhaca Is., Maputo Bay, Mozambique | -26.065 | 32.955 | MNHN-IM-2013-64493 | OR910802 | | | | This study |
| 43 | *Gibberulus gibbosus* (Röding, 1798) | 1. Raivavae, French Polynesia | -23.8704 | -147.699 | MNHN-IM-2013-48344 | OR910806 | | | | This study |
| | | 2. Raivavae, French Polynesia | -23.8704 | -147.699 | MNHN-IM-2013-48341 | OR910804 | OR934120 | OR934250 | OR934003 | This study |
| | | 3. Mo'orea, French Polynesia | – | – | BMOO 03706 | KC706865 | | | | Leray et al. (2013) |
| | | 4. Kabira, Ishigaki Is., Okinawa, Japan | 24.4833 | 124.1167 | NHMUK 2019042 | OR910805 | | | | This study |
| | | 5. Guam | – | – | NV | DQ525211 | | | | Latiolais et al. (2006) |
| | | **6. Kavieng Lagoon, Papua New Guinea** | **-2.625** | **150.775** | **MNHN-IM-2013-46767** | **OR911230** | **OR934404** | **OR934455** | **OR934355** | **This study** |
| | | 7. Alexishafen, Papua New Guinea | -5.0883 | 145.8017 | MNHN-IM-2013-18156 | | OR934119 | OR934249 | OR934004 | This study |
| | | 8. Port Vila, Vanuatu | – | – | MNHN-IM-2013-64499 | | OR934121 | OR934251 | OR934005 | This study |
| 44 | *Conomurex fasciatus* (Born, 1778) | **1. Farasan Is., Abu Lad, Saudi Arabia** | **16.7977** | **42.1992** | **UF 463278** | **OR911213** | **OR934387** | **OR934437** | **OR934339** | **This study** |
| | | 2. Farasan Is., Abu Lad, Saudi Arabia | 16.7977 | 42.1992 | UF 463647 | OR910773 | OR934086 | OR934220 | OR933987 | This study |
| 45 | *Conomurex persicus* (Swainson, 1821) | **1. Asterias Hotel, Ayia Napa Marina, Cyprus** | **34.9816** | **33.9536** | **NHMUK 20230931** | **OR911220** | **OR934393** | **OR934444** | **OR934345** | **This study** |
| 46 | | **1. Monseigneur Beach, S Madagascar** | **-25.035** | **46.9983** | **MNHN-IM-2009-15860** | **OR911211** | **OR934385** | **OR934435** | **OR934337** | **This study** |

| | | | | | | | | | | |
|---|---|---|---|---|---|---|---|---|---|---|
| | *Conomurex decorus* (Röding, 1798) | 2. Inhaca Is., Maputo Bay, Mozambique | -26.105 | 32.9667 | MNHN-IM-2013-64487 | OR910768 | | | | This study |
| | | 3. Inhaca Is., Maputo Bay, Mozambique | -26.1817 | 32.9533 | MNHN-IM-2013-64486 | OR910771 | | | | This study |
| | | 4. N Pemba Bay headland, Mozambique | – | – | NHMUK 20060447 | OR910772 | | | | This study |
| | | 5. Inhaca Is., Maputo Bay, Mozambique | -26.065 | 32.955 | MNHN-IM-2013-64494 | OR910767 | OR934080 | OR934216 | | This study |
| | | 6. Inhaca Is., Maputo Bay, Mozambique | -26.0217 | 32.8983 | MNHN-IM-2013-64497 | OR910769 | | | | This study |
| | | 7. Inhaca Is., Maputo Bay, Mozambique | -26.0217 | 32.8983 | MNHN-IM-2013-64495 | OR910770 | | | | This study |
| 47 | *Conomurex luhuanus* (Linnaeus, 1758) | 1. W Ratua Is., S Aoré Is., Vanuatu | -15.61 | 167.1683 | MNHN-IM-2007-33364 | OR910776 | OR934092 | OR934225 | | This study |
| | | 2. Kranket Is., Papua New Guinea | -5.2017 | 145.8217 | MNHN-IM-2013-10140 | OR910775 | OR934091 | OR934224 | | This study |
| | | 3. Heron Is., Queensland, Australia | – | – | AM C203214 | AY296831 | DQ916425 | | | Colgan et al. (2003, 2007) |
| | | 4. Hainan Province, China | – | – | NV | NC_035726 | NC_035726 | NC_035726 | | Zhao et al. (2018) |
| | | 5. Wenchang, Hainan Province, China | – | – | OUC SLWC02 | JF693429 | | | | Sun et al. (2012) |
| | | 6. Wenchang, Hainan Province, China | – | – | OUC SLWC01 | JF693430 | | | | Sun et al. (2012) |
| | | 7. Wenchang, Hainan Province, China | – | – | OUC SLWC03 | JF693431 | | | | Sun et al. (2012) |
| | | 8. Wenchang, Hainan Province, China | – | – | OUC SLWC04 | JF693432 | | | | Sun et al. (2012) |

| | | | | | | | | | | | |
|---|---|---|---|---|---|---|---|---|---|---|---|
| | | 9. Xincun, Lingshui, Hainan Province, China | – | – | OUC G222 | MN389059 | | | | Ran et al. (2020) | |
| | | 10. Biking, Pangalo Is., Philippines | 9.5883 | 123.8417 | MNHN-IM-2007-33343 | OR910777 | | | | This study | |
| | | 11. W Ratua Is., S Aoré Is., Vanuatu | -15.61 | 167.1683 | MNHN-IM-2007-33379 | OR910778 | | | | This study | |
| | | **12. Fusaki, Okinawa, Ishigaki Is., Japan** | **24.365** | **124.11** | **NHMUK 2019039** | **OR911216** | **OR934390** | **OR934440** | **OR934342** | **This study** | |
| | | 13. Amani-U-Shima, Japan | – | – | NV | DQ525231 | | | | Latiolais et al. (2006) | |
| | | 14. Cape Bise, Motobu, Okinawa Is., Japan | 26.71 | 127.8767 | NHMUK 2019040 | OR910774 | OR934090 | OR934223 | OR933988 | This study | |
| | | 15. W Ratua Is., S Aoré Is., Vanuatu | -15.61 | 167.1683 | MNHN-IM-2007-33376 | OR910779 | | | | This study | |
| | | 16. Wenchang, Hainan Province, China | – | – | OUC G223 | MN389060 | | | | Ran et al. (2020) | |
| | | 17. NL | – | – | NV | | | AF174212 | | Duda et al. (2001) | |
| 48 | *Lentigo pipus* (Röding, 1798) | **1. Kavieng Lagoon, Papua New Guinea** | **-2.6933** | **150.6483** | **MNHN-IM-2013-51774** | **OR911236** | **OR934410** | **OR934461** | **OR934361** | **This study** | |
| | | 2. S Faux-Cap, S Madagascar | -25.9082 | 45.5533 | MNHN-IM-2009-15862 | OR910842 | OR934144 | OR934274 | OR934030 | This study | |
| | | 3. Koki, N Kyoda IC, Nago, Okinawa, Japan | 26.545 | 127.46 | NHMUK 2019051 | | OR934143 | OR934273 | | This study | |
| 49 | *Lentigo thersites* (Swainson, 1823) comb. nov. | **1. Banc Ouest, S Province, New Caledonia** | **-22.446** | **166.483** | **UF 510153** | **OR911255** | **OR934429** | **OR934480** | **OR934380** | **This study** | Formerly *Thersistrombus thersites* (Swainson, 1823) |

| | | | | | | | | | | |
|---|---|---|---|---|---|---|---|---|---|---|
| 50 | *Lentigo lentiginosus* (Linnaeus, 1758) | **1. Kavieng Lagoon, Papua New Guinea** | **-2.66** | **150.665** | **MNHN-IM-2013-53752** | **OR911235** | **OR934409** | **OR934460** | **OR934360** | **This study** |
| | | 2. Kavieng Lagoon,Papua New Guinea | -2.66 | 150.665 | MNHN-IM-2013-53756 | OR910841 | OR934137 | OR934267 | OR934027 | This study |
| | | 3. Lingshui, Hainan Province, China | – | – | OUC SLLS01 | JF693421 | | | | Sun et al. (2012) |
| | | 4. Xishaqundao, Hainan Province, China | – | – | OUC SLXSQD02 | JF693422 | | | | Sun et al. (2012) |
| | | 5. Fusaki, Okinawa, Ishigaki Is., Japan | 24.365 | 124.11 | NHMUK 2019050 | | OR934136 | OR934266 | OR934028 | This study |
| | | 6. Sesoko Is., Okinawa, Japan | 26.65 | 127.875 | NHMUK 2019049 | | OR934138 | OR934268 | OR934029 | This study |
| 51 | *Latissistrombus sinuatus* ([Lightfoot], 1786) | **1. Kranket Is., Papua New Guinea** | **-5.1885** | **145.8237** | **MNHN-IM-2013-15351** | **OR910839** | **OR934182** | **OR934320** | **OR934026** | **This study** |
| | | 2. Mindanao, Selinog Is., Philippines | – | – | NV | DQ525229 | | | | Latiolais et al. (2006) |
| 52 | *Latissistrombus latissimus* (Linnaeus, 1758) | **1. SE off Isle of Pines, New Caledonia** | **-23.4198** | **168.075** | **MNHN-IM-2013-63890** | **OR910837** | **OR934177** | **OR934313** | **OR934024** | **This study** |
| | | 2. N Port Benier Bay, E Aoré Is., Vanuatu | -15.5631 | 167.2075 | MNHN-IM-2007-33388 | OR910838 | OR934178 | OR934314 | OR934025 | This study |
| 53 | *Latissistrombus taurus* (Reeve, 1857) | 1. Hospital Point, Guam | – | – | UF 282384 | OR910840 | OR934183 | OR934321 | | This study |
| | | **2. Hospital Point., Guam** | – | – | **UF 282384** | **OR911250** | **OR934424** | **OR934475** | **OR934375** | **This study** |
| | | 3. Hospital Point, Guam | – | – | NV | DQ525228 | | | | Latiolais et al. (2006) |

| | | | | | | | | | | |
|---|---|---|---|---|---|---|---|---|---|---|
| 54 | *Ophioglossolambis digitata* (Perry, 1811) | **1. Choumare Islet, S Madagascar** | **-24.8367** | **47.115** | **MNHN-IM-2009-15861** | **OR911244** | **OR934418** | **OR934469** | **OR934369** | **This study** |
| 55 | *Tricornis tricornis* ([Lightfoot], 1786) | **1. Farasan Banks, Marca Is., Saudi Arabia** | **18.2206** | **41.3243** | **UF 478460** | **OR911256** | **OR934430** | **OR934481** | **OR934381** | **This study** |
| | | 2. Farasan Banks, Marca Is., Saudi Arabia | 18.2206 | 41.3243 | UF 478459 | OR910987 | OR934194 | OR934331 | OR934062 | This study |
| 56 | *Harpago chiragra* (Linnaeus, 1758) | 1. Kavieng Lagoon, Papua New Guinea | -2.5867 | 150.77 | MNHN-IM-2013-46488 | OR910807 | OR934122 | OR934252 | | This study |
| | | **2. Okinawa Is., Nago, N Kyoda IC, Koki, Japan** | **26.545** | **127.46** | **NHMUK 2019044** | **OR911231** | **OR934405** | **OR934456** | **OR934356** | **This study** |
| | | 3. Quanfu Is., S China Sea | – | – | NV | MH122656 | MH122656 | MH122656 | | Jiang et al. (2019) |
| | | 4. Xisha, Hainan province, China | 16.7099 | 112.3519 | TMBC 030691 | MN885884 | MN885884 | MN885884 | | Liu et al. (2020) |
| 57 | *Harpago rugosa* (G. B. Sowerby II, 1842) | **1. Great Koumac Reef, New Caledonia** | **-20.7883** | **164.28** | **MNHN-IM-2019-1739** | **OR910808** | **OR934123** | **OR934253** | **OR934006** | **This study** |
| 58 | *Lambis scorpius* (Linnaeus, 1758) | 1. Catarman, Panglao Is., Philippines | 9.61 | 123.8733 | MNHN-IM-2007-33350 | OR910830 | | | | This study |
| | | 2. Koki, N Kyoda IC, Nago, Okinawa, Japan | 26.545 | 127.46 | NV | OR910829 | OR934147 | OR934277 | OR934020 | This study |
| | | **3. E Sek Is., Papua New Guinea** | **-5.0817** | **145.8233** | **MNHN-IM-2013-18324** | **OR911239** | **OR934413** | **OR934464** | **OR934364** | **This study** |
| | | 4. Off Aimbuei Bay, E Aoré Is., Vanuatu | -15.5533 | 167.2133 | MNHN-IM-2007-33352 | OR910831 | | | | This study |
| | | 5. N Aimbuei Bay, NE Aoré Is., Vanuatu | -15.5383 | 167.2183 | MNHN-IM-2007-33370 | OR910832 | | | | This study |

| | | | | | | | | | | | |
|---|---|---|---|---|---|---|---|---|---|---|---|
| 59 | *Lambis millepeda* (Linnaeus, 1758) | **1. Kavieng Lagoon, Papua New Guinea** | **-2.635** | **150.7733** | **MNHN-IM-2013-46848** | **OR910825** | **OR934139** | **OR934269** | **OR934018** | **This study** | |
| | | 2. S Megas Islet, Papua New Guinea | -5.0883 | 145.81 | MNHN-IM-2013-13064 | OR910826 | OR934140 | OR934270 | OR934019 | This study | |
| | | 3. Catarman, Panglao Is., Philippines | 9.61 | 123.8733 | MNHN-IM-2007-33369 | OR910827 | OR934141 | OR934271 | | This study | |
| | | 4. Cocos (Keeling) Is., Australia | – | – | NV | DQ525238 | | | | Latiolais et al. (2006) | |
| 60 | *Lambis truncata* ([Lightfoot], 1786) | **1. Inhaca Is., Maputo Bay, Mozambique** | **-25.91** | **33.0483** | **MNHN-IM-2013-64484** | **OR911240** | **OR934414** | **OR934465** | **OR934365** | **This study** | |
| 61 | *Lambis sowerbyi* (Mörch, 1872) | 1. E Aoré Is., Vanuatu | -15.576 | 167.2272 | MNHN-IM-2007-33386 | OR910836 | | | | This study | Recognized at species rank; formerly *Lambis truncata sowerbyi* |
| | | **2. Great Koumac Reef, New Caledonia** | **-20.6425** | **164.1825** | **MNHN-IM-2013-84270** | **OR910835** | **OR934150** | **OR934280** | **OR934023** | **This study** | |
| | | 3. N coast New Ireland, Papua New Guinea | -2.5868 | 150.8382 | MNHN-IM-2013-55288 | OR910833 | OR934148 | OR934278 | OR934021 | This study | |
| | | 4. Planet Rock, Papua New Guinea | -5.2582 | 145.8188 | MNHN-IM-2013-11082 | OR910834 | OR934149 | OR934279 | OR934022 | This study | |
| | | 5. Quanfu Is., S China Sea | – | – | NV | MH115428 | MH115428 | MH115428 | | Jiang et al. (2019) | |
| 62 | *Lambis crocata* (Link, 1807) | **1. Kavieng Lagoon, Papua New Guinea** | **-2.6102** | **150.6769** | **MNHN-IM-2013-53676** | **OR910820** | **OR934128** | **OR934258** | **OR934015** | **This study** | |
| | | 2. Aimbuei Bay, Aoré Is., Vanuatu | – | – | MNHN-IM-2007-33382 | OR910821 | OR934129 | OR934259 | | This study | |
| | | 3. SW Mavéa Is., Vanuatu | -15.3954 | 167.2224 | MNHN-IM-2007-33387 | OR910822 | | | | This study | |
| 63 | *Lambis robusta* (Swainson, 1821) | **1. Society Is., Moorea Temae, French Polynesia** | **-17.479** | **-149.858** | **UF 427905** | **OR910828** | **OR934146** | **OR934276** | | **This study** | |

| | | | | | | | | | | | |
|---|---|---|---|---|---|---|---|---|---|---|---|
| | | 2. Society Is., Moorea Temae, French Polynesia | -17.479 | -149.7643 | UF 427893 | OR911238 | OR934412 | OR934463 | OR934363 | This study | |
| 64 | *Lambis lambis* A | **1. S Dumduman Is., Papua New Guinea** | **-5.0033** | **145.7933** | **MNHN-IM-2013-12410** | **OR911234** | **OR934408** | **OR934459** | **OR934359** | **This study** | Three putative cryptic species. True *Lambis lambis* (Linnaeus, 1758) is type species of *Lambis.* Note that the *L. lambis* cryptic species reported by Li et al. (2022b) are here identified as *L. sowerbyi* and *L. lambis* B. |
| | | 2. Nagada Harbour, Papua New Guinea | -5.1567 | 145.8 | MNHN-IM-2013-17026 | OR910824 | OR934135 | OR934265 | OR934017 | This study | |
| 65 | *Lambis lambis* B | **1. Cape Bise, Motobu, Okinawa Is., Japan** | **26.71** | **127.8767** | **NHMUK 2019046** | **OR911233** | **OR934407** | **OR934458** | **OR934358** | **This study** | |
| | | 2. Sanya, Hainan Province, China | 18.2394 | 109.3783 | HNU *L. lambis*-SY | ON840105 | ON840105 | | | Li et al. (2022) | |
| | | 3. Cape Bise, Motobu, Okinawa Is., Japan | 26.71 | 127.8767 | NHMUK 2019045 | OR910823 | OR934134 | OR934264 | OR934016 | This study | |
| | | 4. Zhaoshu Island, China | 16.9797 | 112.2719 | HNU *L. lambis*-QF | ON840106 | | | | Li et al. (2022) | |
| | | 5. Wenchang, Hainan Province, China | 19.5167 | 110.8 | OUC 21907 | HQ834110 | | HQ833978 | | Zou et al. (2011) | |
| | | 6. Sanya, Hainan Province, China | – | – | OUC LLSY01 | JF693383 | | | | Sun et al. (2012) | |
| | | 7. Sanya, Hainan Province, China | – | – | OUC LLSY02 | JF693384 | | | | Sun et al. (2012) | |
| | | 8. Sanya, Hainan Province, China | – | – | OUC LLSY03 | JF693385 | | | | Sun et al. (2012) | |
| | | 9. S lagoon, Pulau Subar Darat, Singapore | – | – | ZRC MOL.010721 | MN690208 | | | | Ip et al. (2019) | |
| 66 | *Lambis lambis* C | **1. Bay of Bengal** | **–** | **–** | **NV** | **MH908185** | | | | **Labeeb et al. (2018)** | |
| | | 2. Bay of Bengal | – | – | NV | MH908189 | | | | Labeeb et al. (2018) | |

| | | | | | | | | | | | |
|---|---|---|---|---|---|---|---|---|---|---|---|
| | | 3. Bay of Bengal | – | – | NV | MH908186 | | | | Labeeb et al. (2018) | |
| | | 4. Bay of Bengal | – | – | NV | MH908187 | | | | Labeeb et al. (2018) | |
| 67 | *"Canarium" wilsonorum* A | 1. SW Sainte Marie Cape, S Madagascar | -25.8067 | 44.86 | MNHN-IM-2009-15852 | OR910765 | | | | This study | Two putative cryptic species. Generic affinity uncertain; requires molecular confirmation with type species *Canarium wilsonorum* (R.T. Abbott, 1967) |
| | | **2. SW Sainte Marie Cape S Madagascar** | **-25.7617** | **44.8667** | **MNHN-IM-2009-15840** | **OR911223** | **OR934396** | **OR934447** | | **This study** | |
| 68 | *"Canarium" wilsonorum* B | **1. Orote Peninsula, Guam** | **–** | **–** | **NV** | **DQ525214** | | | | **Latiolais et al. (2006)** | |
| 69 | *Fusistrombus fusiformis* (G. B. Sowerby II, 1842) | **1. In front of Narendry Bay, Madagascar** | **-14.5317** | **47.4423** | **MNHN-IM-2007-36710** | **OR911214** | **OR934388** | **OR934438** | **OR934340** | **This study** | |
| | | 2. In front of Narendry Bay, Madagascar | -14.5317 | 47.4423 | MNHN-IM-2007-36647 | OR910867 | OR934087 | | | This study | |
| | | 3. Zanzibar, Tanzania | – | – | NV | DQ525212 | | | | Latiolais et al. (2006) | |
| 70 | *Hawaiistrombus scalariforme* (Duclos, 1833) comb. nov. | 1. Orote Peninsula, Guam | – | – | NV | DQ525213 | | | | Latiolais et al. (2006) | Formerly *Canarium scalariforme* (Duclos, 1833) |
| | | **2. Tubuai Is., French Polynesia** | **-23.4254** | **-149.4264** | **MNHN-IM-2013-48350** | **OR910763** | **OR934101** | **OR934233** | | **This study** | |
| | | 3. Tubuai Is., French Polynesia | -23.4254 | -149.4264 | MNHN-IM-2013-48351 | OR910764 | OR934102 | OR934234 | | This study | |
| 71 | *Dolomena epidromis* | 1. Kavieng Lagoon, Papua New Guinea | -2.5709 | 150.4939 | MNHN-IM-2013-54830 | OR910809 | OR934130 | OR934260 | OR934007 | This study | Formerly *Labiostrombus epidromis* (Linnaeus, 1758) |

| | | | | | | | | | | | |
|---|---|---|---|---|---|---|---|---|---|---|---|
| | (Linnaeus, 1758) comb. nov. | **2. SW Pandop Point, New Caledonia** | **-20.595** | **164.2533** | **MNHN-IM-2019-1733** | **OR910810** | **OR934131** | **OR934261** | **OR934008** | **This study** | |
| 72 | *Dolomena vittata* (Linnaeus, 1758) comb. nov. | 1. Shoal Bay, Darwin Harbour, Australia | -12.25 | 130.8333 | NTM P015366 | OR910791 | OR934113 | OR934243 | | This study | Formerly *Doxander operosus* (Röding, 1798), a synonym of *Doxander vittatus* (type species of *Doxander*), now *Dolomena vittata.* |
| | | **2. St John's Is., Singapore** | – | – | **ZRC MOL.7536** | **OR911224** | **OR934397** | **OR934448** | **OR934348** | **This study** | |
| | | 3. Weizhoudao, Guangxi Province, China | – | – | OUC SVWZD01 | JF693433 | | | | Sun et al. (2012) | |
| | | 4. Xincun, Lingshui, Hainan Province, China | – | – | OUC G45 | MN389057 | | | | Ran et al. (2020) | |
| | | 5. Xincun, Lingshui, Hainan Province, China | – | – | OUC G46 | MN389058 | | | | Ran et al. (2020) | |
| | | 6. Shenchong, Hainan Province, China | – | – | OUC G43 | MN389055 | | | | Ran et al. (2020) | |
| | | 7. Shenchong, Hainan Province, China | – | – | OUC G44 | MN389056 | | | | Ran et al. (2020) | |
| | | 8. Beihai, Guangxi Province, China | – | – | OUC SVWZD02 | JF693434 | | | | Sun et al. (2012) | |
| | | 9. Beihai, Guangxi Province, China | – | – | OUC SVWZD03 | JF693435 | | | | Sun et al. (2012) | |
| 73 | *Dolomena labiosa* (W. Wood, 1828) | 1. Cortes, Bohol Is., Philippines | 9.7033 | 123.8433 | MNHN-IM-2007-33347 | OR910780 | OR934069 | OR934199 | OR933989 | This study | Formerly *Dolomena abbotti* Dekkers & Liverani, 2011, a synonym of *Dolomena labiosa* |
| | | 2. E Baler Bay, Philippines | 15.85 | 121.5783 | MNHN-IM-2007-34491 | OR910781 | OR934070 | OR934200 | OR933990 | This study | |
| | | 3. E Baler Bay, Philippines | 15.85 | 121.5783 | MNHN-IM-2007-34495 | OR910782 | OR934071 | OR934201 | OR933991 | This study | |

| | | | | | | | | | | | |
|---|---|---|---|---|---|---|---|---|---|---|---|
| | | 4. In front of Narendry Bay, Madagascar | -14.4833 | 47.4333 | MNHN-IM-2007-36618 | OR910785 | OR934110 | OR934240 | OR933994 | This study | |
| | | **5. NW Majunga, Madagascar** | **-15.4** | **45.9333** | **MNHN-IM-2007-36711** | **OR911225** | **OR934398** | **OR934449** | **OR934349** | **This study** | |
| | | 6. In front of Narendry Bay, Madagascar | -14.5367 | 47.4233 | MNHN-IM-2007-36651 | OR910787 | | | | This study | |
| | | 7. N Saint-André Cape, Madagascar | -15.7917 | 44.75 | MNHN-IM-2007-38201 | OR910786 | | | OR933996 | This study | |
| | | 8. N Saint-André Cape, Madagascar | -15.7917 | 44.75 | MNHN-IM-2007-38127 | OR910788 | | | | This study | |
| | | 9. Betw. Nosy Be and Nosy Tanikely, Madagascar | – | – | UF 423433 | | OR934107 | OR934238 | OR933995 | This study | |
| 74 | *Dolomena robusta* (G. B. Sowerby III, 1875) comb. nov. | 1. Cyrene Reef, Singapore | – | – | ZRC MOL.14909 | OR910866 | OR934157 | OR934293 | | This study | Formerly *Neodilatilabrum robustum* (G. B. Sowerby III, 1875) |
| | | 2. Beihai, Guangxi Province, China | – | – | OUC SMRBH05 | JF693437 | | | | Sun et al. (2012) | |
| | | 3. Beihai, Guangxi Province, China | – | – | OUC SMRBH03 | JF693428 | | | | Sun et al. (2012) | |
| | | 4. Beihai, Guangxi Province, China | – | – | OUC SMRBH04 | JF693436 | | | | Sun et al. (2012) | |
| | | **5. Changi, Singapore** | **1.3095** | **103.9712** | **ZRC MOL.2897** | **OR911242** | **OR934416** | **OR934467** | **OR934367** | **This study** | |
| | | 6. Beihai, Guangxi Province, China | – | – | OUC SMRBH01 | JF693426 | | | | Sun et al. (2012) | |
| | | 7. Beihai, Guangxi Province, China | – | – | OUC SMRBH02 | JF693427 | | | | Sun et al. (2012) | |
| 75 | *Dolomena variabilis* | **1. Plateau Karembé (Pte Nord), New Caledonia** | **-20.6267** | **164.2883** | **MNHN-IM-2019-1736** | **MW244824** | **MW244824** | **MW244824** | **MW244824** | **Irwin et al. (2021)** | Formerly *Ministrombus variabilis* (Swainson, 1820) |

| | | | | | | | | | | |
|---|---|---|---|---|---|---|---|---|---|---|
| | (Swainson, 1820) comb. nov. | 2. Kavieng Lagoon, Papua New Guinea | -2.605 | 150.77 | MNHN-IM-2013-46526 | OR910863 | OR934158 | OR934294 | OR934038 | This study |
| | | 3. Ogtong Beach, Bantayan Is., Philippines | – | – | NHMUK 20040774 | OR910865 | | | | This study |
| | | 4. Cyrene Reef, Singapore | – | – | ZRC MOL.3309 | OR910864 | OR934159 | OR934295 | | This study |
| 76 | *Dolomena pulchella* (Reeve, 1851) | **1. N Kavieng Harbour, Papua New Guinea** | **-2.5783** | **150.7867** | **MNHN-IM-2013-47346** | **OR911226** | **OR934399** | **OR934450** | **OR934350** | **This study** |
| | | 2. S Kranket Is., Papua New Guinea | -5.205 | 145.8133 | MNHN-IM-2013-17852 | OR910789 | OR934112 | OR934242 | OR933997 | This study |
| | | 3. W Ratua Is., S Aoré Is., Turtle Bay, Vanuatu | -15.3567 | 167.2133 | MNHN-IM-2007-33358 | OR910790 | | | | This study |
| 77 | *Ministrombus minimus* (Linnaeus, 1771) | **1. S Yabob Is., Papua New Guinea** | **-5.2583** | **145.7883** | **MNHN-IM-2013-17612** | **OR911241** | **OR934415** | **OR934466** | **OR934366** | **This study** |
| | | 2. Tadwai Is., Papua New Guinea | -4.9883 | 145.7917 | MNHN-IM-2013-14458 | OR910860 | OR934111 | OR934241 | OR934037 | This study |
| | | 3. S Palikulo Bay, Vanuatu | -15.4933 | 167.2483 | MNHN-IM-2007-33333 | OR910861 | OR934156 | OR934292 | | This study |
| | | 4. S Palikulo Bay, Vanuatu | -15.4933 | 167.2483 | MNHN-IM-2007-33354 | OR910862 | | | | **This study** |
| 78 | *Dolomena swainsoni* (Reeve, 1850) | **1. Xincun, Lingshui, Hainan Province, China** | **–** | **–** | **OUC G129** | **MN389053** | | | | **Ran et al. (2020)** |
| | | 2. Xincun, Lingshui, Hainan Province, China | – | – | OUC G130 | MN389054 | | | | Ran et al. (2020) |

| | | | | | | | | | | | |
|---|---|---|---|---|---|---|---|---|---|---|---|
| 79 | *Dolomena dilatata* (Swainson, 1821) | 1. Grand Pass, N Province, New Caledonia | -19.88 | 163.83 | MNHN-IM-2007-34735 | OR910783 | OR934108 | OR934239 | OR933992 | This study | *Dolomena dilatata* (Swainson, 1821) |
| | | **2. Grand Pass, N Province, New Caledonia** | **-19.83** | **163.795** | **MNHN-IM-2007-34736** | **OR910784** | **OR934164** | **OR934300** | **OR933993** | **This study** | |
| 80 | *Dolomena turturella* A comb. nov. | 1. Bantayan, Cebu, Philippines | – | – | NV | DQ525210 | | | | Latiolais et al. (2006) | Formerly *Laevistrombus turturella* (Röding, 1798) |
| | | 2. Yagaji Is. Harbour, Nago, Okinawa, Japan | – | – | NHMUK 2019052 | OR910814 | OR934152 | OR934283 | OR934012 | This study | |
| | | **3. Yagaji Is. Harbour, Nago, Okinawa, Japan** | – | – | **NHMUK 2019053** | **OR910815** | **OR934153** | **OR934284** | **OR934013** | **This study** | |
| 81 | *Dolomena turturella* B comb. nov. | **1. Reef off Pulau Tekong, E Johor Strait, Singapore** | **1.4367** | **104.0483** | **ZRC MOL.28354** | **OR911232** | **OR934406** | **OR934457** | **OR934357** | **This study** | Formerly *Laevistrombus turturella* (Röding, 1798) |
| | | 2. Penghu Is., Taiwan | 23.5 | 119.5 | NTOU LC-01-2020 | NC_053786 | NC_053786 | NC_053786 | | Lee et al. (2021) | |
| | | 3. SW Pulau Semakau, Singapore | 1.1976 | 103.7582 | ZRC MOL.2917 | | | OR934282 | | This study | |
| 82 | *Dolomena taeniata* (Quoy & Gaimard, 1834) comb. nov. | 1. Kavieng Lagoon, Papua New Guinea | -2.5709 | 150.4939 | MNHN-IM-2013-54834 | OR910811 | OR934124 | OR934254 | OR934009 | This study | Formerly *Laevistrombus taeniatus* (Quoy & Gaimard, 1834) |
| | | 2. Kavieng Lagoon, Papua New Guinea | -2.5709 | 150.4939 | MNHN-IM-2013-54839 | OR910812 | OR934125 | OR934255 | OR934010 | This study | |
| | | **3. Duadnatun Is., N Riwo, Papua New Guinea** | **-5.2025** | **145.8218** | **MNHN-IM-2013-17251** | **OR910813** | **OR934151** | **OR934281** | **OR934011** | **This study** | |
| 83 | *Dolomena vanikorensis* | **1. Near Paagoumène, New Caledonia** | **-20.4833** | **164.175** | **MNHN-IM-2019-1740** | **OR910816** | **OR934126** | **OR934256** | **OR934014** | **This study** | |

| | | | | | | | | | | | |
|---|---|---|---|---|---|---|---|---|---|---|---|
| | (Quoy & Gaimard, 1834) comb. nov. | 2. Belmoul Lagoon, Vanuatu | -15.5967 | 167.1017 | MNHN-IM-2009-11060 | HQ401579 | HQ401625 | HQ401689 | | Puillandre et al. (2011) | Formerly *Laevistrombus vanikorensis* (Quoy & Gaimard, 1834) |
| | | 3. Aisari Bay, E Aoré Is., Vanuatu | -15.571 | 167.1985 | MNHN-IM-2007-33377 | OR910819 | | | | This study | |
| | | 4. Maritime College, Segond Channel, Vanuatu | -15.5226 | 167.1659 | MNHN-IM-2007-33375 | OR910817 | | | | This study | |
| | | 5. Maritime College, Segond Channel, Vanuatu | -15.5226 | 167.1659 | MNHN-IM-2007-33383 | OR910818 | | | | This study | |
| 84 | *Terestrombus terebellatus* (G. B. Sowerby II, 1842) | **1. Kavieng Lagoon, Papua New Guinea** | **-2.5867** | **150.485** | **MNHN-IM-2013-54138** | **OR911252** | **OR934426** | **OR934477** | **OR934377** | **This study** | |
| 85 | *Terestrombus fragilis* (Röding, 1798) | 1. Guam | – | – | NV | DQ525216 | | | | Latiolais et al. (2006) | |
| | | 2. Balicasag Is., Philippines | 9.5167 | 123.6833 | MNHN-IM-2007-34707 | OR934185 | | | | This study | |
| | | **3. Balicasag Is., Philippines** | **9.5167** | **123.6833** | **MNHN-IM-2007-34708** | **OR910982** | **OR934186** | | | **This study** | |
| 86 | *Tridentarius dentatus* (Linnaeus, 1758) | 1. S Kranket Is., Papua New Guinea | -5.205 | 145.8133 | MNHN-IM-2013-12550 | OR910988 | OR934184 | OR934322 | OR934063 | This study | |
| | | **2. Kendec Is., New Caledonia** | **-20.6683** | **164.2567** | **MNHN-IM-2019-1738** | **MW244820** | **MW244820** | **MW244820** | **MW244820** | **Irwin et al. (2021)** | |
| | | 3. Maria, Austral Islands, French Polynesia | -21.8029 | -154.7196 | MNHN-IM-2013-48340 | OR910989 | | OR934323 | OR934064 | This study | |
| | | 4. Maria, Austral Islands, French Polynesia | -21.8029 | -154.7196 | MNHN-IM-2013-48342 | OR910990 | | | | This study | |

| | | | | | | | | | | |
|---|---|---|---|---|---|---|---|---|---|---|
| | | 5. Scorff Passage, NW Tutuba Is., Vanuatu | -15.5183 | 167.2633 | MNHN-IM-2007-33335 | OR910991 | | | | This study |
| | | 6. Orote Peninsula, Guam | – | – | NV | DQ525215 | | | | Latiolais et al. (2006) |
| 87 | *Canarium manintveldi* Dekkers & S. J. Maxwell, 2020 | **1. S Palikulo Bay, Vanuatu** | **-15.4931** | **167.2487** | **MNHN-IM-2007-33362** | **OR910736** | **OR934103** | **OR934235** | | **This study** |
| 88 | *Canarium darwinense* S. J. Maxwell & Dekkers, 2021 | **1. Darwin Harbour, Northern Territory, Australia** | **-12.4917** | **130.8967** | **NTM P058594** | **OR911222** | **OR934395** | **OR934446** | **OR934347** | **This study** |
| 89 | *Canarium incisum* (W. Wood, 1828) | 1. Yagaji Is. Harbour, Nago, Okinawa, Japan | 26.61 | 127.9933 | NHMUK 2019034 | OR910731 | OR934104 | OR934236 | OR934068 | This study |
| | | 2. Yagaji Is. Harbour, Nago, Okinawa, Japan | **26.61** | **127.9933** | **NHMUK 2019033** | **OR911221** | **OR934394** | **OR934445** | **OR934346** | **This study** |
| | | 3. Wenchang, Hainan Province, China | – | – | OUC SUWC01 | JF693423 | | | | Sun et al. (2012) |
| | | 4. Wenchang, Hainan Province, China | – | – | OUC SUWC02 | JF693424 | | | | Sun et al. (2012) |
| | | 5. Wenchang, Hainan Province, China | – | – | OUC SUWC03 | JF693425 | | | | Sun et al. (2012) |
| | | 6. Linchangjiao, Hainan Province, China | – | – | OUC G182 | MN389049 | | | | Ran et al. (2020) |
| | | 7. Linchangjiao, Hainan Province, China | – | – | OUC G183 | MN389050 | | | | Ran et al. (2020) |

| | | | | | | | | | | |
|---|---|---|---|---|---|---|---|---|---|---|
| | | 8. Linchangjiao, Hainan Province, China | – | – | OUC G184 | MN389051 | | | | Ran et al. (2020) |
| | | 9. Kavieng Lagoon, Papua New Guinea | -2.6867 | 150.69 | MNHN-IM-2013-47991 | OR910730 | OR934105 | OR934237 | OR933978 | This study |
| 90 | *Canarium olydium* (Duclos, 1844) | **1. N Pemba Bay headland, Mozambique** | – | – | **NHMUK 20060448** | **OR911219** | | **OR934443** | | **This study** |
| 91 | *Canarium erythrinum* (Dillwyn, 1817) | 1. In front of Narendry Bay, Madagascar | -14.5317 | 47.4423 | MNHN-IM-2007-36759 | OR910723 | | | | This study |
| | | 2. In front of Narendry Bay, Madagascar | -14.5317 | 47.4423 | MNHN-IM-2007-36733 | OR910722 | | | | This study |
| | | **3. In front of Narendry Bay, Madagascar** | **-14.5317** | **47.4423** | **MNHN-IM-2007-36698** | **OR911212** | **OR934386** | **OR934436** | **OR934338** | **This study** |
| | | 4. In front of Flacourt lighthouse, S Madagascar | -25.0217 | 47.0083 | MNHN-IM-2009-15839 | OR910719 | OR934082 | | | This study |
| 92 | *Canarium labiatum* (Röding, 1798) | 1. Pasarwajo Bay, SE Sulawesi, Indonesia | 24.35 | 124.1 | NHMUK 20050686 | OR910732 | OR934088 | OR934221 | OR933979 | This study |
| | | **2. Kavieng Lagoon, Papua New Guinea** | **-2.6867** | **150.69** | **MNHN-IM-2013-47992** | **OR911215** | **OR934389** | **OR934439** | **OR934341** | **This study** |
| | | 3. Onna-son, Okinawa Is., Japan | 26.4833 | 127.8333 | NHMUK 2019027 | OR910733 | OR934089 | OR934222 | OR933980 | This study |
| | | 4. Doljo Point, Panglao Is., Philippines | 9.5917 | 123.7217 | MNHN-IM-2007-33348 | OR910734 | | | | This study |
| | | 5. Bingag/Tabalong, Panglao Is., Philippines | 9.63 | 123.8067 | MNHN-IM-2007-33349 | OR910735 | | | | This study |

| | | | | | | | | | | |
|---|---|---|---|---|---|---|---|---|---|---|
| 93 | *Canarium radians* (Duclos, 1844) | 1. Coolidge wreck, Segond Channel, Vanuatu | -15.5233 | 167.235 | MNHN-IM-2007-33331 | OR910727 | | | | This study |
| | | 2. Tangadiou Is., New Caledonia | -20.5533 | 164.2167 | MNHN-IM-2019-1731 | OR910717 | OR934083 | OR934218 | OR933976 | This study |
| | | **3. Kranket Is., Papua New Guinea** | **-5.2** | **145.8083** | **MNHN-IM-2013-11485** | **OR910716** | **OR934081** | **OR934217** | **OR933975** | **This study** |
| | | 4. W Kranket Is., Papua New Guinea | -5.195 | 145.8133 | MNHN-IM-2013-11478 | OR910721 | | | | This study |
| | | 5. Wonat Is., Papua New Guinea | -5.1333 | 145.8217 | MNHN-IM-2013-10029 | OR910720 | OR934085 | | | This study |
| | | 6. Coolidge wreck, Segond Channel, Vanuatu | -15.5233 | 167.235 | MNHN-IM-2007-33366 | OR910728 | | | | This study |
| | | 7. Coolidge wreck, Segond Channel, Vanuatu | -15.5233 | 167.235 | MNHN-IM-2007-33334 | OR910729 | | | | This study |
| | | 8. Coolidge wreck, Segond Channel, Vanuatu | -15.5233 | 167.235 | MNHN-IM-2007-33371 | OR910724 | | | | This study |
| | | 9. Coolidge wreck, Segond Channel, Vanuatu | -15.5233 | 167.235 | MNHN-IM-2007-33368 | OR910725 | | | | This study |
| | | 10. Coolidge wreck, Segond Channel, Vanuatu | -15.5233 | 167.235 | MNHN-IM-2007-33367 | OR910726 | | | | This study |
| | | 11. W Ratua Is., S Aoré Is., Vanuatu | -15.61 | 167.1683 | MNHN-IM-2007-33384 | OR910718 | OR934084 | OR934219 | OR933977 | This study |
| 94 | | 1. Rangiroa, Tuamotu Is., French Polynesia | – | – | NV | DQ525217 | | | | Latiolais et al. (2006) |

| | | | | | | | | | | | |
|---|---|---|---|---|---|---|---|---|---|---|---|
| | *Maculastrombus maculatus* (G. B. Sowerby II, 1842) | **2. Rimatara, French Polynesia** | **-22.6332** | **-152.8041** | **MNHN-IM-2013-48346** | **OR910857** | **OR934093** | **OR934226** | **OR934035** | **This study** | |
| | | 3. Rimatara, French Polynesia | -22.6332 | -152.8041 | MNHN-IM-2013-48348 | OR910858 | OR934094 | OR934227 | OR934036 | This study | |
| | | 4. Rimatara, French Polynesia | -22.6332 | -152.8041 | MNHN-IM-2013-48345 | OR910859 | | | | This study | |
| 95 | *Maculastrombus microurceus* (Kira, 1959) comb. nov. | 1. Kavieng Lagoon, Papua New Guinea | -2.5867 | 150.77 | MNHN-IM-2013-46788 | OR910737 | OR934095 | OR934228 | OR933981 | This study | Formerly *Canarium microurceum* Kira, 1959 |
| | | 2. Kranket Is., Papua New Guinea | -5.1883 | 145.8233 | MNHN-IM-2013-10138 | OR910738 | | | | This study | |
| | | **3. Kavieng Lagoon, Papua New Guinea** | **-2.6283** | **150.67** | **MNHN-IM-2013-55168** | **OR911217** | **OR934391** | **OR934441** | **OR934343** | **This study** | |
| | | 4. Orote Peninsula, Guam | – | – | NV | DQ525230 | | | | Latiolais et al. (2006) | |
| 96 | *Maculastrombus mutabilis* A comb. nov. | 1. Amani-U-Shima, Japan | – | – | NV | DQ525218 | | | | Latiolais et al. (2006) | Two putative cryptic species of (formerly) *Canarium mutabile* (Swainson, 1821). |
| | | **2. N Riwo, Papua New Guinea** | **-5.145** | **145.8033** | **MNHN-IM-2013-18307** | **OR910741** | **OR934099** | **OR934231** | **OR933984** | **This study** | |
| | | 3. Wenchang, Hainan Province, China | – | – | OUC G87 | MN389052 | | | | Ran et al. (2020) | |
| | | 4. Palikulo Peninsula, Vanuatu | -15.48 | 167.255 | MNHN-IM-2007-33380 | OR910762 | | | | This study | |
| | | 5. Rurutu, French Polynesia | -22.4449 | -151.3446 | MNHN-IM-2013-48347 | OR910742 | OR934100 | OR934232 | OR933985 | This study | |
| | | 6. Raivavae, French Polynesia | -23.8826 | -147.6461 | MNHN-IM-2013-48339 | OR910760 | | | | This study | |
| | | 7. Bingag/Tabalong, Panglao Is., Philippines | 9.63 | 123.8067 | MNHN-IM-2007-33344 | OR910740 | OR934098 | OR934230 | OR933983 | This study | |
| | | 8. Cape Bise, Motobu, Okinawa Is., Japan | 26.71 | 127.8767 | NHMUK 2019030 | | OR934096 | | | This study | |

| | | | | | | | | | | |
|---|---|---|---|---|---|---|---|---|---|---|
| 97 | *Maculastrombus mutabilis* B comb. nov. | 1. Evatra Point, S Madagascar | -24.9683 | 47.1017 | MNHN-IM-2009-15855 | OR910748 | | | | This study |
| | | **2. Monseigneur Beach, S Madagascar** | **-24.9417** | **47.1183** | **MNHN-IM-2009-15843** | **OR911218** | **OR934392** | **OR934442** | **OR934344** | **This study** |
| | | 3. Ambatobe, Near Soamanitse, S Madagascar | -25.4567 | 44.9567 | MNHN-IM-2009-15857 | OR910744 | | | | This study |
| | | 4. Ambatobe, Near Soamanitse, S Madagascar | -25.4567 | 44.9567 | MNHN-IM-2009-15845 | OR910743 | | | | This study |
| | | 5. Ambatomainty, S Madagascar | -25.4383 | 44.9417 | MNHN-IM-2009-15848 | OR910746 | | | | This study |
| | | 6. Evatra Point, S Madagascar | -24.9683 | 47.1017 | MNHN-IM-2009-15851 | OR910749 | | | | This study |
| | | 7. Ambatomainty, S Madagascar | -25.4383 | 44.9417 | MNHN-IM-2009-15849 | OR910745 | | | | This study |
| | | 8. Evatra Point, S Madagascar | -24.9683 | 47.1017 | MNHN-IM-2009-15837 | OR910750 | | | | This study |
| | | 9. Monseigneur Beach, S Madagascar | -25.035 | 46.9983 | MNHN-IM-2009-15841 | OR910757 | | | | This study |
| | | 10. Ambatobe, Near Soamanitse, S Madagascar | -25.4567 | 44.9567 | MNHN-IM-2009-15838 | OR910747 | | | | This study |
| | | 11. Lokaro Is., S Madagascar | -24.942 | 47.118 | MNHN-IM-2009-15850 | OR910751 | | | | This study |
| | | 12. Inhaca Is., Maputo Bay, Mozambique | -26.0733 | 32.9517 | MNHN-IM-2013-64496 | OR910759 | | | | This study |
| | | 13. N Pemba Bay headland, Mozambique | -25.035 | 46.9983 | NHMUK 20060446 | OR910739 | OR934097 | OR934229 | OR933982 | This study |

| | | | | | | | | | | | |
|---|---|---|---|---|---|---|---|---|---|---|---|
| | | 14. Monseigneur Beach, S Madagascar | -25.035 | 46.9983 | MNHN-IM-2009-15856 | OR910752 | | | | This study | |
| | | 15. Monseigneur Beach, S Madagascar | -25.035 | 46.9983 | MNHN-IM-2009-15853 | OR910753 | | | | This study | |
| | | 16. Monseigneur Beach, S Madagascar | -25.035 | 46.9983 | MNHN-IM-2009-15846 | OR910754 | | | | This study | |
| | | 17. Monseigneur Beach, S Madagascar | -25.035 | 46.9983 | MNHN-IM-2009-15844 | OR910755 | | | | This study | |
| | | 18. Monseigneur Beach, S Madagascar | -25.035 | 46.9983 | MNHN-IM-2009-15842 | OR910756 | | | | This study | |
| | | 19. Monseigneur Beach, S Madagascar | -25.035 | 46.9983 | MNHN-IM-2009-15847 | OR910758 | | | | This study | |
| | | 20. Ranavalona Cape, S Madagascar | -25.075 | 46.9633 | MNHN-IM-2009-15854 | OR910761 | | | | This study | |
| 98 | *Dolomena campbellii* (J. E. Gray, 1833) comb. nov. | **2. Casuarina Beach, Darwin, Australia** | – | – | **NTM P058595** | | **OR934106** | | | **This study** | Formerly *Doxander campbellii* (J. E. Gray, 1833) |
| 99 | *Dolomena japonica* (Reeve, 1851) comb. nov. | **1. Tateyama, Chiba, Japan** | – | – | **NHMUK 20230926** | | **OR934109** | | **OR933998** | **This study** | Formerly *Doxander japonicus* (Reeve, 1851) |
| 100 | *Dolomena wienekei* Wiersma & D. Monsecour, 2012 | **1. Yomba Is., Papua New Guinea** | **-5.245** | **145.7883** | **MNHN-IM-2013-17824** | | **OR934400** | **OR934451** | **OR934351** | **This study** | |
| Outgroup: | | | | | | | | | | | |

| | | | | | | | | | |
|---|---|---|---|---|---|---|---|---|---|
| *Echinolittorina interrupta* (R. A. Philippi, 1847) | Bull Is., South Carolina, USA | – | – | ANSP A19977-01* | AJ623011 | | | | Rosenberg and Sei (2012) |
| | Bocas del Toro, Colon Is., Panama | – | – | NHMUK NINT.BDT.1* | | JQ990646 | JQ990538 | AJ488672 | Williams and Reid (2004) |
| *Littorina littorea* (Linnaeus, 1758) | Koster Arch., Sweden | – | – | NHMUK LLIT.URS.1* | AJ622946 | | | AJ488672 | Williams and Reid (2004) |
| | Dominion Beach, Nova Scotia, Canada | 46.217 | -60.038 | FMNH 336506* | | KC583385 | KC583385 | | Golding et al. (2014) |
