## Supplementary Material 3 for "Molecular phylogenetics of the superfamily Stromboidea (Caenogastropoda): new insights from increased taxon sampling"

**Supplementary Material 3.** Literature for fossil calibrations and geographic ranges, and characteristics of shell morphology in stromboid families (table), used to identify fossils to family level to aid finding the earliest representatives of each group. A complete reference list is given below the table, with those used for extant geographic ranges of stromboids (not included in the main text) added.

| | **Aporrhaidae** | **Xenophoridae** | **Struthiolariidae** | **Rostellariidae** | **Seraphsidae** | **Strombidae** |
|---|---|---|---|---|---|---|
| **Overall shape** | Fusiform | Low, depressed to conical, trochiform | Fusiform, inflated, ovate | Slender, fusiform | Long, torpedo-shaped, fusiform | Fusiform |
| **Protoconch shape** | Dome-shaped, four convex whorls | Rounded or median angular; 3 to 4 whorls, depressed to conical | Rounded whorls | 3 to over 6 rounded whorls, conical | Flattened, relatively large | 3 to 4.5, rounded whorls; conical |
| **Teleoconch shape** | Tall, conical spire with apical angle of ca. 35-40° | Broad base (flattened or concave). Narrow to wide, simple to spinose peripheral flange | Angular spire; large, stout, rounded lower whorls | High spire and ovoid body whorl | Body whorl covers majority or all of the higher whorls | Diverse shapes |
| **Protoconch sculpture** | Smooth | Smooth or with spiral ribs | Smooth | Smooth | Smooth | Growth lines, and may be smooth or ornamented |
| **Teleoconch sculpture** | Heavily sculptured spire and body whorl, with axial and/or spiral ridges | Typically lacks sculpture (except growth lines and a spiral cord). Diverse objects (e.g. shells, coral rubble, pebbles) agglutinated to surface | Nodulose spire; smooth lower whorls (except axial growth lines). Columella broad, callused | Variable; smooth to ornamented by axial ribs and variable varices | Smooth and lacking sculpture (few exceptions) | Spire and body whorl often heavily sculptured with ridges and knobs, but can be almost smooth. Thick columellar callus, often with teeth or lirae |
| **Aperture** | Elongate, with long siphonal canal | Large, strongly oblique, with thin margins. Siphonal canal absent. | Large, oval, thickened margin. Short siphonal canal | Elongated or elliptical. Anterior siphonal canal is typically long (shorter in | Slightly flaring or thin. Shell forms a wide siphonal groove | Narrow, with short siphonal canal |

| | | | | | | |
|---|---|---|---|---|---|---|
| | | | | Rimellinae), continuing in a narrow posterior groove | | |
| **Outer lip** | Thickened and expanded outer lip. Three or more projecting digits –variety of spines, hooks, knobs, and fan-like shapes with internal grooves | No terminate growth (outer lip not thickened) | Thickened outer lip | Elongate and pointed with 'stromboid notch' (Rimellinae), or thickened, convex outer lip. Often with ridges or wrinkles, and a variable number of finger-like projections. | Basal margin of the body whorl does not extend to the level of the columella base, or continues in a curved callus groove to the apex | Thickened and expanded outer lip. Shallow to well-defined 'stromboid notch' at anterior end. Other diverse sculpture present on outer lip |
| **References** | Popenoe, 1983; Bandel 1993; Bandel et al., 1997; Kiel and Bandel, 1999; Simone, 2005; Gründel et al., 2009 | Morton, 1958; Ponder 1983; Bandel, 1993; Wells, 1998; Kreipl and Alf, 1999; Simone, 2005; Bandel, 2007 | Wells, 1998; Bandel, 2007 | Kronenberg and Berkhout, 1984; Wells, 1998; Bandel, 2007 | Wells, 1998; Jung, 1974; Kronenberg and Berkhout, 1984; Simone, 2005; Bandel, 2007 | Walls, 1980; Popenoe, 1983; Savazzi, 1991; Simone, 2005; Bandel, 2007 |
