## Supplementary Material 4 for "Molecular phylogenetics of the superfamily Stromboidea (Caenogastropoda): new insights from increased taxon sampling"

**Supplementary Material 4.** Summary of COI species delimitation results. Putative species are listed in the order shown in Fig. 2 and Suppl. Mat. 5 and numbered (no.) for ease of cross-reference in these figures and Table 2, with number of specimens (n). Whether putative species were delimited in each method (Y, yes; S, split; L, lump) is listed with total ESUs delimited by each method in parentheses in column headings. Relevant support values (parentheses) are: ASV, values closer to one signify closer agreement with the ML delimitation; GYMC, values closer to one indicate maximal support for delimited species. Support for species nodes in trees is also given (BEAST, IQ-TREE) (Fig. 2 and Suppl. Mat. 5–6). Comments added if species boundaries differ from current species concepts (text highlighted in bold), or when delimitation results differ among methods. No support values were given for species represented by a single sequence (NA).

| **Putative species** | | | | **Putative species delimited** | | | | | **Phylogenetic support for putative species** | | |
|---|---|---|---|---|---|---|---|---|---|---|---|
| **No.** | **Family** | **Species/clade** | **n** | **GMYC (102)** | **bPTP (104)** | **mPTP (86)** | **ASAP (97)** | **ABGD (92)** | **BEAST** | **IQ-TREE** | **Comments** |
| **1** | Xenophoridae | *Onustus indicus* | 1 | Y (NA) | Y (1) | L (0.94) | Y | Y | NA | NA | mPTP: putative species 12–15 delimited as single ESU. |
| **2** | Xenophoridae | *Aspidophoreas chinensis* | 1 | Y (NA) | Y (1) | | Y | Y | NA | NA | |
| **3** | Xenophoridae | *Stellaria solaris* | 1 | Y (NA) | Y (1) | | Y | Y | NA | NA | |
| **4** | Xenophoridae | *Onustus exutus* | 1 | Y (NA) | Y (1) | | Y | Y | NA | NA | |
| **5** | Xenophoridae | *Xenophora solarioides* | 2 | Y (1) | Y (1) | Y (0.93) | Y | Y | 1 | 100 | |
| **6** | Xenophoridae | *Xenophora pallidula* | 1 | Y (NA) | Y (1) | L (0.93) | Y | Y | NA | NA | mPTP: putative species 17–21 delimited as single ESU. |
| **7** | Xenophoridae | *Xenophora japonica* | 2 | Y (1) | Y (1) | | Y | Y | 1 | 100 | |
| **8** | Xenophoridae | *Onustus longleyi* | 2 | Y (1) | Y (1) | | Y | Y | 1 | 100 | |
| **9** | Xenophoridae | *Xenophora conchyliophora* | 1 | Y (NA) | Y (1) | | Y | Y | NA | NA | |
| **10** | Xenophoridae | *Onustus caribaeus* | 2 | Y (1) | Y (1) | | Y | Y | 1 | 100 | |

| | | | | | | | | | | | |
|---|---|---|---|---|---|---|---|---|---|---|---|
| **11** | Struthiolariidae | *Struthiolaria papulosa* | 2 | Y (1) | Y (1) | Y (0.84) | Y | Y | 0.99 | 83 | |
| **12** | Struthiolariidae | *Pelicaria vermis* | 2 | Y (1) | Y (1) | Y (0.84) | Y | Y | 1 | 100 | |
| **13** | Aporrhaidae | *Aporrhais serresiana* | 3 | Y (1) | Y (1) | L (1) | Y | Y | 1 | 100 | mPTP: putative species 5–6 delimited as single ESU. bPTP: oversplit species 6 into six ESUs |
| **14** | Aporrhaidae | *Aporrhais pespelecani* | 10 | Y (1) | S (all: 1) | | Y | Y | 0.99 | 81 | |
| **15** | **Seraphsidae** | ***Terebellum terebellum* A** | **1** | **Y (NA)** | **Y (1)** | **Y (1)** | **Y** | **Y** | **NA** | **NA** | |
| **16** | **Seraphsidae** | ***Terebellum terebellum* B** | **5** | **Y (1)** | **Y (1)** | **Y (1)** | **Y** | **Y** | **1** | **100** | |
| **17** | Seraphsidae | *Terebellum delicatum* | 5 | Y (1) | Y (1) | Y (1) | Y | Y | 1 | 100 | |
| **18** | **Seraphsidae** | ***Terebellum terebellum* C** | **1** | **Y (NA)** | **Y (1)** | **Y (1)** | **Y** | **Y** | **NA** | **NA** | |
| **19** | **Seraphsidae** | ***Terebellum terebellum* D** | **6** | **S (0.81/1)** | **Y (1)** | **Y (1)** | **Y** | **Y** | **1** | **100** | **GMYC: split (two ESUs)** |
| **20** | Rostellariidae | *Varicospira crispata* | 2 | Y (1) | Y (1) | Y (1) | Y | Y | 1 | 100 | |
| **21** | Rostellariidae | *Varicospira cancellata* | 3 | Y (1) | Y (1) | Y (1) | Y | Y | 1 | 100 | |
| **22** | Rostellariidae | *Tibia fusus* | 1 | Y (NA) | Y (1) | L (1) | Y | Y | NA | NA | mPTP: putative species 22–23 delimited as single ESU |
| **23** | Rostellariidae | *Tibia insulaechorab* | 2 | Y (1) | Y (1) | | Y | Y | 1 | 100 | |
| **24** | Rostellariidae | *Tenuitibia martinii* | 5 | Y (1) | Y (1) | Y (1) | Y | Y | 1 | 100 | |
| **25** | Rostellariidae | *Rostellariella lorenzi* | 1 | Y (NA) | Y (0.94) | Y (1) | Y | L | NA | NA | ABGD: putative species 25–26 delimited as single ESU |
| **26** | Rostellariidae | *Rostellariella delicatula* | 28 | Y (1) | Y (0.94) | Y (1) | Y | | 0.98 | 67 | |
| **27** | **Rostellariidae** | ***Rimellopsis powisii*** | **2** | **Y (1)** | **Y (1)** | **Y (1)** | **Y** | **Y** | **1** | **100** | |
| **28** | **Rostellariidae** | ***Rimellopsis laurenti*** | **49** | **Y (1)** | **Y (1)** | **Y (1)** | **Y** | **Y** | **1** | **100** | |
| **29** | Strombidae | *Barneystrombus kleckhamae* | 3 | Y (1) | Y (1) | Y (1) | Y | Y | 1 | 100 | |

| | | | | | | | | | | | |
|---|---|---|---|---|---|---|---|---|---|---|---|
| **30** | Strombidae | *Euprotomus aurisdianae* | 2 | Y (1) | Y (1) | L (0.86) | Y | Y | 1 | 100 | |
| **31** | Strombidae | *Euprotomus aratrum* | 1 | Y (NA) | Y (1) | Y (0.86) | Y | Y | NA | NA | |
| **32** | **Strombidae** | ***Euprotomus bulla + Euprotomus aurora*** | **8** | **Y (1)** | **Y (1)** | **Y (1)** | **Y** | **Y** | **1** | **99** | All methods delimit *E. bulla + E. aurora* as single ESU |
| **33** | Strombidae | *Thetystrombus latus* | 3 | Y (1) | Y (1) | Y (1) | Y | Y | 1 | 100 | |
| **34** | Strombidae | *Persististrombus granulatus* | 2 | Y (1) | S (0.24) | Y (1) | Y | Y | 1 | 100 | bPTP: split (two ESUs) |
| **35** | Strombidae | *Strombus gracilior* | 2 | Y (1) | Y (1) | Y (1) | Y | Y | 1 | 100 | |
| **36** | **Strombidae** | ***Strombus pugilis + Strombus alatus*** | **4** | **S (0.65/0.65)** | **Y (1)** | **Y (1)** | **Y** | **Y** | **1** | **100** | GMYC: split (two ESUs). All other methods delimit *S. pugilis + S. alatus* as single ESU |
| **37** | Strombidae | *Titanostrombus galeatus* | 2 | Y (1) | Y (1) | Y (1) | Y | Y | 1 | 100 | |
| **38** | Strombidae | *Lobatus raninus* | 8 | Y (1) | Y (1) | Y (1) | Y | Y | 1 | 100 | |
| **39** | Strombidae | *Aliger gallus* | 6 | Y (1) | Y (1) | Y (1) | Y | Y | 1 | 100 | |
| **40** | Strombidae | *Aliger costatus* | 11 | Y (1) | Y (1) | Y (1) | Y | Y | 1 | 100 | |
| **41** | Strombidae | *Aliger gigas* | 18 | Y (1) | Y (1) | Y (1) | Y | Y | 1 | 100 | |
| **42** | Strombidae | *Gibberulus gibberulus* | 5 | S (0.95/NA) | Y (1) | Y (1) | Y | Y | 1 | 97 | GMYC: split (two ESUs) |
| **43** | Strombidae | *Gibberulus gibbosus* | 6 | Y (1) | Y (1) | Y (1) | Y | Y | 1 | 100 | |
| **44** | Strombidae | *Conomurex fasciatus* | 2 | Y (1) | Y (1) | Y (1) | Y | Y | 1 | 100 | |
| **45** | Strombidae | *Conomurex persicus* | 1 | Y (NA) | Y (0.95) | Y (0.86) | Y | L | NA | NA | ABGD: putative species 45–46 delimited as single ESU |
| **46** | Strombidae | *Conomurex decorus* | 7 | Y (1) | Y (0.95) | Y (0.86) | Y | | 1 | 100 | |

| | | | | | | | | | | | |
|---|---|---|---|---|---|---|---|---|---|---|---|
| **47** | Strombidae | *Conomurex luhuanus* | 16 | S (0.84/0.76) | Y (1) | Y (0.96) | Y | Y | 1 | 97 | GMYC: split (two ESUs) |
| **48** | Strombidae | *Lentigo pipus* | 2 | Y (1) | Y (1) | Y (1) | Y | Y | 1 | 100 | |
| **49** | Strombidae | *Lentigo thersites* | 1 | Y (NA) | Y (1) | Y (0.98) | Y | Y | NA | NA | |
| **50** | Strombidae | *Lentigo lentiginosus* | 4 | Y (1) | Y (1) | Y (0.98) | Y | Y | 1 | 100 | |
| **51** | Strombidae | *Latissistrombus sinuatus* | 2 | Y (1) | Y (1) | Y (0.93) | Y | Y | 1 | 100 | |
| **52** | Strombidae | *Latissistrombus latissimus* | 2 | Y (1) | Y (1) | Y (0.93) | Y | Y | 1 | 100 | |
| **53** | Strombidae | *Latissistrombus taurus* | 3 | Y (1) | Y (1) | Y (0.97) | Y | Y | 1 | 100 | |
| **54** | Strombidae | *Ophioglossolambis digitata* | 1 | Y (NA) | Y (1) | L (0.99) | Y | Y | NA | NA | mPTP: putative species 54–55 delimited as single ESU |
| **55** | Strombidae | *Tricornis tricornis* | 2 | Y (1) | Y (1) | | Y | Y | 1 | 100 | |
| **56** | **Strombidae** | ***Harpago chiragra*** | **4** | **Y (1)** | **Y (1)** | **Y (0.99)** | **Y** | **Y** | **1** | **100** | |
| **57** | **Strombidae** | ***Harpago rugosa*** | **1** | **Y (NA)** | **Y (1)** | **Y (0.99)** | **Y** | **Y** | **NA** | **NA** | |
| **58** | Strombidae | *Lambis scorpius* | 5 | Y (1) | Y (1) | Y (1) | Y | Y | 1 | 100 | |
| **59** | Strombidae | *Lambis millepeda* | 4 | Y (1) | Y (1) | Y (1) | Y | Y | 1 | 100 | |
| **60** | **Strombidae** | ***Lambis truncata*** | **1** | **Y (NA)** | **Y (1)** | **Y (0.77)** | **Y** | **L** | **NA** | **NA** | ABGD: putative species 60–61 delimited as single ESU |
| **61** | **Strombidae** | ***Lambis sowerbyi*** | **5** | **Y (1)** | **Y (1)** | **Y (0.77)** | **Y** | | **1** | **100** | |
| **62** | Strombidae | *Lambis crocata* | 3 | Y (1) | Y (1) | Y (1) | Y | Y | 1 | 100 | |
| **63** | Strombidae | *Lambis robusta* | 2 | Y (1) | Y (1) | Y (1) | Y | Y | 1 | 100 | |
| **64** | **Strombidae** | **_Lambis lambis_ A** | **2** | **Y (1)** | **Y (1)** | **Y (0.99)** | **Y** | **Y** | **1** | **100** | |
| **65** | **Strombidae** | **_Lambis lambis_ B** | **7** | **Y (1)** | **Y (1)** | **Y (0.99)** | **Y** | **Y** | **1** | **100** | |
| **66** | **Strombidae** | **_Lambis lambis_ C** | **4** | **Y (1)** | **Y (1)** | **Y (0.99)** | **Y** | **Y** | **1** | **100** | |
| **67** | **Strombidae** | **_"Canarium" wilsonorum_ A** | **2** | **Y (1)** | **Y (1)** | **Y (0.87)** | **Y** | **Y** | **1** | **100** | |
| **68** | **Strombidae** | **_"Canarium" wilsonorum_ B** | **1** | **Y (NA)** | **Y (1)** | **Y (0.87)** | **Y** | **Y** | **NA** | **NA** | |

| | | | | | | | | | | | |
|---|---|---|---|---|---|---|---|---|---|---|---|
| **69** | Strombidae | *Fusistrombus fusiformis* | 3 | Y (1) | Y (1) | Y (1) | Y | Y | 1 | 100 | |
| **70** | Strombidae | *Hawaiistrombus scalariforme* | 3 | Y (1) | Y (1) | Y (1) | Y | Y | 1 | 100 | |
| **71** | Strombidae | *Dolomena epidromis* | 2 | Y (1) | Y (1) | Y (1) | Y | Y | 1 | 100 | |
| **72** | **Strombidae** | ***Dolomena vittata + Dolomena operosa*** | **9** | **S (1/1)** | **Y (1)** | **Y (1)** | **Y** | **Y** | **1** | **100** | **GMYC split (two ESUs, (1) *D. vittata + D. operosa;* (2) *D. operosa*). All other methods delimit *D. vittata + D. operosa* as single ESU** |
| **73** | **Strombidae** | ***Dolomena labiosa + Dolomena abbotti*** | **8** | **Y (1)** | **Y (1)** | **Y (1)** | **Y** | **Y** | **1** | **100** | **All methods delimit *D. labiosa + D. abbotti* as single ESU** |
| **74** | Strombidae | *Dolomena robusta* | 6 | Y (1) | Y (1) | Y (1) | Y | Y | 1 | 100 | |
| **75** | Strombidae | *Dolomena variabilis* | 4 | Y (1) | Y (1) | Y (1) | Y | Y | 1 | 100 | |
| **76** | Strombidae | *Dolomena pulchella* | 3 | Y (1) | Y (1) | Y (1) | Y | Y | 1 | 100 | |
| **77** | Strombidae | *Ministrombus minimus* | 4 | Y (1) | Y (1) | Y (1) | Y | Y | 1 | 100 | |
| **78** | Strombidae | *Dolomena swainsoni* | 2 | Y (1) | L (1) | L (1) | Y | L | 1 | 100 | bPTP, mPTP and ABGD: putative species 74–75 delimited as single ESU |
| **79** | Strombidae | *Dolomena dilatata* | 2 | Y (1) | | | Y | | 0.99 | 100 | |
| **80** | Strombidae | *Dolomena turturella* A | 3 | Y (0.97) | S (0.74) | Y (0.9) | Y | Y | 1 | 100 | bPTP: split (two ESUs) |
| **81** | Strombidae | *Dolomena turturella* B | 2 | Y (1) | Y (1) | Y (1) | Y | Y | 1 | 100 | |
| **82** | Strombidae | *Dolomena taeniata* | 3 | Y (1) | S (1) | S (0.34) | Y | Y | 1 | 100 | bPTP and mPTP: split (two ESUs); IQ-TREE did not resolve *D. taeniata* |
| **83** | Strombidae | *Dolomena vanikorensis* | 5 | Y (1) | Y (1) | Y (1) | Y | Y | 1 | 100 | |

| | | | | | | | | | | | |
|---|---|---|---|---|---|---|---|---|---|---|---|
| **84** | Strombidae | *Terestrombus terebellatus* | 1 | Y (NA) | Y (1) | Y (0.9) | Y | Y | NA | NA | |
| **85** | Strombidae | *Terestrombus fragilis* | 2 | Y (1) | Y (1) | Y (0.9) | Y | Y | 1 | 100 | |
| **86** | Strombidae | *Canarium incisum* | 9 | Y (1) | Y (1) | Y (0.99) | Y | Y | 1 | 100 | |
| **87** | Strombidae | *Tridentarius dentatus* | 6 | Y (1) | Y (1) | Y (1) | Y | Y | 1 | 100 | |
| **88** | Strombidae | *Canarium olydium* | 1 | Y (NA) | Y (1) | Y (0.98) | Y | Y | NA | NA | |
| **89** | Strombidae | *Canarium erythrinum* | 4 | Y (1) | Y (1) | Y (0.98) | Y | Y | 1 | 100 | |
| **90** | Strombidae | *Canarium labiatum* | 5 | Y (1) | Y (1) | Y (1) | Y | Y | 1 | 100 | |
| **91** | Strombidae | *Canarium manintveldi* | 1 | Y (NA) | Y (1) | L (0.99) | Y | Y | NA | NA | mPTP: putative species 90–91 delimited as single ESU |
| **92** | Strombidae | *Canarium darwinense* | 1 | Y (NA) | Y (1) | | Y | Y | NA | NA | |
| **93** | Strombidae | *Canarium radians* | 11 | Y (1) | Y (1) | Y (1) | Y | Y | 1 | 100 | |
| **94** | Strombidae | *Maculastrombus maculatus* | 4 | Y (1) | Y (1) | Y (1) | Y | Y | 1 | 100 | |
| **95** | Strombidae | *Maculastrombus microurceus* | 4 | Y (1) | Y (1) | Y (1) | Y | Y | 1 | 100 | |
| **96** | **Strombidae** | ***Maculastrombus mutabilis* A** | **7** | **Y (1)** | **Y (1)** | **Y (1)** | **Y** | **L** | **1** | **100** | ABGD: putative species 88–89 delimited as single ESU |
| **97** | **Strombidae** | ***Maculastrombus mutabilis* B** | **20** | **Y (0.63)** | **Y (1)** | **Y (1)** | **Y** | | **1** | **94** | |
