## Supplementary Material 5 for "Molecular phylogenetics of the superfamily Stromboidea (Caenogastropoda): new insights from increased taxon sampling"

**Supplementary Material 5.** Bayesian analysis of COI sequences via BEAST (enlarged version of Figure 2 over three pages; separated for ease of reading values of posterior probability). Specimen number and country of collection are given at tips (Suppl. Mat. 2). If primary species hypotheses lumped taxa, nominal species are listed in grey to the right. ESUs delimited by each method (section 3.2) are depicted via blocks (black, accepted ESUs; red, unaccepted splits; orange, unaccepted lumps), with putative species on the right (green). Putative species names are listed on the far right, with corresponding number in parentheses; species with changed generic assignments are in red font. Support values are posterior probabilities (PP); intraspecific support values were removed for legibility except where relevant (i.e., if there was conflict between delimited ESUs or prior species assignations). Inserts displayed on each page shows position within the tree.

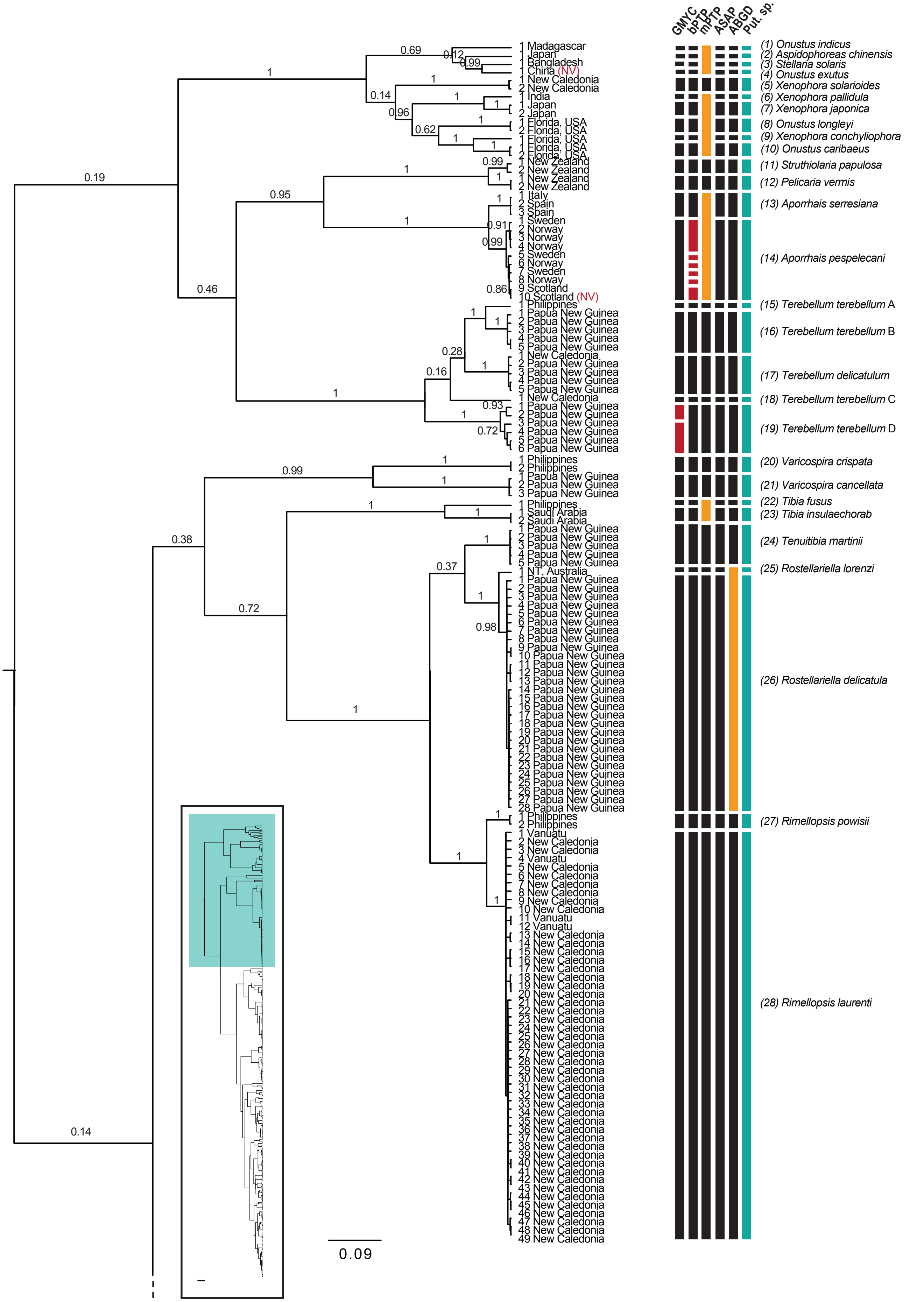

GMYC
bPTP
mPTP
ASAP
ABGD
Put. sp.
(1) Onustus indicus
(2) Aspidophoreas chinensis
(3) Stellaria solaris
(4) Onustus exutus
(5) Xenophora solarioides
(6) Xenophora pallidula
(7) Xenophora japonica
(8) Onustus longleyi
(9) Xenophora conchyliophora
(10) Onustus caribaeus
(11) Struthiolaria papulosa
(12) Pelicaria vermis
(13) Aporrhais serresiana
(14) Aporrhais pespelecani
(15) Terebellum terebellum A
(16) Terebellum terebellum B
(17) Terebellum delicatulum
(18) Terebellum terebellum C
(19) Terebellum terebellum D
(20) Varicospira crispata
(21) Varicospira cancellata
(22) Tibia fusus
(23) Tibia insulaechorab
(24) Tenuitibia martinii
(25) Rostellariella lorenzi
(26) Rostellariella delicatula
(27) Rimellopsis powisii
(28) Rimellopsis laurenti
1 Madagascar
1 Japan
1 Bangladesh
1 China (NV)
1 New Caledonia
2 New Caledonia
1 India
1 Japan
2 Japan
1 Florida, USA
2 Florida, USA
1 Florida, USA
1 Florida, USA
2 Florida, USA
1 New Zealand
2 New Zealand
1 New Zealand
2 New Zealand
1 Italy
2 Spain
3 Spain
1 Sweden
2 Norway
3 Norway
4 Norway
5 Sweden
6 Norway
7 Sweden
8 Norway
9 Scotland
10 Scotland (NV)
1 Philippines
1 Papua New Guinea
2 Papua New Guinea
3 Papua New Guinea
4 Papua New Guinea
5 Papua New Guinea
1 New Caledonia
2 Papua New Guinea
3 Papua New Guinea
4 Papua New Guinea
5 Papua New Guinea
1 New Caledonia
1 Papua New Guinea
2 Papua New Guinea
3 Papua New Guinea
4 Papua New Guinea
5 Papua New Guinea
6 Papua New Guinea
1 Philippines
2 Philippines
1 Papua New Guinea
2 Papua New Guinea
3 Papua New Guinea
1 Philippines
1 Saudi Arabia
2 Saudi Arabia
1 Papua New Guinea
2 Papua New Guinea
3 Papua New Guinea
4 Papua New Guinea
5 Papua New Guinea
1 NT, Australia
1 Papua New Guinea
2 Papua New Guinea
3 Papua New Guinea
4 Papua New Guinea
5 Papua New Guinea
6 Papua New Guinea
7 Papua New Guinea
8 Papua New Guinea
9 Papua New Guinea
10 Papua New Guinea
11 Papua New Guinea
12 Papua New Guinea
13 Papua New Guinea
14 Papua New Guinea
15 Papua New Guinea
16 Papua New Guinea
17 Papua New Guinea
18 Papua New Guinea
19 Papua New Guinea
20 Papua New Guinea
21 Papua New Guinea
22 Papua New Guinea
23 Papua New Guinea
24 Papua New Guinea
25 Papua New Guinea
26 Papua New Guinea
27 Papua New Guinea
28 Papua New Guinea
1 Philippines
2 Philippines
1 Vanuatu
2 New Caledonia
3 New Caledonia
4 Vanuatu
5 New Caledonia
6 New Caledonia
7 New Caledonia
8 New Caledonia
9 New Caledonia
10 New Caledonia
11 Vanuatu
12 Vanuatu
13 New Caledonia
14 New Caledonia
15 New Caledonia
16 New Caledonia
17 New Caledonia
18 New Caledonia
19 New Caledonia
20 New Caledonia
21 New Caledonia
22 New Caledonia
23 New Caledonia
24 New Caledonia
25 New Caledonia
26 New Caledonia
27 New Caledonia
28 New Caledonia
29 New Caledonia
30 New Caledonia
31 New Caledonia
32 New Caledonia
33 New Caledonia
34 New Caledonia
35 New Caledonia
36 New Caledonia
37 New Caledonia
38 New Caledonia
39 New Caledonia
40 New Caledonia
41 New Caledonia
42 New Caledonia
43 New Caledonia
44 New Caledonia
45 New Caledonia
46 New Caledonia
47 New Caledonia
48 New Caledonia
49 New Caledonia
0.09

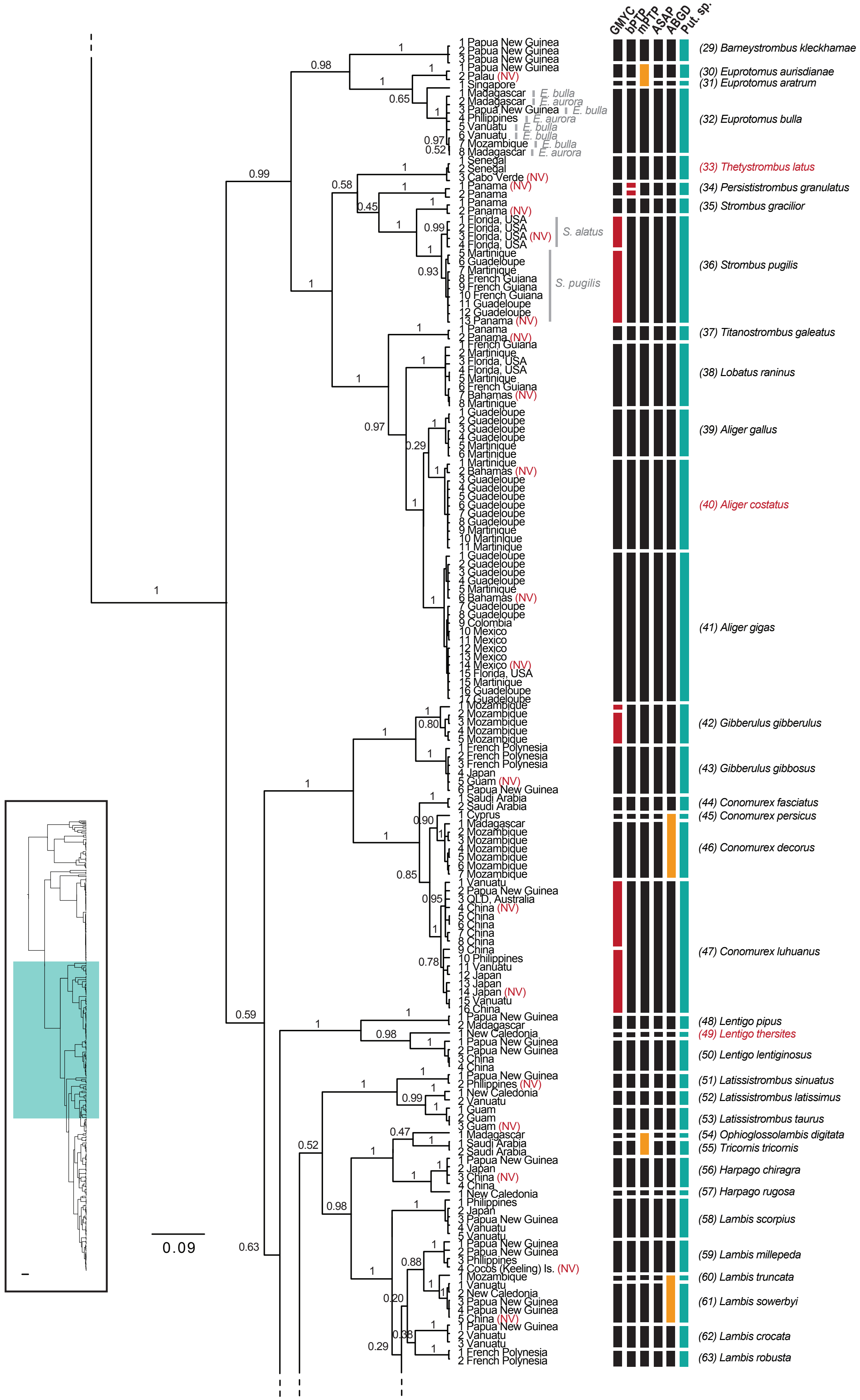
GMYC
bPTP
mPTP
ASAP
ABGD
Put. sp.
1 Papua New Guinea
2 Papua New Guinea
3 Papua New Guinea
(29) Barneystrombus kleckhamae
1 Papua New Guinea
2 Palau (NV)
(30) Euprotomus aurisdianae
1 Singapore
(31) Euprotomus aratrum
1 Madagascar E. bulla
2 Madagascar E. aurora
3 Papua New Guinea E. bulla
4 Philippines E. aurora
5 Vanuatu E. bulla
6 Vanuatu E. bulla
7 Mozambique E. bulla
8 Madagascar E. aurora
(32) Euprotomus bulla
1 Senegal
2 Senegal
3 Cabo Verde (NV)
(33) Thetystrombus latus
1 Panama (NV)
2 Panama
(34) Persististrombus granulatus
1 Panama
2 Panama (NV)
(35) Strombus gracilior
1 Florida, USA
2 Florida, USA
3 Florida, USA (NV)
4 Florida, USA
S. alatus
5 Martinique
6 Guadeloupe
7 Martinique
8 French Guiana
9 French Guiana
10 French Guiana
11 Guadeloupe
12 Guadeloupe
13 Panama (NV)
S. pugilis
(36) Strombus pugilis
1 Panama
2 Panama (NV)
(37) Titanostrombus galeatus
1 French Guiana
2 Martinique
3 Florida, USA
4 Florida, USA
5 Martinique
6 French Guiana
7 Bahamas (NV)
8 Martinique
(38) Lobatus raninus
1 Guadeloupe
2 Guadeloupe
3 Guadeloupe
4 Guadeloupe
5 Martinique
6 Martinique
(39) Aliger gallus
1 Martinique
2 Bahamas (NV)
3 Guadeloupe
4 Guadeloupe
5 Guadeloupe
6 Guadeloupe
7 Guadeloupe
8 Guadeloupe
9 Martinique
10 Martinique
11 Martinique
(40) Aliger costatus
1 Guadeloupe
2 Guadeloupe
3 Guadeloupe
4 Guadeloupe
5 Martinique
6 Bahamas (NV)
7 Guadeloupe
8 Guadeloupe
9 Colombia
10 Mexico
11 Mexico
12 Mexico
13 Mexico
14 Mexico (NV)
15 Florida, USA
15 Martinique
16 Guadeloupe
17 Guadeloupe
(41) Aliger gigas
1 Mozambique
2 Mozambique
3 Mozambique
4 Mozambique
5 Mozambique
(42) Gibberulus gibberulus
1 French Polynesia
2 French Polynesia
3 French Polynesia
4 Japan
5 Guam (NV)
6 Papua New Guinea
(43) Gibberulus gibbosus
1 Saudi Arabia
2 Saudi Arabia
(44) Conomurex fasciatus
1 Cyprus
(45) Conomurex persicus
1 Madagascar
2 Mozambique
3 Mozambique
4 Mozambique
5 Mozambique
6 Mozambique
7 Mozambique
(46) Conomurex decorus
1 Vanuatu
2 Papua New Guinea
3 QLD, Australia
4 China (NV)
5 China
6 China
7 China
8 China
9 China
10 Philippines
11 Vanuatu
12 Japan
13 Japan
14 Japan (NV)
15 Vanuatu
16 China
(47) Conomurex luhuanus
1 Papua New Guinea
2 Madagascar
(48) Lentigo pipus
1 New Caledonia
(49) Lentigo thersites
1 Papua New Guinea
2 Papua New Guinea
3 China
4 China
(50) Lentigo lentiginosus
1 Papua New Guinea
2 Philippines (NV)
(51) Latissistrombus sinuatus
1 New Caledonia
2 Vanuatu
(52) Latissistrombus latissimus
1 Guam
2 Guam
3 Guam (NV)
(53) Latissistrombus taurus
1 Madagascar
(54) Ophioglossolambis digitata
1 Saudi Arabia
2 Saudi Arabia
(55) Tricornis tricornis
1 Papua New Guinea
2 Japan
3 China (NV)
4 China
(56) Harpago chiragra
1 New Caledonia
(57) Harpago rugosa
1 Philippines
2 Japan
3 Papua New Guinea
4 Vanuatu
5 Vanuatu
(58) Lambis scorpius
1 Papua New Guinea
2 Papua New Guinea
3 Philippines
4 Cocos (Keeling) Is. (NV)
(59) Lambis millepeda
1 Mozambique
(60) Lambis truncata
1 Vanuatu
2 New Caledonia
3 Papua New Guinea
4 Papua New Guinea
5 China (NV)
(61) Lambis sowerbyi
1 Papua New Guinea
2 Vanuatu
3 Vanuatu
(62) Lambis crocata
1 French Polynesia
2 French Polynesia
(63) Lambis robusta
0.09

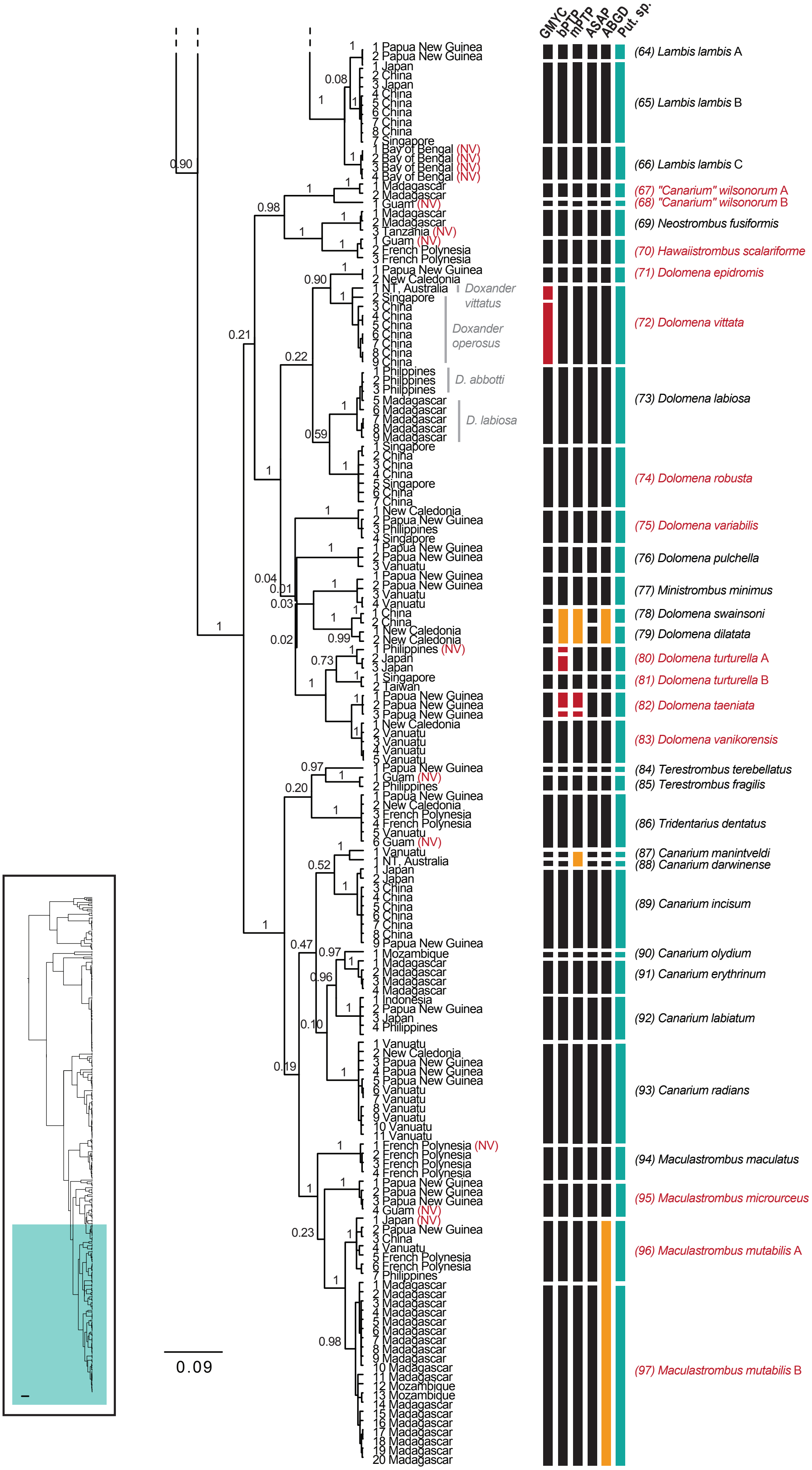
GMYC
bPTP
mPTP
ASAP
ABGD
Put. sp.
1 Papua New Guinea
2 Papua New Guinea
1 Japan
2 China
3 Japan
4 China
5 China
6 China
7 China
8 China
7 Singapore
1 Bay of Bengal (NV)
2 Bay of Bengal (NV)
3 Bay of Bengal (NV)
4 Bay of Bengal (NV)
1 Madagascar
2 Madagascar
1 Guam (NV)
1 Madagascar
2 Madagascar
3 Tanzania (NV)
1 Guam (NV)
2 French Polynesia
3 French Polynesia
1 Papua New Guinea
2 New Caledonia
1 NT, Australia
2 Singapore
3 China
4 China
5 China
6 China
7 China
8 China
9 China
1 Philippines
2 Philippines
3 Philippines
5 Madagascar
6 Madagascar
7 Madagascar
8 Madagascar
9 Madagascar
1 Singapore
2 China
3 China
4 China
5 Singapore
6 China
7 China
1 New Caledonia
2 Papua New Guinea
3 Philippines
4 Singapore
1 Papua New Guinea
2 Papua New Guinea
3 Vanuatu
1 Papua New Guinea
2 Papua New Guinea
3 Vanuatu
4 Vanuatu
1 China
2 China
1 New Caledonia
2 New Caledonia
1 Philippines (NV)
2 Japan
3 Japan
1 Singapore
2 Taiwan
1 Papua New Guinea
2 Papua New Guinea
3 Papua New Guinea
1 New Caledonia
2 Vanuatu
3 Vanuatu
4 Vanuatu
5 Vanuatu
1 Papua New Guinea
1 Guam (NV)
2 Philippines
1 Papua New Guinea
2 New Caledonia
3 French Polynesia
4 French Polynesia
5 Vanuatu
6 Guam (NV)
1 Vanuatu
1 NT, Australia
1 Japan
2 Japan
3 China
4 China
5 China
6 China
7 China
8 China
9 Papua New Guinea
1 Mozambique
1 Madagascar
2 Madagascar
3 Madagascar
4 Madagascar
1 Indonesia
2 Papua New Guinea
3 Japan
4 Philippines
1 Vanuatu
2 New Caledonia
3 Papua New Guinea
4 Papua New Guinea
5 Papua New Guinea
6 Vanuatu
7 Vanuatu
8 Vanuatu
9 Vanuatu
10 Vanuatu
11 Vanuatu
1 French Polynesia (NV)
2 French Polynesia
3 French Polynesia
4 French Polynesia
1 Papua New Guinea
2 Papua New Guinea
3 Papua New Guinea
4 Guam (NV)
1 Japan (NV)
2 Papua New Guinea
3 China
4 Vanuatu
5 French Polynesia
6 French Polynesia
7 Philippines
1 Madagascar
2 Madagascar
3 Madagascar
4 Madagascar
5 Madagascar
6 Madagascar
7 Madagascar
8 Madagascar
9 Madagascar
10 Madagascar
11 Madagascar
12 Mozambique
13 Mozambique
14 Madagascar
15 Madagascar
16 Madagascar
17 Madagascar
18 Madagascar
19 Madagascar
20 Madagascar
Doxander vittatus
Doxander operosus
D. abbotti
D. labiosa
(64) Lambis lambis A
(65) Lambis lambis B
(66) Lambis lambis C
(67) "Canarium" wilsonorum A
(68) "Canarium" wilsonorum B
(69) Neostrombus fusiformis
(70) Hawaiistrombus scalariforme
(71) Dolomena epidromis
(72) Dolomena vittata
(73) Dolomena labiosa
(74) Dolomena robusta
(75) Dolomena variabilis
(76) Dolomena pulchella
(77) Ministrombus minimus
(78) Dolomena swainsoni
(79) Dolomena dilatata
(80) Dolomena turturella A
(81) Dolomena turturella B
(82) Dolomena taeniata
(83) Dolomena vanikorensis
(84) Terestrombus terebellatus
(85) Terestrombus fragilis
(86) Tridentarius dentatus
(87) Canarium manintveldi
(88) Canarium darwinense
(89) Canarium incisum
(90) Canarium olydium
(91) Canarium erythrinum
(92) Canarium labiatum
(93) Canarium radians
(94) Maculastrombus maculatus
(95) Maculastrombus microurceus
(96) Maculastrombus mutabilis A
(97) Maculastrombus mutabilis B
0.09
