## Supplementary Material 6 for "Molecular phylogenetics of the superfamily Stromboidea (Caenogastropoda): new insights from increased taxon sampling"

**Supplementary Material 6.** Bayesian analysis of COI sequences via IQTREE. Specimen number and country of collection are given at tips (Suppl. Mat. 2). If primary species hypotheses lumped species together, the nominal species are listed in grey to the right of the collection locality. Putative species names are listed on the far right (green blocks), with corresponding number in parentheses. Support values are posterior probabilities (PP); branches with PP < 50% are collapsed, and intraspecific support values were removed for legibility except where relevant (i.e., if there was conflict between delimited ESUs or prior species assignations). Inserts displayed on each page shows position within the tree.

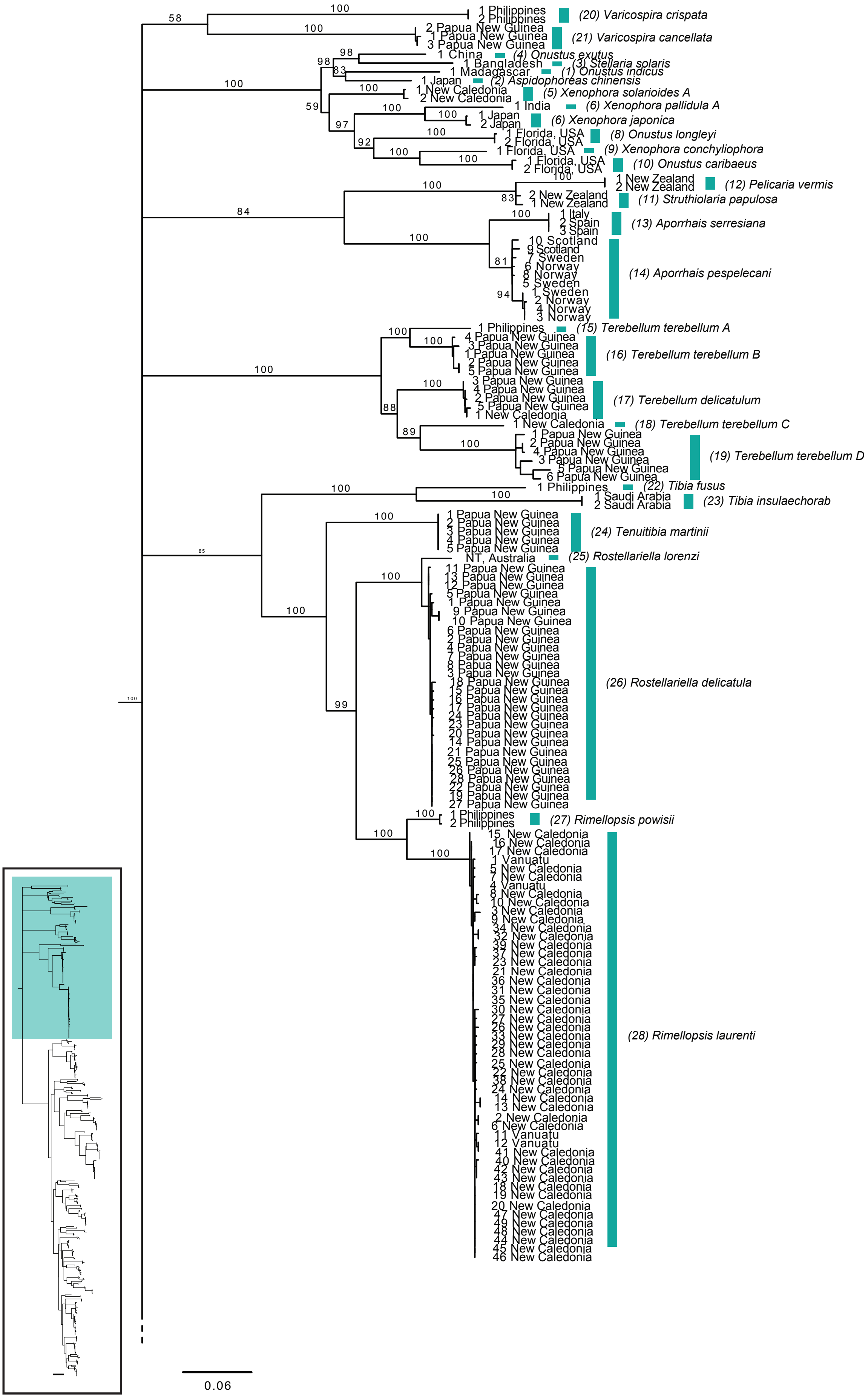

1 Philippines
2 Philippines
(20) Varicospira crispata
2 Papua New Guinea
1 Papua New Guinea
3 Papua New Guinea
(21) Varicospira cancellata
1 China
(4) Onustus exutus
1 Bangladesh
(3) Stellaria solaris
1 Madagascar
(1) Onustus indicus
1 Japan
(2) Aspidophoreas chinensis
1 New Caledonia
2 New Caledonia
(5) Xenophora solarioides A
1 India
(6) Xenophora pallidula A
1 Japan
2 Japan
(6) Xenophora japonica
1 Florida, USA
2 Florida, USA
(8) Onustus longleyi
1 Florida, USA
(9) Xenophora conchyliophora
1 Florida, USA
2 Florida, USA
(10) Onustus caribaeus
1 New Zealand
2 New Zealand
(12) Pelicaria vermis
2 New Zealand
1 New Zealand
(11) Struthiolaria papulosa
1 Italy
2 Spain
3 Spain
(13) Aporrhais serresiana
10 Scotland
9 Scotland
7 Sweden
6 Norway
8 Norway
5 Sweden
1 Sweden
2 Norway
4 Norway
3 Norway
(14) Aporrhais pespelecani
1 Philippines
(15) Terebellum terebellum A
4 Papua New Guinea
3 Papua New Guinea
1 Papua New Guinea
2 Papua New Guinea
5 Papua New Guinea
(16) Terebellum terebellum B
3 Papua New Guinea
4 Papua New Guinea
2 Papua New Guinea
5 Papua New Guinea
1 New Caledonia
(17) Terebellum delicatulum
1 New Caledonia
(18) Terebellum terebellum C
1 Papua New Guinea
2 Papua New Guinea
4 Papua New Guinea
3 Papua New Guinea
5 Papua New Guinea
6 Papua New Guinea
(19) Terebellum terebellum D
1 Philippines
(22) Tibia fusus
1 Saudi Arabia
2 Saudi Arabia
(23) Tibia insulaechorab
1 Papua New Guinea
2 Papua New Guinea
3 Papua New Guinea
4 Papua New Guinea
5 Papua New Guinea
(24) Tenuitibia martinii
NT, Australia
(25) Rostellariella lorenzi
11 Papua New Guinea
13 Papua New Guinea
12 Papua New Guinea
5 Papua New Guinea
1 Papua New Guinea
9 Papua New Guinea
10 Papua New Guinea
6 Papua New Guinea
2 Papua New Guinea
4 Papua New Guinea
7 Papua New Guinea
8 Papua New Guinea
3 Papua New Guinea
18 Papua New Guinea
15 Papua New Guinea
16 Papua New Guinea
17 Papua New Guinea
24 Papua New Guinea
23 Papua New Guinea
20 Papua New Guinea
14 Papua New Guinea
21 Papua New Guinea
25 Papua New Guinea
26 Papua New Guinea
28 Papua New Guinea
22 Papua New Guinea
19 Papua New Guinea
27 Papua New Guinea
(26) Rostellariella delicatula
1 Philippines
2 Philippines
(27) Rimellopsis powisii
15 New Caledonia
16 New Caledonia
17 New Caledonia
1 Vanuatu
5 New Caledonia
7 New Caledonia
4 Vanuatu
8 New Caledonia
10 New Caledonia
3 New Caledonia
9 New Caledonia
34 New Caledonia
32 New Caledonia
39 New Caledonia
37 New Caledonia
23 New Caledonia
21 New Caledonia
36 New Caledonia
31 New Caledonia
35 New Caledonia
30 New Caledonia
27 New Caledonia
26 New Caledonia
33 New Caledonia
29 New Caledonia
28 New Caledonia
25 New Caledonia
22 New Caledonia
38 New Caledonia
24 New Caledonia
14 New Caledonia
13 New Caledonia
2 New Caledonia
6 New Caledonia
11 Vanuatu
12 Vanuatu
41 New Caledonia
40 New Caledonia
42 New Caledonia
43 New Caledonia
18 New Caledonia
19 New Caledonia
20 New Caledonia
47 New Caledonia
49 New Caledonia
48 New Caledonia
44 New Caledonia
45 New Caledonia
46 New Caledonia
(28) Rimellopsis laurenti
0.06

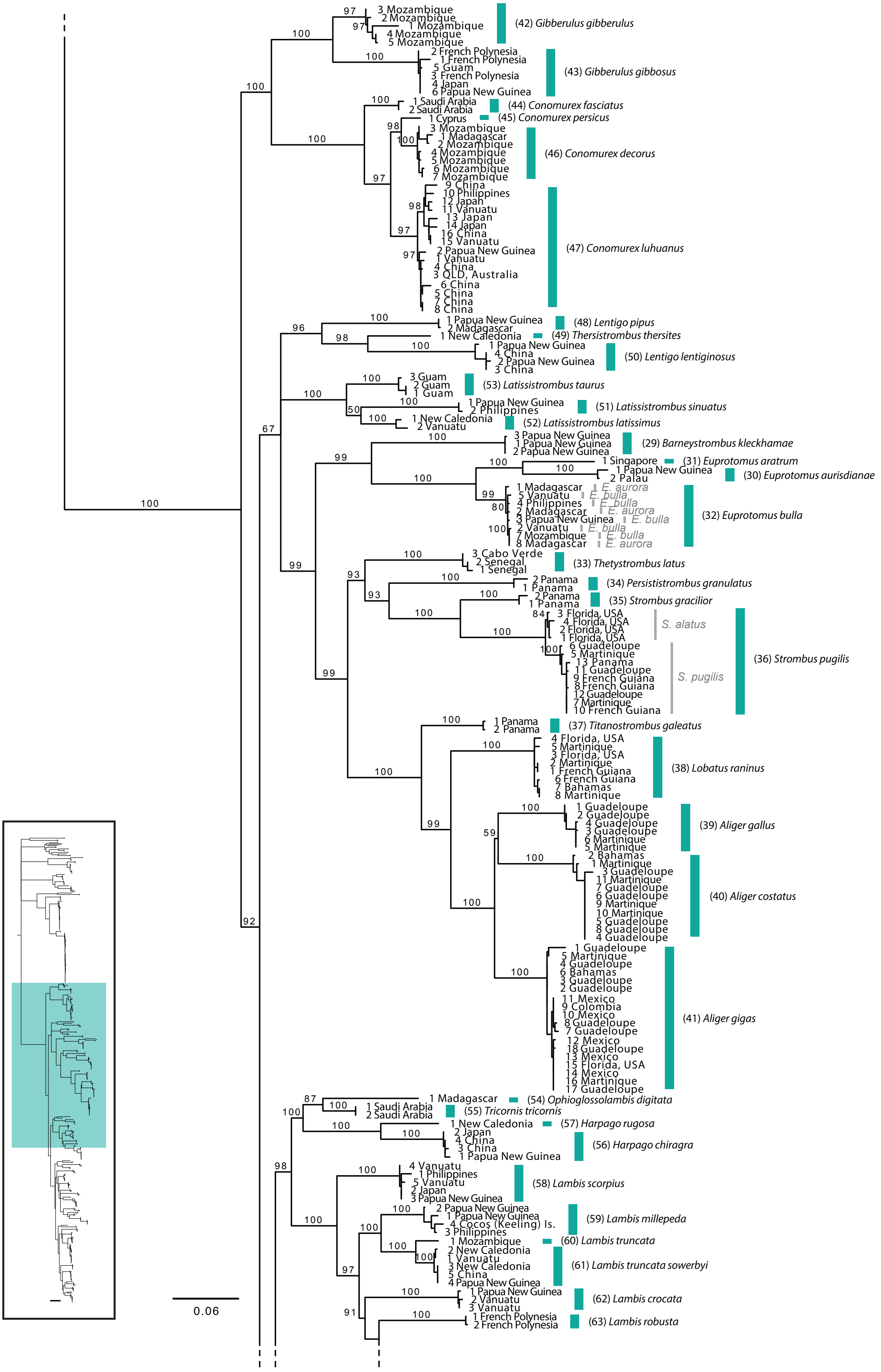
3 Mozambique
2 Mozambique
1 Mozambique
4 Mozambique
5 Mozambique
(42) Gibberulus gibberulus
2 French Polynesia
1 French Polynesia
5 Guam
3 French Polynesia
4 Japan
6 Papua New Guinea
(43) Gibberulus gibbosus
1 Saudi Arabia
2 Saudi Arabia
(44) Conomurex fasciatus
1 Cyprus
(45) Conomurex persicus
3 Mozambique
1 Madagascar
2 Mozambique
4 Mozambique
5 Mozambique
6 Mozambique
7 Mozambique
(46) Conomurex decorus
9 China
10 Philippines
12 Japan
11 Vanuatu
13 Japan
14 Japan
16 China
15 Vanuatu
2 Papua New Guinea
1 Vanuatu
4 China
3 QLD, Australia
6 China
5 China
7 China
8 China
(47) Conomurex luhuanus
1 Papua New Guinea
2 Madagascar
(48) Lentigo pipus
1 New Caledonia
(49) Thersistrombus thersites
1 Papua New Guinea
4 China
2 Papua New Guinea
3 China
(50) Lentigo lentiginosus
3 Guam
2 Guam
1 Guam
(53) Latissistrombus taurus
1 Papua New Guinea
2 Philippines
(51) Latissistrombus sinuatus
1 New Caledonia
2 Vanuatu
(52) Latissistrombus latissimus
3 Papua New Guinea
1 Papua New Guinea
2 Papua New Guinea
(29) Barneystrombus kleckhamae
1 Singapore
(31) Euprotomus aratrum
1 Papua New Guinea
2 Palau
(30) Euprotomus aurisdianae
1 Madagascar
5 Vanuatu
4 Philippines
2 Madagascar
3 Papua New Guinea
2 Vanuatu
7 Mozambique
8 Madagascar
E. aurora
E. bulla
E. bulla
E. aurora
E. bulla
E. bulla
E. bulla
E. aurora
(32) Euprotomus bulla
3 Cabo Verde
2 Senegal
1 Senegal
(33) Thetystrombus latus
2 Panama
1 Panama
(34) Persististrombus granulatus
2 Panama
1 Panama
(35) Strombus gracilior
3 Florida, USA
4 Florida, USA
2 Florida, USA
1 Florida, USA
S. alatus
6 Guadeloupe
5 Martinique
13 Panama
11 Guadeloupe
9 French Guiana
8 French Guiana
12 Guadeloupe
7 Martinique
10 French Guiana
S. pugilis
(36) Strombus pugilis
1 Panama
2 Panama
(37) Titanostrombus galeatus
4 Florida, USA
5 Martinique
3 Florida, USA
2 Martinique
1 French Guiana
6 French Guiana
7 Bahamas
8 Martinique
(38) Lobatus raninus
1 Guadeloupe
2 Guadeloupe
4 Guadeloupe
3 Guadeloupe
6 Martinique
5 Martinique
(39) Aliger gallus
2 Bahamas
1 Martinique
3 Guadeloupe
11 Martinique
7 Guadeloupe
6 Guadeloupe
9 Martinique
10 Martinique
5 Guadeloupe
8 Guadeloupe
4 Guadeloupe
(40) Aliger costatus
1 Guadeloupe
5 Martinique
4 Guadeloupe
6 Bahamas
3 Guadeloupe
2 Guadeloupe
11 Mexico
9 Colombia
10 Mexico
8 Guadeloupe
7 Guadeloupe
12 Mexico
18 Guadeloupe
13 Mexico
15 Florida, USA
14 Mexico
16 Martinique
17 Guadeloupe
(41) Aliger gigas
1 Madagascar
(54) Ophioglossolambis digitata
1 Saudi Arabia
2 Saudi Arabia
(55) Tricornis tricornis
1 New Caledonia
(57) Harpago rugosa
2 Japan
4 China
3 China
1 Papua New Guinea
(56) Harpago chiragra
4 Vanuatu
1 Philippines
5 Vanuatu
2 Japan
3 Papua New Guinea
(58) Lambis scorpius
2 Papua New Guinea
1 Papua New Guinea
4 Cocos (Keeling) Is.
3 Philippines
(59) Lambis millepeda
1 Mozambique
(60) Lambis truncata
2 New Caledonia
1 Vanuatu
3 New Caledonia
5 China
4 Papua New Guinea
(61) Lambis truncata sowerbyi
1 Papua New Guinea
2 Vanuatu
3 Vanuatu
(62) Lambis crocata
1 French Polynesia
2 French Polynesia
(63) Lambis robusta
0.06

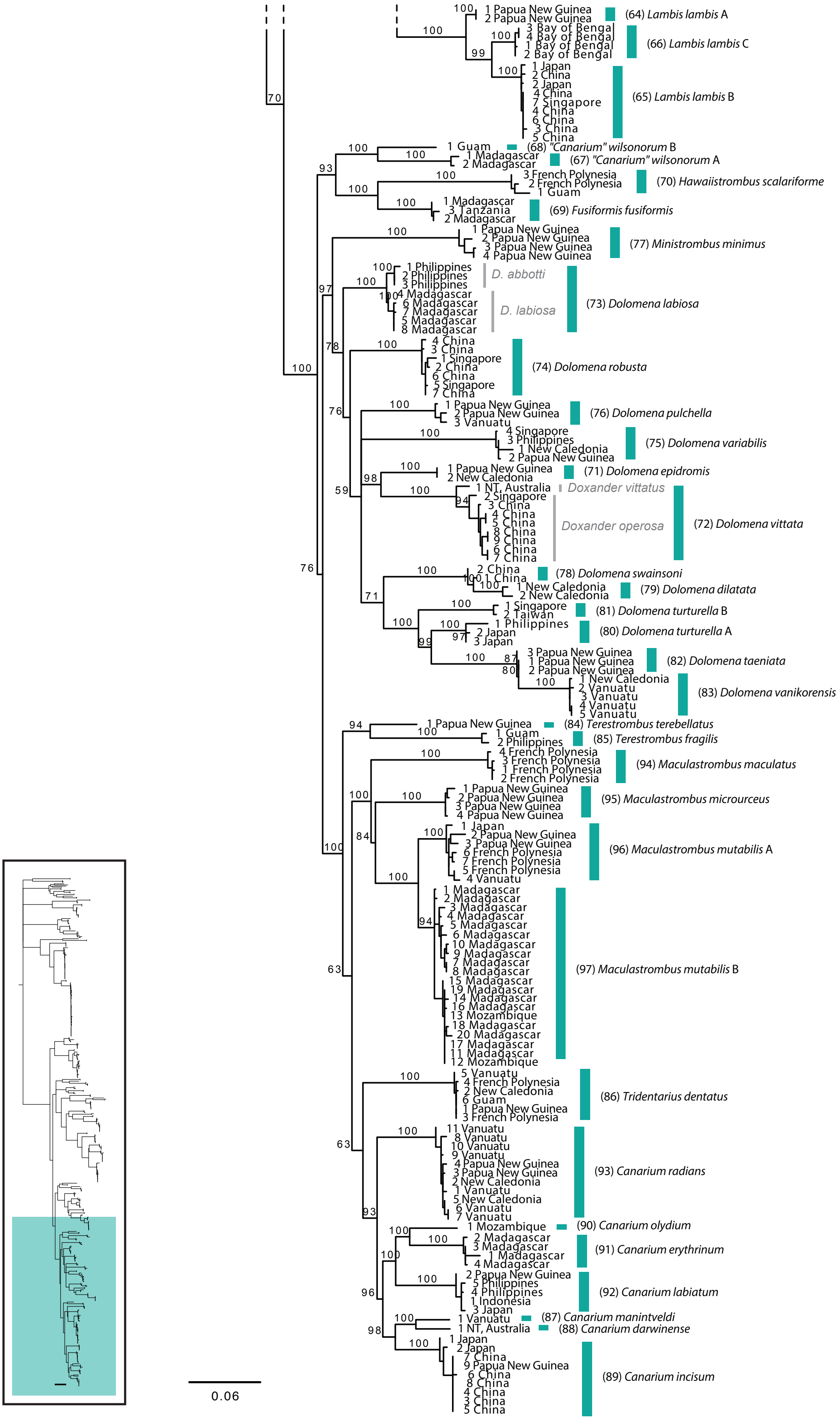

1 Papua New Guinea
2 Papua New Guinea
(64) Lambis lambis A
3 Bay of Bengal
4 Bay of Bengal
1 Bay of Bengal
2 Bay of Bengal
(66) Lambis lambis C
1 Japan
2 China
2 Japan
4 China
7 Singapore
4 China
6 China
3 China
5 China
(65) Lambis lambis B
1 Guam
(68) "Canarium" wilsonorum B
1 Madagascar
2 Madagascar
(67) "Canarium" wilsonorum A
3 French Polynesia
2 French Polynesia
1 Guam
(70) Hawaiistrombus scalariforme
1 Madagascar
3 Tanzania
2 Madagascar
(69) Fusiformis fusiformis
1 Papua New Guinea
2 Papua New Guinea
3 Papua New Guinea
4 Papua New Guinea
(77) Ministrombus minimus
1 Philippines
2 Philippines
3 Philippines
D. abbotti
4 Madagascar
6 Madagascar
7 Madagascar
5 Madagascar
8 Madagascar
D. labiosa
(73) Dolomena labiosa
4 China
3 China
1 Singapore
2 China
6 China
5 Singapore
7 China
(74) Dolomena robusta
1 Papua New Guinea
2 Papua New Guinea
3 Vanuatu
(76) Dolomena pulchella
4 Singapore
3 Philippines
1 New Caledonia
2 Papua New Guinea
(75) Dolomena variabilis
1 Papua New Guinea
2 New Caledonia
(71) Dolomena epidromis
1 NT, Australia
Doxander vittatus
2 Singapore
3 China
4 China
5 China
8 China
9 China
6 China
7 China
Doxander operosa
(72) Dolomena vittata
2 China
1 China
(78) Dolomena swainsoni
1 New Caledonia
2 New Caledonia
(79) Dolomena dilatata
1 Singapore
2 Taiwan
(81) Dolomena turturella B
1 Philippines
2 Japan
3 Japan
(80) Dolomena turturella A
3 Papua New Guinea
1 Papua New Guinea
2 Papua New Guinea
(82) Dolomena taeniata
1 New Caledonia
2 Vanuatu
3 Vanuatu
4 Vanuatu
5 Vanuatu
(83) Dolomena vanikorensis
1 Papua New Guinea
(84) Terestrombus terebellatus
1 Guam
2 Philippines
(85) Terestrombus fragilis
4 French Polynesia
3 French Polynesia
1 French Polynesia
2 French Polynesia
(94) Maculastrombus maculatus
1 Papua New Guinea
2 Papua New Guinea
3 Papua New Guinea
4 Papua New Guinea
(95) Maculastrombus microurceus
1 Japan
2 Papua New Guinea
3 Papua New Guinea
6 French Polynesia
7 French Polynesia
5 French Polynesia
4 Vanuatu
(96) Maculastrombus mutabilis A
1 Madagascar
2 Madagascar
3 Madagascar
4 Madagascar
5 Madagascar
6 Madagascar
10 Madagascar
9 Madagascar
7 Madagascar
8 Madagascar
15 Madagascar
19 Madagascar
14 Madagascar
16 Madagascar
13 Mozambique
18 Madagascar
20 Madagascar
17 Madagascar
11 Madagascar
12 Mozambique
(97) Maculastrombus mutabilis B
5 Vanuatu
4 French Polynesia
2 New Caledonia
6 Guam
1 Papua New Guinea
3 French Polynesia
(86) Tridentarius dentatus
11 Vanuatu
8 Vanuatu
10 Vanuatu
9 Vanuatu
4 Papua New Guinea
3 Papua New Guinea
2 New Caledonia
1 Vanuatu
5 New Caledonia
6 Vanuatu
7 Vanuatu
(93) Canarium radians
1 Mozambique
(90) Canarium olydium
2 Madagascar
3 Madagascar
1 Madagascar
4 Madagascar
(91) Canarium erythrinum
2 Papua New Guinea
5 Philippines
4 Philippines
1 Indonesia
3 Japan
(92) Canarium labiatum
1 Vanuatu
(87) Canarium manintveldi
1 NT, Australia
(88) Canarium darwinense
1 Japan
2 Japan
7 China
9 Papua New Guinea
6 China
8 China
4 China
3 China
5 China
(89) Canarium incisum
0.06
