## Supplementary Material 7 for "Molecular phylogenetics of the superfamily Stromboidea (Caenogastropoda): new insights from increased taxon sampling"

**Supplementary Material 7.** Bayesian analysis of COI, 12S, 16S, and 28S sequences, implemented in *BEAST. Posterior probability (PP) values are on branches; PP = 1 are replaced with red circles at nodes. Xenophoridae + Aporrhaidae + Struthiolariidae is resolved (red box), but not Rostellariidae + Seraphsidae + Strombidae.

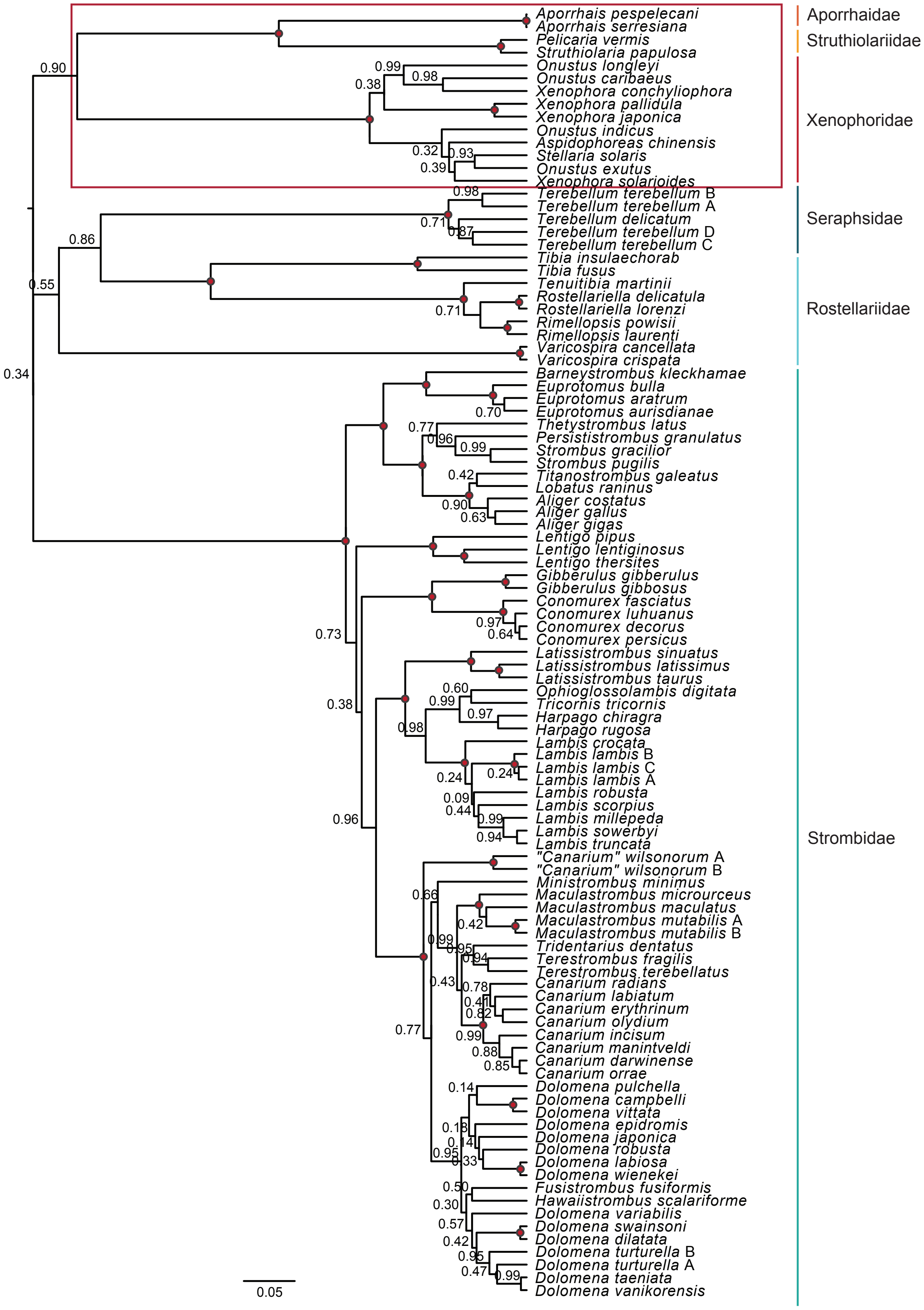

Aporrhais pespelecani
Aporrhais serresiana
Pelicaria vermis
Struthiolaria papulosa
Onustus longleyi
Onustus caribaeus
Xenophora conchyliophora
Xenophora pallidula
Xenophora japonica
Onustus indicus
Aspidophoreas chinensis
Stellaria solaris
Onustus exutus
Xenophora solarioides
Terebellum terebellum B
Terebellum terebellum A
Terebellum delicatum
Terebellum terebellum D
Terebellum terebellum C
Tibia insulaechorab
Tibia fusus
Tenuitibia martinii
Rostellariella delicatula
Rostellariella lorenzi
Rimellopsis powisii
Rimellopsis laurenti
Varicospira cancellata
Varicospira crispata
Barneystrombus kleckhamae
Euprotomus bulla
Euprotomus aratrum
Euprotomus aurisdianae
Thetystrombus latus
Persististrombus granulatus
Strombus gracilior
Strombus pugilis
Titanostrombus galeatus
Lobatus raninus
Aliger costatus
Aliger gallus
Aliger gigas
Lentigo pipus
Lentigo lentiginosus
Lentigo thersites
Gibberulus gibberulus
Gibberulus gibbosus
Conomurex fasciatus
Conomurex luhuanus
Conomurex decorus
Conomurex persicus
Latissistrombus sinuatus
Latissistrombus latissimus
Latissistrombus taurus
Ophioglossolambis digitata
Tricornis tricornis
Harpago chiragra
Harpago rugosa
Lambis crocata
Lambis lambis B
Lambis lambis C
Lambis lambis A
Lambis robusta
Lambis scorpius
Lambis millepeda
Lambis sowerbyi
Lambis truncata
"Canarium" wilsonorum A
"Canarium" wilsonorum B
Ministrombus minimus
Maculastrombus microurceus
Maculastrombus maculatus
Maculastrombus mutabilis A
Maculastrombus mutabilis B
Tridentarius dentatus
Terestrombus fragilis
Terestrombus terebellatus
Canarium radians
Canarium labiatum
Canarium erythrinum
Canarium olydium
Canarium incisum
Canarium manintveldi
Canarium darwinense
Canarium orrae
Dolomena pulchella
Dolomena campbelli
Dolomena vittata
Dolomena epidromis
Dolomena japonica
Dolomena robusta
Dolomena labiosa
Dolomena wienekei
Fusistrombus fusiformis
Hawaiistrombus scalariforme
Dolomena variabilis
Dolomena swainsoni
Dolomena dilatata
Dolomena turturella B
Dolomena turturella A
Dolomena taeniata
Dolomena vanikorensis
Aporrhaidae
Struthiolariidae
Xenophoridae
Seraphsidae
Rostellariidae
Strombidae
0.05
