## Supplementary Material 8 for "Molecular phylogenetics of the superfamily Stromboidea (Caenogastropoda): new insights from increased taxon sampling"

**Supplementary Material 8.** Bayesian analysis of independent genes 28S rRNA, 12S rRNA, and 16S rRNA, implemented in MrBayes. Putative species names are listed on the right, with corresponding number in parentheses (see Fig. 2; Suppl. Mat. 4). Support values are posterior probabilities (PP); branches with PP < 50% were collapsed, and intraspecific support values were removed for legibility except where relevant (i.e., if there was conflict between delimited ESUs or prior species assignations).

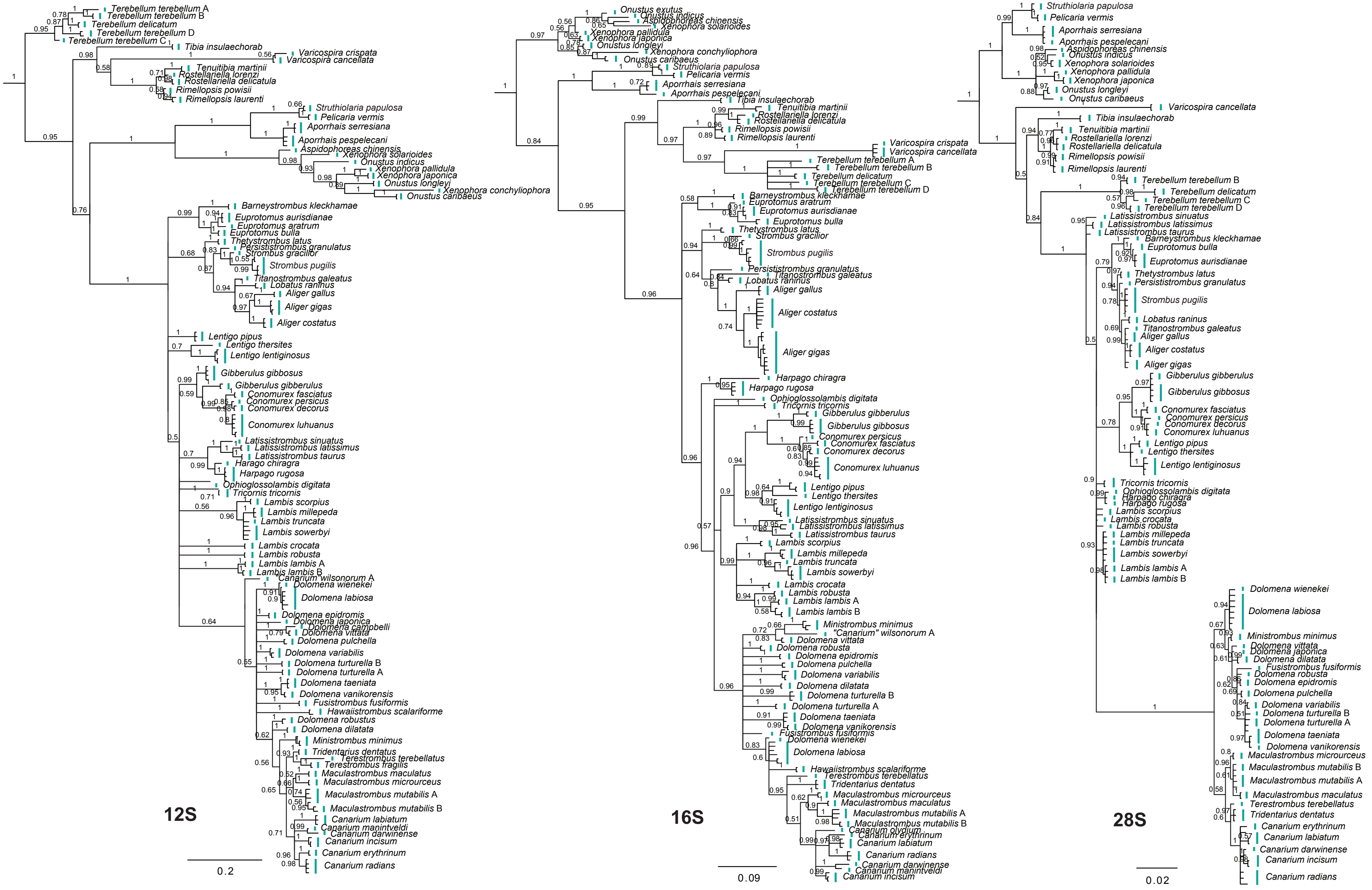
12S
0.2
Terebellum terebellum A
Terebellum terebellum B
Terebellum delicatum
Terebellum terebellum D
Terebellum terebellum C
Tibia insulaechorab
Varicospira crispata
Varicospira cancellata
Tenuitibia martinii
Rostellariella lorenzi
Rostellariella delicatula
Rimellopsis powisii
Rimellopsis laurenti
Struthiolaria papulosa
Pelicaria vermis
Aporrhais serresiana
Aporrhais pespelecani
Aspidophoreas chinensis
Xenophora solarioides
Onustus indicus
Xenophora pallidula
Xenophora japonica
Onustus longleyi
Xenophora conchyliophora
Onustus caribaeus
Barneystrombus kleckhamae
Euprotomus aurisdianae
Euprotomus aratrum
Euprotomus bulla
Thetystrombus latus
Persististrombus granulatus
Strombus gracilior
Strombus pugilis
Titanostrombus galeatus
Lobatus raninus
Aliger gallus
Aliger gigas
Aliger costatus
Lentigo pipus
Lentigo thersites
Lentigo lentiginosus
Gibberulus gibbosus
Gibberulus gibberulus
Conomurex fasciatus
Conomurex persicus
Conomurex decorus
Conomurex luhuanus
Latissistrombus sinuatus
Latissistrombus latissimus
Latissistrombus taurus
Harago chiragra
Harpago rugosa
Ophioglossolambis digitata
Tricornis tricornis
Lambis scorpius
Lambis millepeda
Lambis truncata
Lambis sowerbyi
Lambis crocata
Lambis robusta
Lambis lambis A
Lambis lambis B
Canarium wilsonorum A
Dolomena wienekei
Dolomena labiosa
Dolomena epidromis
Dolomena japonica
Dolomena campbelli
Dolomena vittata
Dolomena pulchella
Dolomena variabilis
Dolomena turturella B
Dolomena turturella A
Dolomena taeniata
Dolomena vanikorensis
Fusistrombus fusiformis
Hawaiistrombus scalariforme
Dolomena robustus
Dolomena dilatata
Ministrombus minimus
Tridentarius dentatus
Terestrombus terebellatus
Terestrombus fragilis
Maculastrombus maculatus
Maculastrombus microurceus
Maculastrombus mutabilis A
Maculastrombus mutabilis B
Canarium labiatum
Canarium maninveldi
Canarium darwinense
Canarium incisum
Canarium erythrinum
Canarium radians
16S
0.09
Onustus exutus
Onustus indicus
Aspidophoreas chinensis
Xenophora solarioides
Xenophora pallidula
Xenophora japonica
Onustus longleyi
Xenophora conchyliophora
Onustus caribaeus
Struthiolaria papulosa
Pelicaria vermis
Aporrhais serresiana
Aporrhais pespelecani
Tibia insulaechorab
Tenuitibia martinii
Rostellariella lorenzi
Rostellariella delicatula
Rimellopsis powisii
Rimellopsis laurenti
Varicospira crispata
Varicospira cancellata
Terebellum terebellum A
Terebellum terebellum B
Terebellum delicatum
Terebellum terebellum C
Terebellum terebellum D
Barneystrombus kleckhamae
Euprotomus aratrum
Euprotomus aurisdianae
Euprotomus bulla
Thetystrombus latus
Strombus gracilior
Strombus pugilis
Persististrombus granulatus
Titanostrombus galeatus
Lobatus raninus
Aliger gallus
Aliger costatus
Aliger gigas
Harpago chiragra
Harpago rugosa
Ophioglossolambis digitata
Tricornis tricornis
Gibberulus gibberulus
Gibberulus gibbosus
Conomurex persicus
Conomurex fasciatus
Conomurex decorus
Conomurex luhuanus
Lentigo pipus
Lentigo thersites
Lentigo lentiginosus
Latissistrombus sinuatus
Latissistrombus latissimus
Latissistrombus taurus
Lambis scorpius
Lambis millepeda
Lambis truncata
Lambis sowerbyi
Lambis crocata
Lambis robusta
Lambis lambis A
Lambis lambis B
Ministrombus minimus
"Canarium" wilsonorum A
Dolomena vittata
Dolomena robusta
Dolomena epidromis
Dolomena pulchella
Dolomena variabilis
Dolomena dilatata
Dolomena turturella B
Dolomena turturella A
Dolomena taeniata
Dolomena vanikorensis
Fusistrombus fusiformis
Dolomena wienekei
Dolomena labiosa
Hawaiistrombus scalariforme
Terestrombus terebellatus
Tridentarius dentatus
Maculastrombus microurceus
Maculastrombus maculatus
Maculastrombus mutabilis A
Maculastrombus mutabilis B
Canarium olydium
Canarium erythrinum
Canarium labiatum
Canarium radians
Canarium darwinense
Canarium maninveldi
Canarium incisum
28S
0.02
Struthiolaria papulosa
Pelicaria vermis
Aporrhais serresiana
Aporrhais pespelecani
Aspidophoreas chinensis
Onustus indicus
Xenophora solarioides
Xenophora pallidula
Xenophora japonica
Onustus longleyi
Onustus caribaeus
Varicospira cancellata
Tibia insulaechorab
Tenuitibia martinii
Rostellariella lorenzi
Rostellariella delicatula
Rimellopsis powisii
Rimellopsis laurenti
Terebellum terebellum B
Terebellum delicatum
Terebellum terebellum C
Terebellum terebellum D
Latissistrombus sinuatus
Latissistrombus latissimus
Latissistrombus taurus
Barneystrombus kleckhamae
Euprotomus bulla
Euprotomus aurisdianae
Thetystrombus latus
Persististrombus granulatus
Strombus pugilis
Lobatus raninus
Titanostrombus galeatus
Aliger gallus
Aliger costatus
Aliger gigas
Gibberulus gibberulus
Gibberulus gibbosus
Conomurex fasciatus
Conomurex persicus
Conomurex decorus
Conomurex luhuanus
Lentigo pipus
Lentigo thersites
Lentigo lentiginosus
Tricornis tricornis
Ophioglossolambis digitata
Harpago chiragra
Harpago rugosa
Lambis scorpius
Lambis crocata
Lambis robusta
Lambis millepeda
Lambis truncata
Lambis sowerbyi
Lambis lambis A
Lambis lambis B
Dolomena wienekei
Dolomena labiosa
Ministrombus minimus
Dolomena vittata
Dolomena japonica
Dolomena dilatata
Fusistrombus fusiformis
Dolomena robusta
Dolomena epidromis
Dolomena pulchella
Dolomena variabilis
Dolomena turturella B
Dolomena turturella A
Dolomena taeniata
Dolomena vanikorensis
Maculastrombus microurceus
Maculastrombus mutabilis B
Maculastrombus mutabilis A
Maculastrombus maculatus
Terestrombus terebellatus
Tridentarius dentatus
Canarium erythrinum
Canarium labiatum
Canarium darwinense
Canarium incisum
Canarium radians
